# Complete post-transcriptional modification profiles in individual *Staphylococcus aureus* tRNA species

**DOI:** 10.1101/2025.10.30.685614

**Authors:** Jose R. Jaramillo-Ponce, Philippe Wolff, Virginie Marchand, Yuri Motorin, Maximilian Kohl, Hiroki Kanazawa, Aila Ruiz-Paterson, Béatrice Chane-Woon-Ming, Caroline Paulus, Johana Chicher, Anne-Sophie Gribling-Burrer, Redmond Smyth, Martina Krämer, Mark Helm, Pascale Romby, Stefano Marzi

**Author notes:** To whom correspondence should be addressed. Stefano Marzi. Jose R. Jaramillo-Ponce.

## Abstract

Transfer RNAs play a critical role in protein synthesis by matching mRNA codons to their corresponding amino acids. Their post-transcriptional modifications, shaping structure, stability, and codon decoding, are now recognized as key regulators of translation and cellular adaptation, including in bacterial pathogens. Here, we provide a comprehensive analysis of tRNA modifications in *Staphylococcus aureus* using extensive oligonucleotide mass spectrometry and deep-sequencing methods, generating a high-confidence modification map for each individual tRNA species, including the non-proteogenic tRNA^Gly^. While the overall tRNA modification landscape is conserved among Gram-positive bacteria, our data uncovered unexpected *S. aureus*-specific features. These include the absence of [m^2^A]37 in tRNAs despite the presence of the methyltransferase RlmN, a single multi-site DusB2 enzyme catalyzing all tRNA dihydrouridylation, and a dedicated pseudouridine synthase responsible for [Ψ]32 formation. Besides, heterogenous modification patterns were observed in tRNA^Leu^(UAA) and tRNA^Lys^(CAA), highlighting a complex interplay in anticodon hypermodification. Integration of ribosome profiling and Nanopore tRNA sequencing offered a global view of the decoding properties of the reduced *S. aureus* tRNA set, revealing efficient four-way wobble recognition, slower translation of rare codons by low abundant tRNAs, and distinctive decoding dynamics of Gly codons potentially influenced by the unusual modification status of tRNA^Gly^(UCC). This work establishes a framework for future research aimed at dissecting the role of specific tRNA modifications in *S. aureus* physiology and pathogenesis.

## INTRODUCTION

Transfer RNA (tRNAs) are key components of the translation apparatus, ensuring accurate decoding of mRNA codons and delivery of the correct amino acids to the ribosome (1). The balance between the abundance of different tRNA species and the codons they decode influences translation speed and fidelity (2, 3), thereby affecting protein synthesis rates, co-translational folding, and protein localization (4). Among the factors modulating this balance, the modified nucleotides present in tRNA molecules play a crucial role by shaping their structural properties and decoding capacities (5). These post-transcriptional modifications can be regulated in response to environmental signals (6), reprogramming translation through mechanisms such as codon-biased decoding (7), changes in tRNA aminoacylation efficiency (8), and effects in tRNA stability and maturation (9, 10), ultimately promoting adaptation. Although still underexplored, this layer of regulation is likely central to bacterial physiology, particularly in pathogenic species, enabling a rapid reconfiguration of gene expression in response to fluctuating environments and host-imposed stresses (11).

Around 60 distinct natural modifications have been identified in bacterial tRNAs (12), which contain on average eight modified nucleotides (13). These modifications are predominantly located in the anticodon loop and in the tRNA core region formed by the interaction of the D- and TΨC loops (14). Within the anticodon loop, positions 34 and 37 are the most frequently modified, exhibiting remarkable chemical diversity among different tRNA species (15). Modifications at position 34 influence wobble base pairing, which enables a single tRNA to recognize multiple synonymous codons (16), while those at position 37, immediately adjacent to the anticodon, modulate the stability of the codon-anticodon interaction and contribute to reading frame maintenance (17). In contrast, modifications in the tRNA core are less diverse and primarily affect structure dynamics and stability of tRNAs (18, 19), with pseudouridine (Ψ), dihydrouridine (D), and 5-methyluridine (m^5^U) being the most common and widespread across tRNA species.

tRNA modifications are introduced by a diverse array of enzymes with distinct substrate specificities and catalytic mechanisms (20). Many modifications, especially those in the tRNA core region, are installed by stand-alone enzymes, whereas modifications in the anticodon loop often arise through the sequential action of multiple enzymes (21). Most enzymes modify specific positions across multiple tRNA species by recognizing structural or sequence features conserved within their substrates (22). The tRNA-dihydrouridine synthase (Dus) families exemplify the diversity of substrate recognition, as distinct species-specific enzymes install the same modification at different tRNA positions (23). In *Escherichia coli*, DusA introduces [D]20 and [D]20a, while DusB and DusC modify specifically positions 17 and 16, respectively (24). In contrast, *Bacillus subtilis* possesses two types of DusB enzymes with overlapping but distinct preferential activities: DusB1 is responsible for all [D]17, [D]20, [D]20a, and [D]47 modifications, while DusB2 is more specific to positions 20 and 20a (25). Notably, some enzymes exhibit modification activities beyond tRNA, such as *E. coli* RlmN, which introduces 2-methyladenosine (m^2^A) in tRNA and 23S rRNA (26).

Despite their recognized importance in translation, comprehensive information on tRNA modifications and their corresponding enzymes is available for a limited number of organisms, including the model bacteria *E. coli* (21) and *B. subtilis* (27), as well as several pathogenic species such as *Vibrio cholerae* (28, 29), *Mycobacterium tuberculosis* (30), *Streptomyces albidoflavus* (31), *Bartonella henselae/quintana* (32)*, Pseudomonas aeruginosa* (33), and *Streptococci* (34). While these studies showed that many tRNA modifications and their biosynthetic enzymes are broadly conserved across bacteria, they revealed species-specific modification patterns, underscoring the need for more systematic investigations in additional bacterial species across phylogeny. However, these studies remained rather global, combining approaches such as nucleoside mass spectrometry, deep-sequencing, genetic validations, and bioinformatic predictions. While informative, they do not provide high-confidence modification maps for every individual tRNA, leaving the full diversity and heterogeneity of modification profiles uncompleted. To date, *Mycoplasma capricolum* remains the only organism in which the complete set of native tRNA sequences has been systematically determined (35).

We previously reported a partial map of tRNA modifications in *Staphylococcus aureus*, a major human pathogen responsible for a wide range of diseases (36). The genome of this Gram-positive bacterium encodes 30 distinct tRNA species decoding all 61 sense codons, with the number of gene copies for each isoacceptor varying between strains (37). In addition, *S. aureus* harbors three non-proteogenic tRNA^Gly^ species (38, 39) that are not used for protein synthesis (40). Instead, they serve as substrates for the aminoacyl-tRNA transferases FemX, FemA, and FemB, which catalyze the formation of pentaglycine bridges in the peptidoglycan cell wall (41). In this work, we combined oligonucleotide mass spectrometry with multiple deep sequencing methods to generate complete, high-confidence maps of post-transcriptional modifications in each individual *S. aureus* tRNA species, including the non-proteogenic tRNAs. This enabled assignment of the corresponding tRNA-modifying enzymes by gene orthology, the analysis of their expression profiles, and the identification of several unexpected modification patterns in *S. aureus*. By integrating ribosome profiling and Nanopore tRNA abundance measurements, we also provide a global picture of the decoding properties of *S. aureus* tRNAs and dynamics of codon-decoding.

## RESULTS

### Integrating mass spectrometry and deep sequencing for tRNA modification profiling

We previously developed a method for identifying tRNA modifications within their sequence context using ultra-performance liquid chromatography coupled with tandem mass spectrometry (UPLC-MS/MS) (42). In this strategy, bulk tRNAs are fractionated by two-dimensional polyacrylamide gel electrophoresis (2D-PAGE), and the excised spots are digested with a specific RNase to generate oligonucleotides for UPLC-MS/MS analysis. Application of this method to *S. aureus*, analyzing RNase T1 (G-specific) and RNase A (C/U-specific) digests, enabled partial mapping of modifications in 24 of the 30 proteogenic tRNA species, with an average sequence coverage of 60% (36). However, this strategy did not allow detection of Ψ as this abundant modification is isobaric with U. To overcome these limitations and achieve a complete map of modifications in all *S. aureus* tRNA species, we refined the method by incorporating additional steps (Figure 1A, Figure S1). Bulk tRNAs were first separated by reversed-phase chromatography to enrich low-abundance species, and the resulting fractions were treated with acrylonitrile prior to 2D-PAGE to generate stable cyanoethylated (ce) Ψ sites (43, 44). Interestingly, 4-thiouridine (s^4^U) was also labeled, but only when the reaction was performed before electrophoresis (Figure 1B). These data suggesting that s^4^U had been artificially lost in our previous study, most likely due to gel UV exposure (45). Other modifications were also cyanoethylated, including lysidine (k^2^C) (Figure 1E), inosine (I), 5-methoxyuridine (mo^5^U), queuosine (Q), 5-methylaminomethyluridine (mnm^5^U), and the 2-thiolated mnm^5^s^2^U (Supplementary Data 1). Near full-sequence coverage was reach by including digestion with RNase U2 (purine specific) (46), which mainly cleaved after A, leaving 2’, 3’ cyclic phosphate termini (>p) and yielding a mixture of shorter and longer fragments due to frequent missed cleavages. This facilitated analysis of G- and U-rich tRNA regions poorly resolved by RNase T1 and RNase A, such as the 5’ termini (Figure 1B), the D loop (Figure 1C), and the TΨC stem (Figure 1G). UPLC-MS/MS results are compiled in Supplementary Data 1, which provides, for each tRNA, a representative set of deconvoluted MS/MS spectra showing sequence coverage and detected modifications.

**Figure 1.**
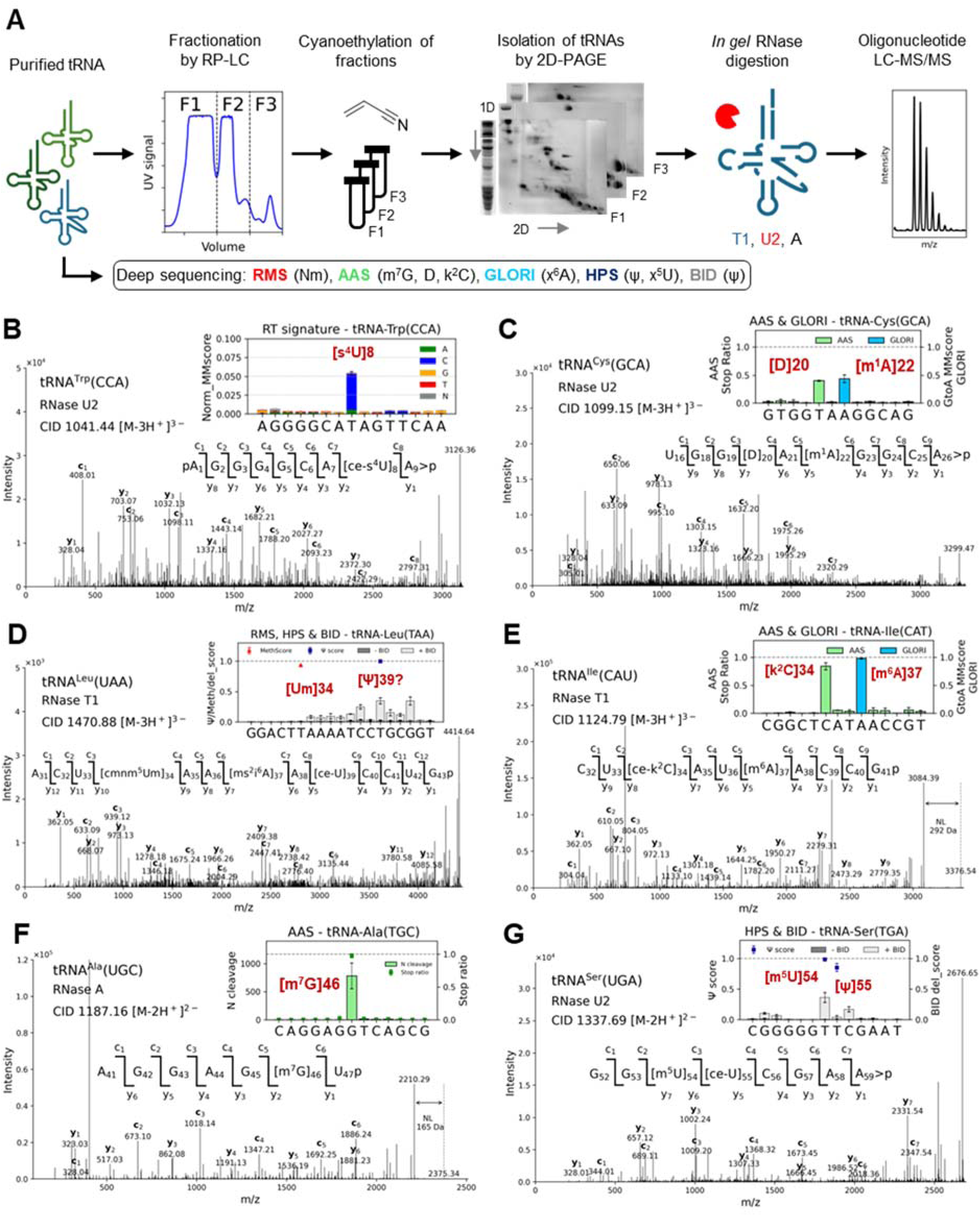
Integrative approach for detecting tRNA modifications in *S. aureus*. (**A**) Workflow for UPLC-MS/MS analysis of individual tRNAs. Separation of bulk tRNA by reversed-phase liquid chromatography (RP-LC) and 2D-PAGE of each fraction enabled isolation of all tRNA species. Cyanoethylation prior to electrophoresis revealed potential Ψ sites and allowed detection of s^4^U. Digestion with RNases T1, A, and U2 provided near-complete sequence coverage. Several deep-sequencing methods were applied to bulk tRNAs: RMS (RiboMethSeq), AAS (AlkAnilineSeq), GLORI (glyoxal/nitrite A deamination), HPS (HydraPsiSeq), BID (bisulfite-induced deletion). (**B to G**) Deconvoluted CID-MS/MS spectra of tRNA fragments containing modified nucleosides. The corresponding tRNA, RNase employed, and precursor ion (mass and charge) are indicated. Relevant *y*- and *c*-product ions are labeled. Insets show deep-sequencing signals. (**B**) [ce-s^4^U]8 in tRNA^Trp^(CCA), also detected by RT-signature. (**C**) [D]20 and [m^1^A]22 in tRNA^Cys^(GCA), confirmed by AAS and GLORI, respectively. (**D**) Modifications in the anticodon loop of tRNA^Leu^(UAA). RMS detected [Um]34, and Ψ (likely 39) was supported by HPS and BID. (**E**) [k^2^C]34 and [m^6^A]37 in tRNA^Ile^(CAU), also detected by AAS and GLORI, respectively. (**F**) [m^7^G]46 in tRNA^Ala^(UGC), validated by AAS. (**G**) [m^5^U]54 and [Ψ]55 in tRNA^Ser^(UGA). HPS detected both, while BID misassigned Ψ to -1 due to consecutive T54 and T55.

Our UPLC-MS/MS mapping was complemented with several deep-sequencing approaches (Figure 1A), enabling detection of specific modifications across all tRNA sequences (Supplementary Data 2). This was particularly important for methylations, as oligonucleotide mass spectrometry is unable to determine the exact location of methyl groups within the nucleotide. RiboMethSeq (47) confirmed 2’-O-methylations (Nm) (Figure 1D) and the extracted reverse transcriptase (RT) misincorporation signatures corroborated s^4^U sites (Figure 1B). AlkAnilineSeq (48, 49) readily detected 7-methylguanosine (m^7^G) and supported UPLC-MS/MS assignments, which were characterized by a characteristic 165 Da neutral loss (Figure 1F). It also contributed to detect several D positions (Figure 1C) and k^2^C, the latter likewise associated with a typical neutral loss in UPLC-MS/MS (Figure 1E). GLORIseq (50) detected several N^6^-modified adenosines, such as N^6^-methyladenosine (m^6^A) (Figure 1E), N^6^-isopentenyladenosine (i^6^A), and N^6^-threonylcarbamoyladenosine (t^6^A). It also detected 1-methyladenosine (m^1^A), with a signal arising from sub-stoichiometric Dimroth rearrangement into m^6^A (51), as evidenced by systematic lower scores (Figure 1C). For assignment of Ψ sites, several distinct sequencing protocols were combined, as cyanoethylation in our conditions produced several false positives (Supplementary Data 1). HydraPsiSeq (52) detected Ψ and 5-modified uridines, such as m^5^U, mnm^5^U, and mo^5^U, based on their resistance to the hydrazine cleavage, quantified by a Ψ-score (53). Furthermore, we applied BIDseq (54), which uses bisulfite to generate an adduct producing a deletion signature at Ψ sites (Figure 1D and 1G). Finally, Nanopore direct RNA sequencing, primarily used here to quantify tRNA abundance, also provided additional information about modified positions, including Ψ sites (Figure S2).

### Repertoire of modifications and enzymes in proteogenic tRNAs

Our integrative analysis provided a high-confidence map of tRNA modifications in *S. aureus* under rich medium conditions, covering all 30 proteogenic tRNA species (Table 1, Figure 2A, Figure S3). Here, isodecoders tRNA^His^(GUG) (2 genes), tRNA^Leu^(UAA) (2 genes), tRNA^Lys^(UUU) (3 genes), and tRNA^Ser^(UGA) (3 genes), differing in only one or two positions, were treated as single tRNA species. With the repertoire of modifications established, the corresponding enzymes were assigned based on literature data and FoldSeek-based ortholog predictions (55) (Figure 2B). The genomic context of the identified enzymes in the HG001 strain is shown in Figure S4.

**Figure 2.**
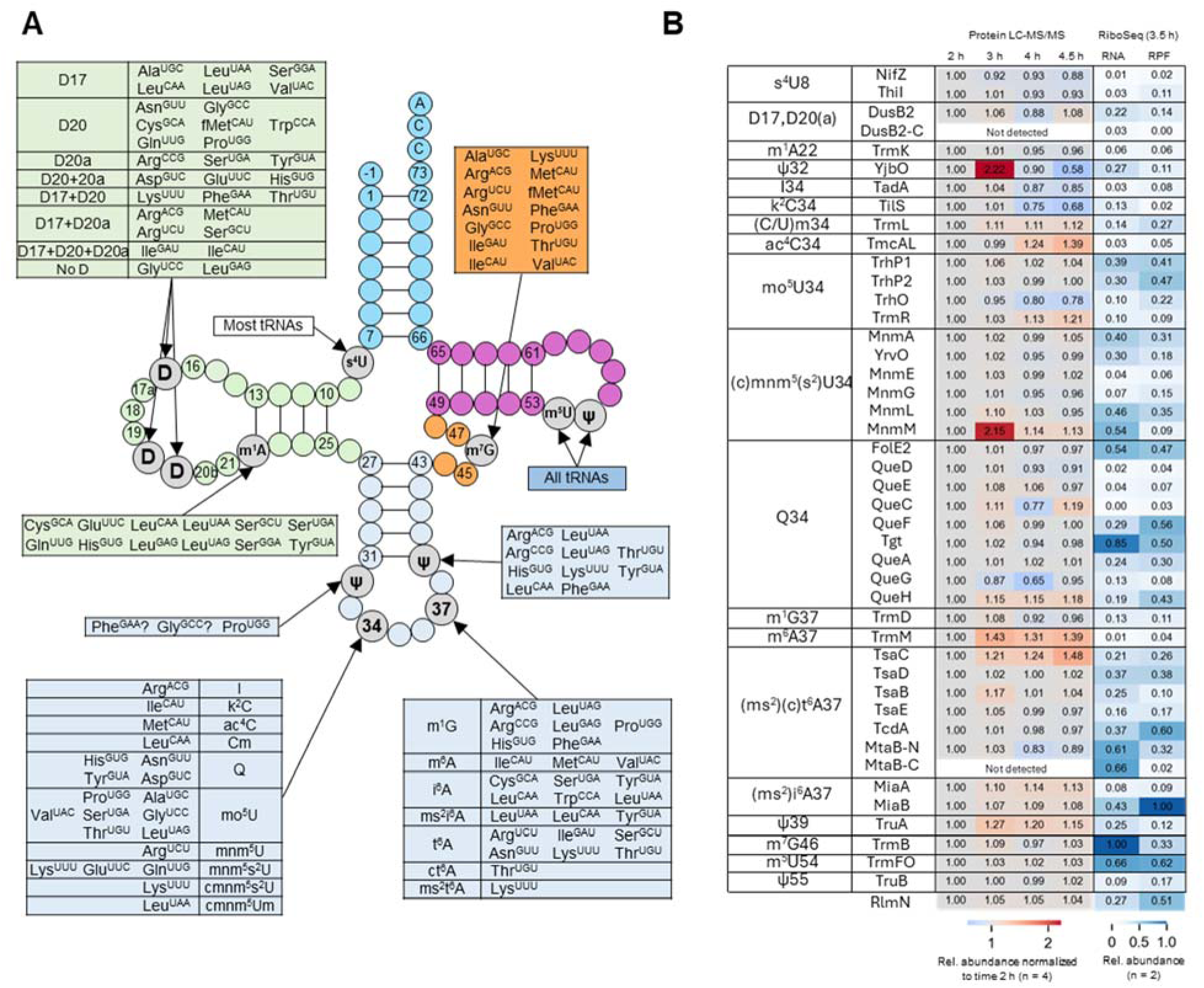
Repertoire of tRNA modifications and their enzymes in *S. aureus*. (**A**) Summary of identified modifications and their positions within tRNA sequence. The species carrying each modification are indicated. For D, tRNAs are grouped according to their modification patterns at positions 17, 20 and 20a. (**B**) Assignment of tRNA-modifying enzymes by gene orthology and analysis of their expression profiles. The levels of each enzyme were monitored during exponential growth by time-course protein LC-MS/MS (left heatmap). Protein abundances across conditions are normalized to the first time point, with values higher or lower than 1 indicating increasing or decreasing expression, respectively. The relative abundance of the tRNA-modifying enzymes at the transcriptome (RNA) and translatome (RPF) level was estimated from total RNA sequencing and ribosome profiling performed at late exponential phase (right heatmap). TPM (transcript per million) was used to compare the abundance of each enzyme within samples, and were normalized based on the highest (1) and lowest (0) values.

**Table 1.**
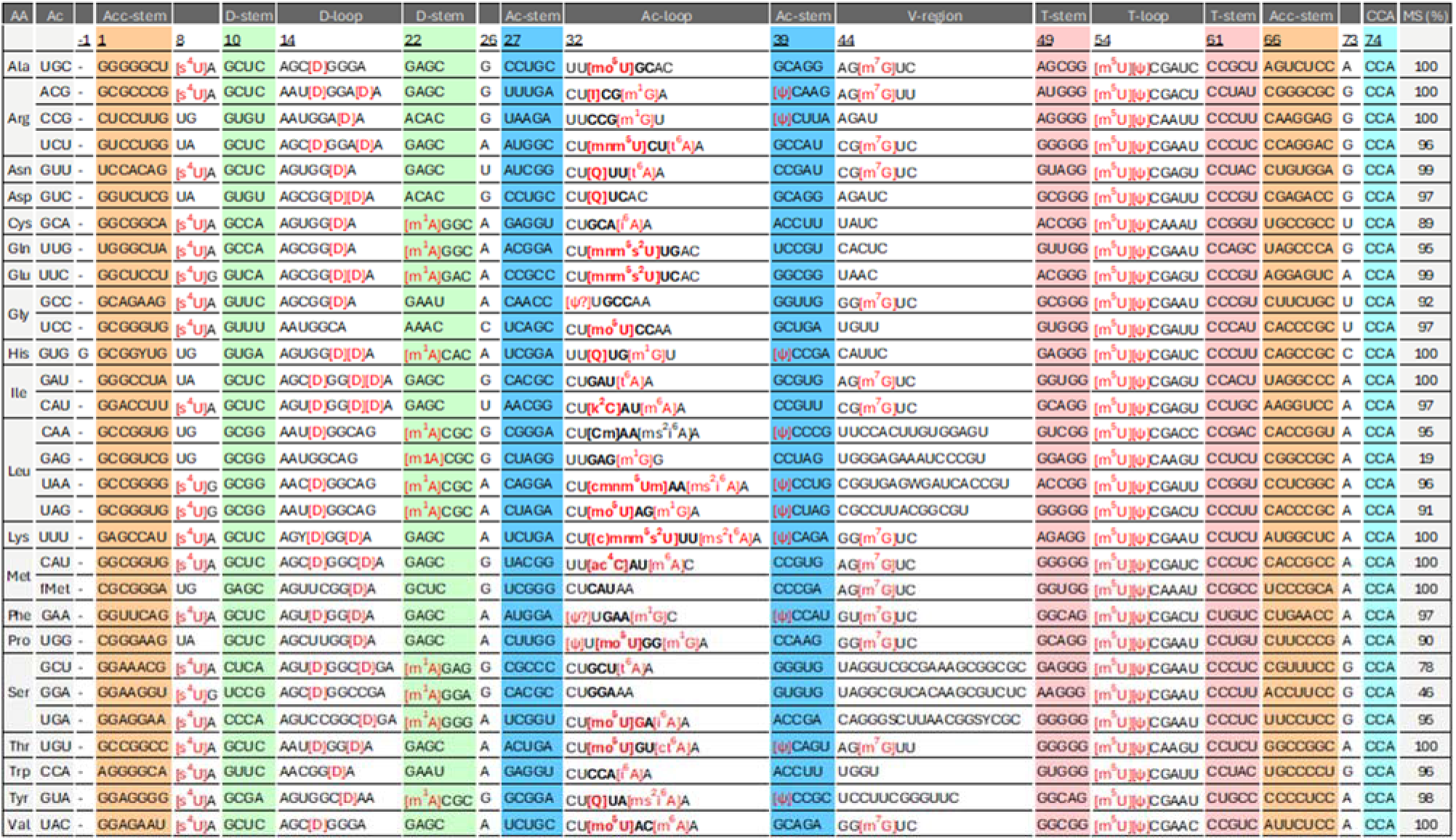
Native sequences of *S. aureus* tRNAs. The 30 proteogenic tRNA species are grouped by cognate amino acid (AA). Sequences are aligned according to the cloverleaf secondary structure, using standard tRNA numbering (62). Regions forming the acceptor (Acc), anticodon (Ac) and TΨC (T) stems are colored. The CCA 3’-end is highlighted in light blue. Modified nucleotides are shown in brackets and highlighted in red. Anticodons are indicated in bold. Sequence coverage by mass spectrometry is indicated in the last column. Single-nucleotide polymorphisms are denoted using conventional nucleotide codes: Y (C or U), S: (G or C), and W (A or U).

Six modified nucleotides were identified within the tRNA core region: s^4^U, D, m^5^U, and Ψ present in most tRNAs, while m^1^A and m^7^G appear in distinct specific subsets (Figure 2A). All proteogenic tRNAs contained the two highly conserved [m^5^U]54 and [Ψ]55 in the TΨC-loop (56), installed by TrmFO (57) and TruB (58), respectively. In the D-loop of nearly all tRNAs, D was found at positions 17, 20 and 20a, except for tRNA^Gly^(UCC) and tRNA^Leu^(GAG) lacking U at these sites. In the acceptor-stem/D-stem junction, [s^4^U]8 was detected in 24 tRNAs, while the remaining 6 contained an unmodified U. In the D-arm, [m^1^A]22 was found in 12 tRNAs spanning the seven isoacceptor families reported in previous studies (59), all of which carry a purine at position 13, a strict requirement for TrmK activity (60). Consistent with this, tRNA^Arg^(CCG), tRNA^Asp^(GUC), and tRNA^Gly^(UCC) containing U13:A22 base pairs lacked this modification. In the variable region, 14 tRNAs carried [m^7^G]46 (substrates of TrmB) within a 5-nucleotide loop, where only G46 and U47 were strictly conserved. All of these species contain a C13:G22 base pair known to interact with [m^7^G]46, stabilizing the tRNA structure (61). In contrast, all other non-substrates of TrmB either lacked G46 or exhibit shorter or longer variable regions.

A greater diversity of modifications was identified in the anticodon loop, specifically at positions 34 and 37 (Figure 2A). Only two tRNAs lacked any modification in this region: initiator tRNA^fMet^(CAU) and tRNA^Ser^(GGA). Modifications at position 34 were found in 20 tRNAs and could occur on any nucleoside. The A34- and C34-modifications were restricted to specific tRNAs, each catalyzed by well-conserved stand-alone enzymes: [I]34 in Arg(ACG) (TadA) (63), [k^2^C]34 in Ile(CAU) (TilS) (64), [Cm]34 in Leu(CAA) (TrmL) (65), and N^4^-acetylcytidine 34 (ac^4^C) in Met(CAU) (TmcAL) (66). No other tRNA contained unmodified A34, while C34 remained unmodified in tRNA^Arg^(CCG) and tRNA^Trp^(CCA). The only modified G34 was queuosine (Q), present in the four tRNAs – Asn(GUU), Asp(GUC), His(GUG), and Tyr(GUA) –, while it remained unmodified in all other G34-containing tRNAs. In contrast, U34 was always modified, displaying either mo^5^U or xm^5^U derivatives. In addition to the six species reported to contain [mo^5^U]34 in *B. subtilis* (67), tRNA^Gly^(UCC) in *S. aureus* also harbored this modification instead of the usual mnm^5^U (56). Among the xm^5^U modifications, we detected mnm^5^U in tRNA^Arg^(UCU); mnm^5^s^2^U in tRNAs – Glu(UUC), Gln(UUG), and Lys(UUU) –; cmnm^5^s^2^U also in tRNA^Lys^(UUU); and cmnm^5^Um in tRNA^Leu^(UAA), where TrmL introduces the 2’-O-ribomethylation. The synthesis of [Q]34, [mo^5^U]34 and [xm^5^U]34 modifications requires multiple enzymes (Figure 2B). The order of the enzymatic reactions within these complex pathways are shown in Figure S5.

Modifications at position 37 are all purine-derived. All seven tRNAs with G37 carried 1-methylguanosine (m^1^G), also containing G36, which is required for efficient modification by TrmD (68). In contrast, A37 could remain unmodified or be modified to m^6^A, i^6^A or t^6^A (Table 1). The [m^6^A]37 modification was restricted to three tRNAs – Ile(CAU), Met(CAU), and Val(UAC) – and a TrmM ortholog (69) was identified in *S. aureus*. In contrast, i^6^A and t^6^A were more broadly distributed, each present in six different tRNAs. In addition to the conserved TsaCBDE enzymes responsible for [t^6^A]37 synthesis, we identified a TcdA ortholog known to catalyze formation of the cyclic hydantoin form of t^6^A (ct^6^A) (70) (Figure S5). This modification was only validated in tRNA^Thr^(UGU) (Supplementary Data 1), but its presence cannot be excluded in other tRNAs as this modification is labile (71). The methylthiolated derivatives of i^6^A and t^6^A were also detected: 2-methylthio-N^6^-isopentenyladenosine (ms^2^i^6^A) in tRNAs – Leu(UAA), Leu(CAA) and Tyr(GUA) –, and 2-methylthio-N^6^-threonylcarbamoyladenosine (ms^2^t^6^A) in tRNA^Lys^(UUU) when purified from the *S. aureus* USA300 JE2 strain. In the remaining eight tRNAs, A37 was unmodified.

Finally, two Ψ sites were identified in the anticodon stem-loop: [Ψ]32 in a few tRNAs, and [Ψ]39 in all tRNAs carrying U39, except for tRNA^Gln^(UUG). However, detection of Ψ at these positions was often challenging (Supplementary Data 2). In tRNA^Gly^(GCC), U32 showed significant HPS Ψ-score (ScoreMean > 0.92 and Score A2 > 0.5) but was not detected by BIDseq, whereas in tRNA^Pro^(UGG), U32 produced a clear BID signal, but its Ψ-score did not meet the detection thresholds. Both HPS and BID signals for U32 in tRNA^Phe^(GAA) were only moderately convincing. The gene *yjbO*, responsible for [Ψ]31 and [Ψ]32 in *B. subtilis* (27), is also present in *S. aureus* (Figure 2B), but only tRNA^Pro^(UGG) is a common substrate, supporting the presence of [Ψ]32 in this tRNA. For tRNA^Arg^(CCG), assignment of [Ψ]39 was based on cyanoethylation and Nanopore C-to-U mismatch (Figure S2), as reliable HPS or BIDseq data could not be obtained, possibly due to the low abundance of this tRNA. Intriguingly, cyanoethylation in tRNA^Leu^(UAA) occurred only at U39, but higher HPS and BID signals were unexpectedly observed at U42 (Figure 1D), a position not known to be modified by TruA (72).

### Expression profiles of tRNA-modifying enzymes

The expression of tRNA modification enzymes in *S. aureus* HG001 was monitored during exponential growth in rich medium by label-free protein LC-MS/MS (Figure 2B, left heatmap). Growth curves, sampled time points and the intensity-based quantification data are provided in Supplementary Data 3. Additionally, the relative abundance of these enzymes in the transcriptome and translatome was estimated using normalized read counts from ribosome profiling and total RNA sequencing data obtained at late exponential growth phase (Figure 2B, right heatmap).

With the exception of MtaB and one of the two DusB2 proteins, all tRNA-modifying enzymes were detected in the time-course proteomics experiment, revealing distinct expression profiles. Many enzymes showed highest levels during early exponential growth, which then declined over time. This was the case for ThiI and NifZ, both required for [s^4^U]8 synthesis (73), consistent with findings in *E. coli* showing that s^4^U levels decrease with growth rate (74). In contrast, other enzymes started to accumulate later, with their levels either remaining stable or gradually increasing over time. Among these, TmcAL and TrmM showed a progressive increase in expression, both targeting the anticodon loop of the elongator tRNA^Met^(CAU). Likewise, the [ms^2^i^6^A]37 synthesis enzymes MiaA and MiaB showed similar positive trends, together with TrmL, in line with the requirement of [ms^2^i^6^A]37 for TrmL-dependent methylation of position 34 (65, 75). Distinct expression patterns were also observed among the enzymes involved in complex modification pathways (Figure S5). Synthesis of [mo^5^U]34 occurs through a two-step process, involving hydroxylation of U34 to 5-hydroxyuridine (ho^5^U) followed by subsequent methylation. The first step is catalyzed independently by two types of hydroxylases, TrhP1/P2 and the O_2_-dependent TrhO, which displayed distinct expression profiles. While TrhP1 and TrhP2 levels remained relatively stable throughout growth, TrhO expression decreased markedly over time (Figure 2B), likely reflecting reduced O_2_ availability at higher cell density. In contrast, TrmR, the enzyme catalyzing the final methylation step, appeared to progressively accumulate during growth. Similarly, among the enzymes involved in [Q]34 synthesis, the two reductases QueG (cobalamin-dependent) and QueH (Fe-dependent) (76) – which redundantly catalyze the final step of the pathway in *S. aureus* – displayed distinct expression profiles, possibly reflecting differences in the availability of their distinct cofactors, with QueH being more translated than QueG at late exponential phase.

Taken together, these data strongly suggest that the expression of the enzymes is adjusted according to the availability of their cofactors and is sensitive to the metabolism adaptation during growth.

### Divergence of dual-specific RNA modification activities

In *E. coli*, three rRNA-modifying enzymes also act on tRNA (Figure 3A). RluF is responsible for [Ψ]2604 in 23S rRNA and [Ψ]35 in tRNA Tyr(GUA) (77), RluA introduces [Ψ]746 in 23S rRNA and [Ψ]32 in four tRNA species (78), and RlmN synthesizes [m^2^A]2503 in 23S rRNA and [m^2^A]37 in six tRNAs (26). However, such dual-specific activities do not appear to be conserved in *S. aureus*. Consistent with the absence of an RluF ortholog and the lack of modification at the site equivalent to [Ψ]2604 in *B. subtilis* 23S rRNA (U2633) (79, 80), we did not detect any convincing [Ψ]35 in *S. aureus* tRNAs. Although [Ψ]32 was present in some tRNA species, the enzyme YjbO, experimentally linked to this modification in *B. subtilis* (27), does not appear to be an RluA ortholog, as it contains an N-terminal S4-like domain absent in *E. coli* RluA (81) (Figure 3B, left panel). Moreover, none of the potential YjbO substrates in *B. subtilis* or *S. aureus* contained the [Ψ]UXXAAA motif recognized by *E. coli* RluA (81) and the 23S rRNA position corresponding to [Ψ]746 in *B. subtilis* (U793) is not modified despite the presence of the motif (79, 80). Whether YjbO exhibits dual activities specific to *S. aureus* remains to be determined.

*S. aureus* harbors a clear RlmN ortholog, and m^2^A at position 2530 of 23S rRNA (equivalent to 2503 in *E. coli*) has been experimentally linked to this enzyme (82). However, our analysis of *S. aureus* tRNAs revealed no methylated A37, except for [m^6^A]37 in three tRNAs (Table 1). LC-MS/MS nucleoside quantification confirmed that m^2^A is essentially absent from total tRNA, whereas common modifications such as m^5^U, Ψ, D, and m^7^G were detected at levels consistent with their distribution across tRNAs (Figure 3C, Supplementary Data 4). Since RlmN is expressed, as indicated by proteomics and ribosome profiling data (Figure 2B), we asked whether the absence of m^2^A could be explained by the lack of tRNA features recognized by RlmN (Figure 3D). Among the *S. aureus* tRNAs carrying unmodified A37, tRNAs – Ala(UGC), Gly(GCC), Gly(UCC), fMet(CAU), and Ser(GGA) – are not RlmN substrates in *E. coli,* and all except tRNA^Ala^(UGC) possess A38, which is known to abolish the activity of the enzyme *in vitro* (83). In contrast, tRNAs – Gln(UUG), Glu(UUC), and Asp(GUC) –, which have anticodon stem-loop sequences nearly identical to their *E. coli* counterparts carrying [m^2^A]37, yet remain unmodified (Table 1, Figure S3). Hence, our data suggests that *S. aureus* RlmN activity may be restricted to 23S rRNA. We then compared *S. aureu*s RlmN with the *E. coli* dual-specific ortholog and *P. aeruginosa* TrmV, which introduces m^2^A specifically in tRNAs (33). While *S. aureus* RlmN retains the conserved R204 essential for tRNA recognition, it lacked several residues mediating non-sequence-specific tRNA contacts as observed in the crystal structure of the *E. coli* RlmN-tRNA complex (84) (Figure S6). Furthermore, analysis of electrostatic surface potential revealed that tRNA-binding regions are strongly positive in both *E. coli* RlmN and *P. aeruginosa* TrmV but significantly weaker in *S. aureus* RlmN, potentially limiting its ability to bind tightly tRNA (Figure 3E). Surprisingly, *B. subtilis* and *Enterococcus faecalis* RlmN also exhibit weaker positive binding surfaces (Figure S6), and m^2^A has been found in tRNAs from these species (27, 85). The loss of [m^2^A]37 in *S. aureus* tRNA illustrates the evolution of the enzyme activity across bacteria, which most probably reflects an adaptation of *S. aureus* to its ecological niche.

**Figure 3.**
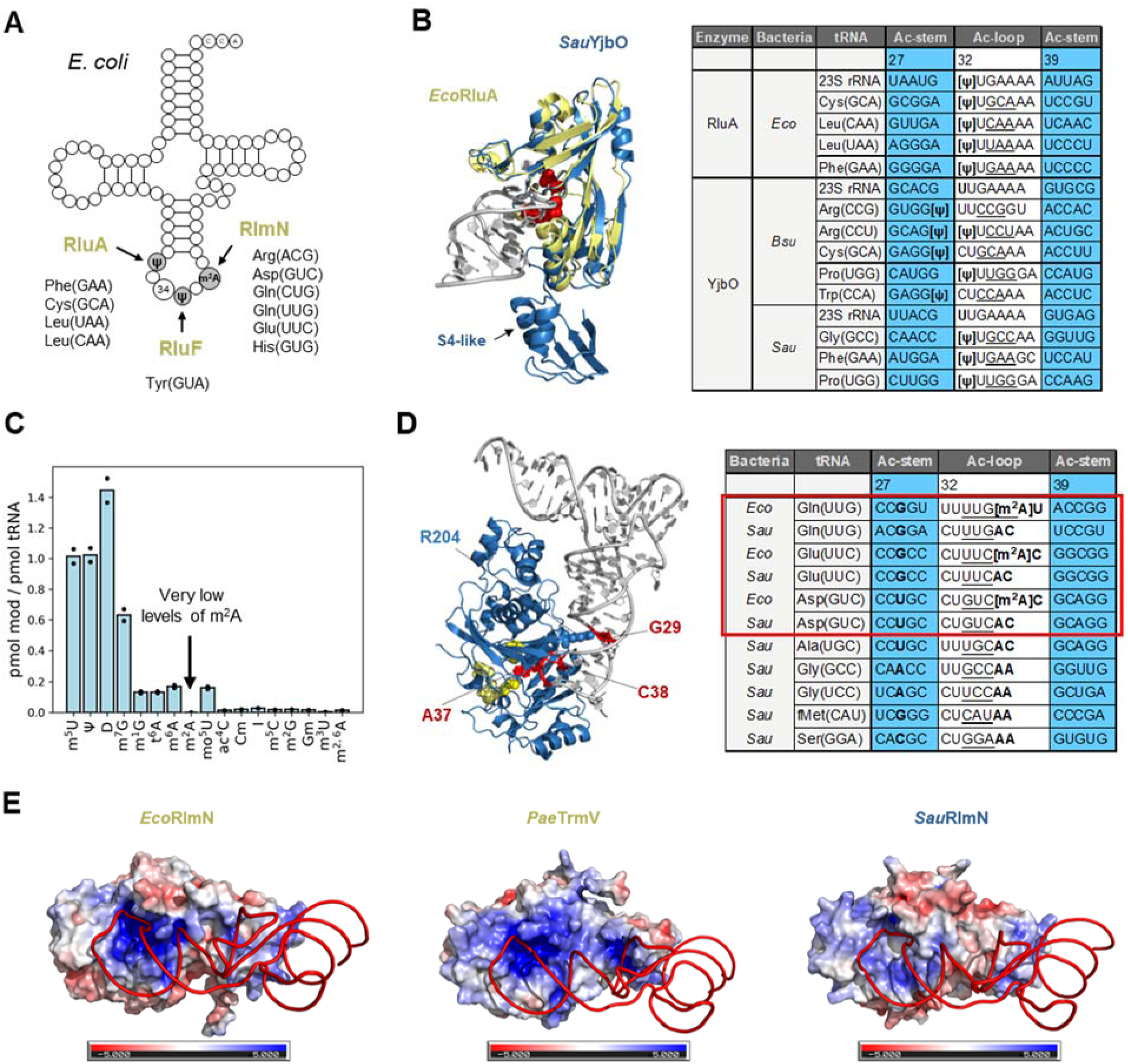
Divergence of dual-specific RNA modification enzymes in *E. coli* and *S. aureus*. (**A**) Three *E. coli* rRNA-modifying enzymes also target tRNA. The distinct tRNA substrates are shown for each enzyme. (**B**) YjbO, responsible for [Ψ]32 in *B. subtilis,* is not an RluA ortholog. *Left:* AlphaFold3 model of *S. aureus* YjbO (blue) superposed to the structure of *E. coli* RluA (green) in complex with tRNA Phe(GAA) (gray) (PDB 2I82). YjbO contains an N-terminal S4-like domain not present in RluA. *Right*: Sequence alignment of RluA and YjbO substrates. Only a 17-nucleotides region encompassing the target position (in bold) is shown. For tRNAs, the anticodon sequence is underlined. For *B. subtilis* and *S. aureus* 23S rRNA, the presence of Ψ at the target position has not been detected. (**C**) LC-MS/MS absolute quantification of nucleosides in *S. aureus* tRNA. The stoichiometric levels of mapped modifications are consistent with their frequency across tRNA species. Levels of m^2^A are even lower than those of nucleosides derived from residual rRNA contamination (m^6, 6^A and m^3^U). (**D**) Conservation of tRNA features recognized by RlmN. *Left:* AlphaFold3 model of *S. aureus* RlmN (blue) in complex with tRNA Glu(UUC) (gray). tRNA nucleotides required for efficient recognition are shown in red. Protein cysteines involved in catalysis and coordination of [4Fe-4S] cluster are represented as yellow and green spheres, respectively. R204, essential for tRNA modification, is also shown as spheres. *Right:* Sequence alignment of *E. coli* RlmN substrates and *S. aureus* tRNAs containing unmodified A37. Potential *S. aureus* RlmN substrates are indicated with a red box. (**E**) Electrostatic surface potentials of bacterial enzymes synthesizing m^2^A. Both dual-specific *E. coli* RlmN (PDB 5HR6) and tRNA-specific *P. aeruginosa* TrmV (AlphaFold3) display RNA-binding surfaces with strong positive potential, whereas *S. aureus* RlmN (AlphaFold3) exhibits markedly lower potentials in these regions. *Eco*: *Escherichia coli*, *Bsu*: *Bacillus subtilis*, *Sau*: *Staphylococcus aureus*, *Pae*: *Pseudomonas aeruginosa*.

### Heterogeneous patterns of A37 methylthiolation

Among the six *S. aureus* tRNAs containing [i^6^A]37, installed by MiaA, only the three tRNAs – Leu(UAA), Leu(CAA) and Tyr(GUA) – are substrates of MiaB and carry [ms^2^i^6^A]37 (Figure 2A). This aligns with the presence of an A31:[Ψ]39 base pair in these tRNAs, proposed to be required for methylthiolation of tRNA^Phe^(GAA) and tRNA^Tyr^(GUA) in *B. subtilis* (86). However, UPLC-MS/MS analysis indicated that [ms^2^i^6^A]37 was most prominent in tRNA^Leu^(UAA), while this modification was detected only rarely and at low intensities in tRNA^Leu^(CAA) and tRNA^Tyr^(GUA), with most fragments carrying [i^6^A]37 (Supplementary Data 1). Comparison of the GLORIseq profiles for these three tRNAs revealed that tRNA^Leu^(CAA) and tRNA^Tyr^(GUA) were likely fully modified at position 37, whereas tRNA^Leu^(UAA) displayed markedly lower scores, suggesting sub-stoichiometric [i^6^A]37 levels (Figure 4A). Interestingly, UPLC-MS/MS clearly detected [ms^2^A]37 within the anticodon loop of tRNA^Leu^(UAA) (Figure 4B), implying that *S. aureus* MiaB can introduce the methylthio group directly onto unmodified A37. Yet, fragments containing only [i^6^A]37 were also observed (Supplementary Data 1), supporting the canonical MiaA-to-MiaB pathway for [ms^2^i^6^A]37 formation (Figure S5).

**Figure 4.**
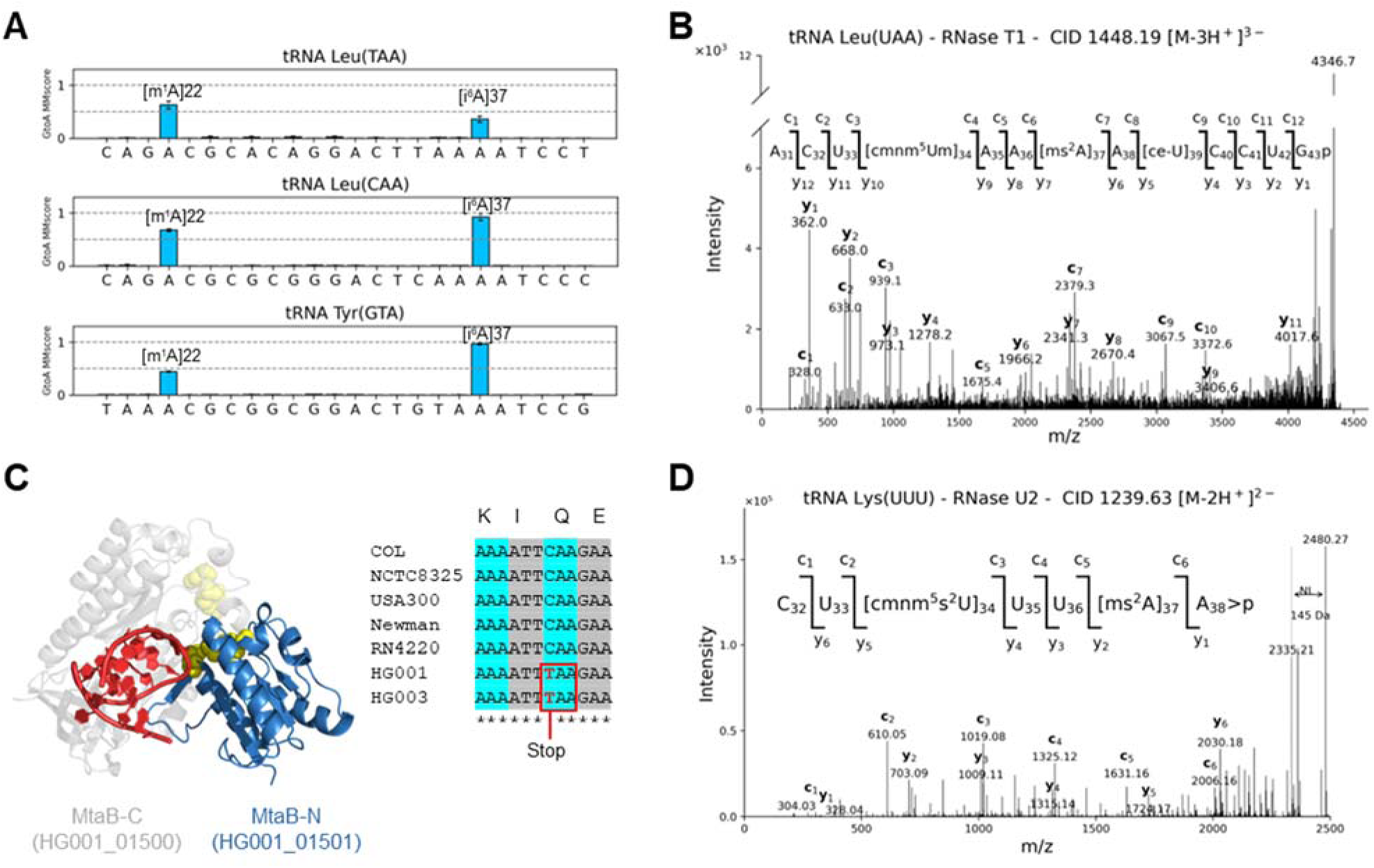
Profiles of A37 methylthiolation in *S. aureus*. (**A**) GLORIseq profiles of tRNAs – Leu(UAA), Leu(CAA), and Tyr(GUA) – showing signals resulting from [m^1^A]22 and [i^6^A]37. Scores close to 1 indicate stoichiometric modification. Signal from [m^1^A]22 arises from partial rearrangement into [m^6^A]22, reflected by lower scores. Levels of [i^6^A]37 in tRNA^Leu^(UAA) are lower than in tRNA^Leu^(CAA) and tRNA^Tyr^(GUA). (**B**) Deconvoluted CID MS/MS spectrum corresponding to fragment ACU[cmnm^5^Um]AA[ms^2^A]A[ce-U]CCUGp (4346.57 Da, *m/z =* 1448.19) derived from RNase T1 digestion of spot VIII-06. Relevant visible *c*- and *y*-series are labeled. (**C**) AlphaFold2 model of *S. aureus* MtaB. In the HG001 strain, only the inactive N-terminal domain (blue) is expressed due to an internal stop codon (sequence alignment). Cysteine residues involved in coordination of [4Fe-4S] clusters are shown as yellow spheres. The non-expressed C-terminal region (light gray) contains one [4Fe-4S] cluster important for catalysis and makes extensive contacts with the tRNA substrate (red). (**D**) Deconvoluted CID MS/MS spectrum showing the presence of [ms^2^t^6^A]37 in tRNA Lys(UUU) purified from USA300 JE2 strain, which express a functional *mtaB* gene. The 2480.27 Da fragment (*m/z =* 1239.63) shows a 145 Da neutral loss typical of t^6^A, resulting in fragmentation of the sequence CU[cmnm^5^s^2^U]UU[ms^2^A]A>p.

Detection of [ms^2^t^6^A]37 in tRNA^Lys^(UUU) was only possible when tRNA was isolated from the *S. aureus* USA300 JE2 strain, as the corresponding *mtaB* gene in HG001 strain was disrupted by a naturally occurring internal stop codon (Figure 4C). The same mutation was also identified in the HG003 strain, which, like HG001, was generated through genetic manipulation of the NCTC8325 parental strain carrying an intact *mtaB* gene (87). Although LC-MS/MS detected expression of the N-terminus of MtaB (Figure 2B), this truncated protein is very likely inactive, as it lacks most of the tRNA-binding surface as well as one of the two [4Fe-4S] clusters involved in catalysis (88, 89). Assuming that MtaB recognizes its substrates through similar determinants as MiaB, the presence of [ms^2^t^6^A]37 was assessed in purified tRNA^Lys^(UUU) and tRNA^Thr^(UGU), which are the only species containing [t^6^A]37 and A31:[Ψ]39 base pairs. Similar to *B. subtilis*, only tRNA^Lys^(UUU) carried [ms^2^t^6^A]37 (Figure 4D), while this modification could not be found in tRNA^Thr^(UGU) (Supplementary Data 1). Notably, methylthiolation was detected in tRNA^Lys^(UUU) only modified with [cmnm^5^s^2^U]34, but not [mnm^5^s^2^U]34, suggesting a possible interdependence between modifications at positions 34 and 37 in this tRNA.

### A single DusB2 enzyme is responsible for tRNA dihydrouridylation

Among the three Dus families, only DusB is present in Gram-positive bacteria, where it occurs as one or more members of the DusB1, DusB2, or DusB3 subgroups (90). In *S. aureus*, two genes are annotated as DusB2, but only one encodes a full-length enzyme (DusB2) containing the catalytic site, whereas the second (DusB2-C) encodes only the C-terminal tRNA binding domain (91, 92) (Figure 5A). The absence of the catalytic site in the truncated protein, together with its lack of detection in the proteomics analysis (Figure 2B), suggested that DusB2 is the sole enzyme responsible for D synthesis, prompting us to generate a Δ*dusB2* mutant to test this hypothesis.

**Figure 5.**
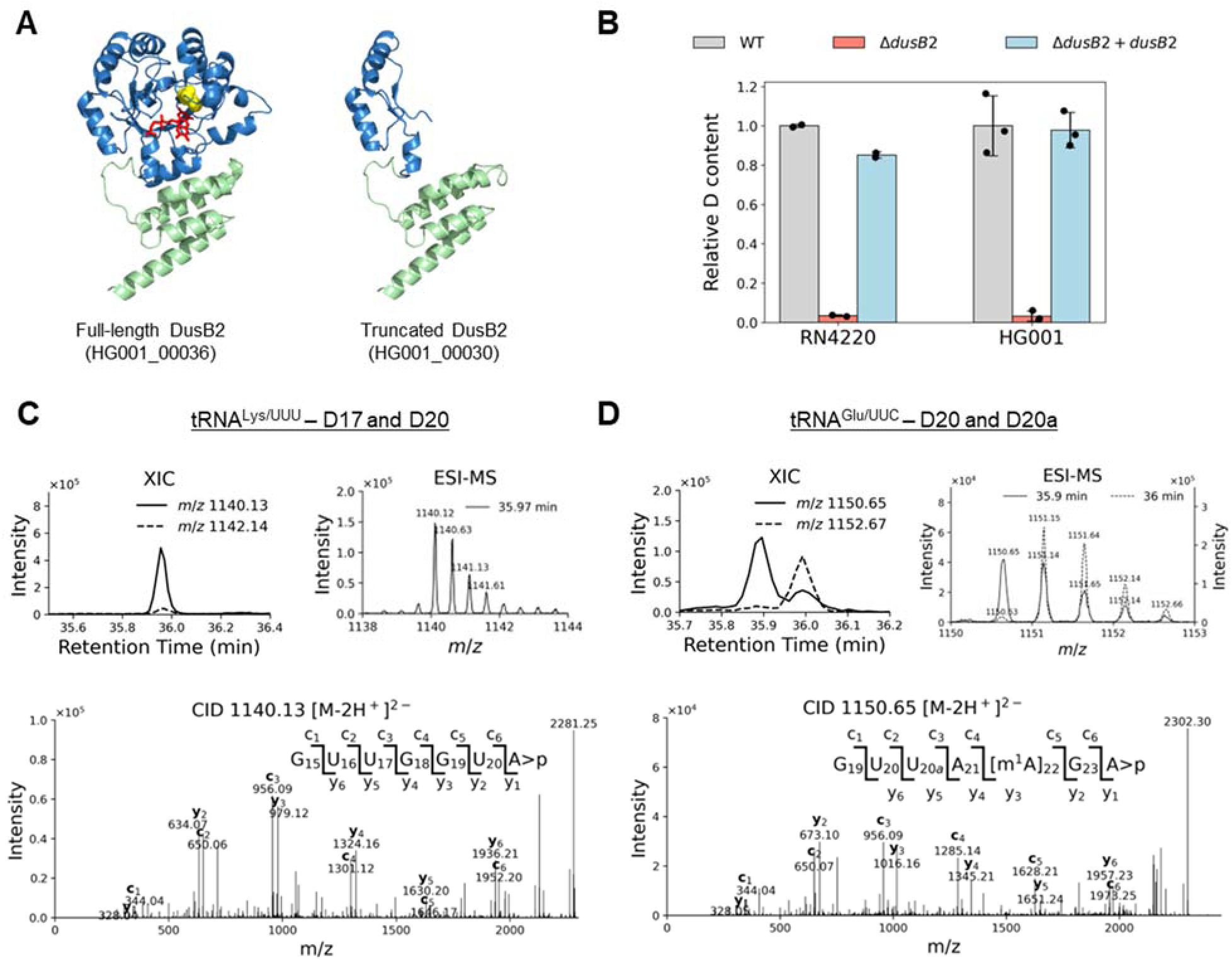
DusB2 is responsible for [D]17, [D]20, and [D]20a in *S. aureus* tRNAs. (**A**) AlphaFold3 models of the two DusB2 proteins encoded in the *S. aureus* HG001 genome. Gene HG001_00036 encodes a full-length DusB2 composed of an N-terminal TIM-barrel domain (purple) containing a catalytic cysteine (yellow spheres) and the binding site for FMN cofactor (red sticks), followed by a C-terminal helical domain involved in tRNA binding (green). The second gene, HG001_00030, encodes only the C-terminal domain and lacks most of the catalytic domain. (**B**) Colorimetric quantification of D in bulk tRNA isolated from WT, Δ*dusB2* and complemented Δ*dusB2* + *dusB2* strains in RN4220 and HG001 backgrounds. D was nearly undetectable in Δ*dusB2* mutants and was restored upon complementation. (**C**) UPLC-MS/MS analysis of tRNA^Lys^(UUU) and (**D**) tRNA^Glu^(UUC) isolated from the RN4220 Δ*dusB2* showing the absence of D in the D-loop. Fragments were generated by RNase U2 digestion of 2D-PAGE spots. For each tRNA, extracted ion chromatogram (XIC), electrospray ionization mass spectrum (ESI-MS), and deconvoluted CID-MS/MS spectra are shown. The 2281.25 Da fragment (*m/z* 1140.13) corresponds to the sequence GUUGGUA>p in tRNA^Lys^(UUU), while the 2302.20 Da fragment (*m/z* 1150.15) matches the sequence GUUA[m^1^A]GA>p in tRNA^Glu^(UUC). XIC and ESI-MS in dashed lines correspond to the expected modified fragments GU[D]GG[D]A>p (*m/z* 1142.14) and G[D][D]A[m^1^A]GA>p (*m/z* 1152.66) for tRNA^Lys^(UUU) and tRNA^Glu^(UUC), respectively. The modified fragment was not detected in tRNA^Lys^(UUU) and the potential signal for the modified fragment in tRNA^Glu^(UUC) corresponded to another sequence (GGAGUCA>p, *m/z* 1151.148).

Deletion of *dusB2* in RN4220 and HG001 strains caused a strong reduction of D levels in bulk tRNA, as quantified by a colorimetric assay (93), which were restored to near wild-type levels upon complementation with a plasmid carrying the *dusB2* gene (Figure 5B). To determinate which specific D sites were affected, individual tRNAs were isolated by 2D-PAGE, digested with RNase U2, and analyzed by UPLC-MS/MS. In the absence of DusB2, tRNA^Lys^(UUU), which normally carries [D]17 and [D]20, yielded only unmodified D-loop fragments (Figure 5C), and tRNA^Glu^(UUC) was likewise devoid of D at positions 20 and 20a (Figure 5D). [D]17 was also found absent in tRNA^Leu^(UAA) and tRNA^Val^(UAC), [D]20 in tRNA^Asn^(GUU) and tRNA^Gly^(GCC), and [D]20a in tRNA^Tyr^(GUA) (Figure S7). Together, these results indicate that D synthesis in *S. aureus* tRNA is mediated by a single multi-site DusB2 able to modify positions 17, 20, and 20a.

Furthermore, our comprehensive mapping of D sites across all *S. aureus* tRNAs revealed distinct dyhidrouridylation patterns, indicating that sequence features within the D-loop determinate the sites targeted by DusB2 (Table 1). The tRNAs carrying only [D]17 never contained U at positions 20 or 20a, whereas those containing only [D]20 and/or [D]20a lacked position 17 or displayed another nucleotide. Notably, tRNA^fMet^(CAU) and tRNA^Pro^(UGG) exhibited unmodified U17 together with a pyrimidine at position 17a, suggesting that this insertion may be a negative determinant for DusB2 modification. In contrast, all other tRNAs with U17, U20 and/or U20a were always modified.

### Non-proteogenic tRNAs are substoichiometrically hypomodified

In addition to the two proteogenic tRNA^Gly^(GCC) “P1” and tRNA^Gly^(UCC) “P2”, *S. aureus* expresses three additional tRNA^Gly^(UCC) species, designated NP1, NP2, and NP3 (Figure 6A), which are involved in the synthesis of pentaglycine bridges in the peptidoglycan cell wall. These non-proteogenic tRNAs are excluded from translation due to the absence of strong EF-Tu determinants (94) and display several unusual sequence features. Notably, they lack the highly conserved G18-G19 and U55 residues in the D and TΨC loops, respectively, and contain an extra C31-G38 base pair that shortens the anticodon loop (95). Moreover, the NP3 species is characterized by a shortened three-base-pair D-stem due to unpaired U10 and U25 (96).

**Figure 6.**
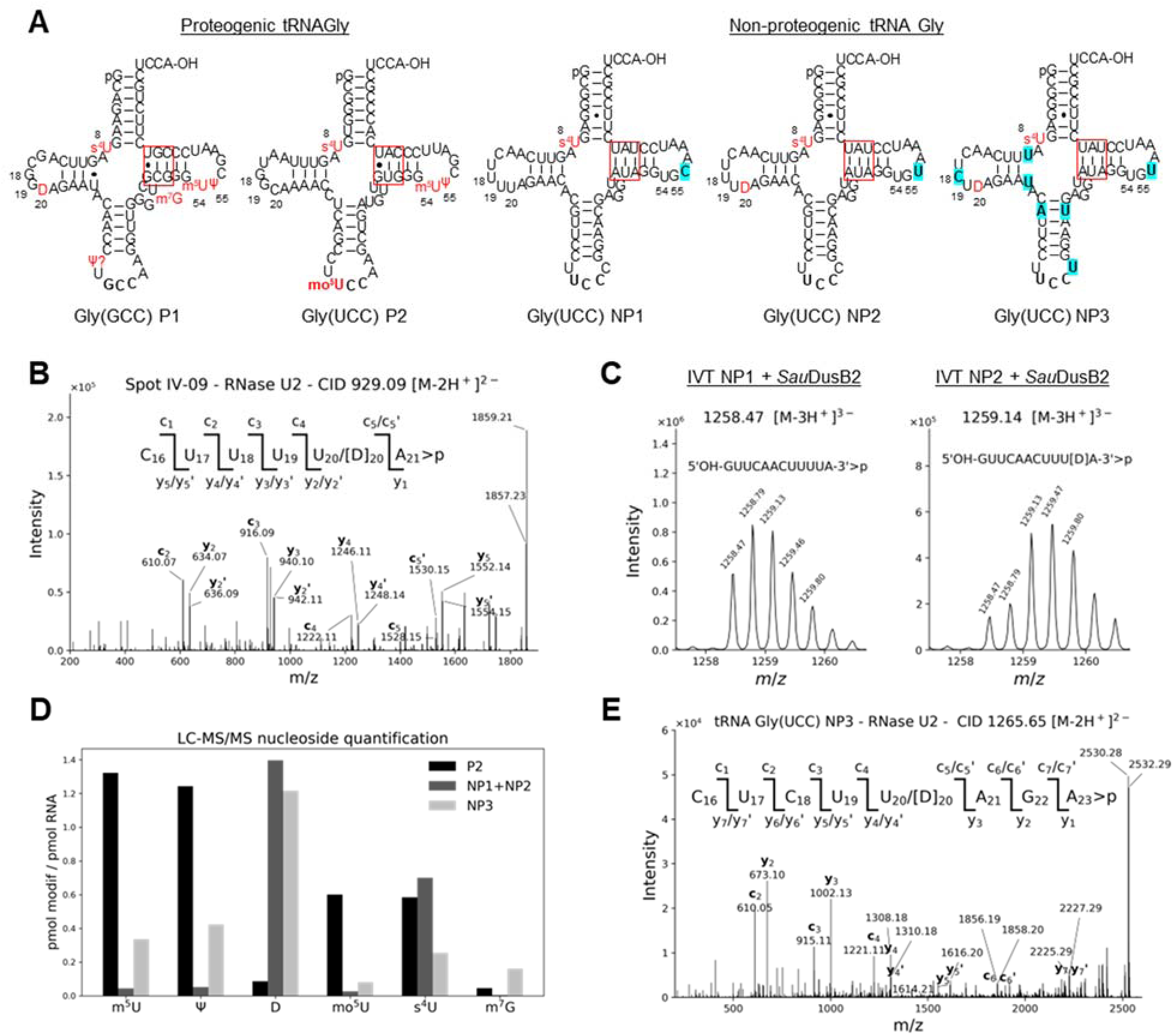
Modification profiles of non-proteogenic *S. aureus* tRNA^Gly^(UCC) species. (**A**) Cloverleaf representation of proteogenic tRNA^Gly^(GCC) P1 and tRNA^Gly^(UCC) P2 and non-proteogenic tRNA^Gly^(UCC) NP1, NP2 and NP3, showing mapped modifications in red. Nucleotides differing among NP1, NP2 and NP3 are highlighted in blue. The anticodon sequence is in bold. EF-Tu binding determinants are indicated with red boxes. Numbering is shown for some relevant positions. (**B**) Deconvoluted CID MS/MS spectrum corresponding to fragments CUUUUA>p (1857.19 Da, *m/z =* 928.088) and CUUU[D]A>p (1859.20 Da, *m/z =* 929.096) derived from RNase U2 digestion of spot IV-09 containing both NP1 and NP2. Relevant *c-* and *y*-series for unmodified and modified (primed) fragments are labeled. (**C**) Dihydrouridylation of synthetic NP1 and NP2 by recombinant *S. aureus* DusB2. Samples were digested with RNase U2 and analyzed by UPLC-MS/MS. The ESI-MS spectra correspond to NP1 (left) and NP2 (right) D-loop fragments. Only NP2 was modified by DusB2. (**D**) Absolute quantification of nucleosides in tRNAs – Gly(UCC) P2 (black), NP1+NP2 (dark gray), and NP3 (light gray) – isolated from an edited *S. aureus* RN4220 strain. In P2, levels of m^5^U and Ψ are stoichiometric, while mo^5^U and s^4^U levels are lower but relevant. In NP1+NP2 and NP3, only D is stoichiometric, and s^4^U modification is partial. NP3 contains appreciable amounts of m^5^U and Ψ, suggesting contamination with proteogenic tRNAs. (**E**) Deconvoluted CID MS/MS spectrum corresponding to fragments CUCUUAGA>p (2530.30 Da, *m/z =* 1264.65) and CUCU[D]AGA>p (2532.32 Da, *m/z =* 1265.65) derived from RNase U2 digestion of the purified NP3 sample.

UPLC-MS/MS analysis of *S. aureus* HG001 non-proteogenic tRNAs revealed the presence of only two modifications, [s^4^U]8 and [D]20, which were not uniformly distributed. NP1 and NP2, differing by a single nucleotide, could not be separated by 2D-PAGE, and the resulting gel spot (IV-09, Figure S1) contained a mixture of both species. In contrast, tRNA^Gly^(UCC) NP3, carrying more sequence differences, was resolved as separate spot (IV-13, Figure S1). While [s^4^U]8 was identified in both samples, [D]20 was only detected in the NP1+NP2 mixture, with fragmentation spectra indicating coexistence of modified and unmodified fragments (Figure 6B). This pattern suggests that either one of the two species, NP1 or NP2, is modified by DusB2, or that both are partially modified. To resolve this ambiguity, we performed an *in vitro* assay using recombinant DusB2 with synthetic NP1 and NP2 substrate, and subsequent UPLC-MS/MS analysis indicated that only NP2 was dihydrouridylated (Figure 6C).

As HydraPsiSeq and BIDseq were inefficient in detecting Ψ in non-proteogenic tRNA^Gly^(UCC) species (Supplementary Data 2), we purified the NP1+NP2 mixture and NP3 using specific biotinylated probes and assessed the presence of this modification by LC-MS/MS nucleoside quantification (Supplementary Data 5). As a control, the same analysis was performed on the proteogenic tRNA^Gly^(UCC) P2. These tRNAs were isolated from an edited *S. aureus* RN4220 strain lacking one gene copy of tRNA^Gly^(GCC) P1, which expresses higher levels of the non-proteogenic species (40) and reduces potential contamination from P1. This analysis confirmed that Ψ was present at stoichiometric levels in P2, but not in the NP1+NP2 mixture (Figure 6D). Although higher levels of Ψ were detected in NP3, this likely reflects contamination with proteogenic tRNAs, as indicated by the concurrent detection of similar levels of m^5^U. Consistent with our mapping data in the HG001 strain, higher levels of D were detected in the NP1+NP2 sample, but not in P2. Unexpectedly, substantial amounts of D were also observed in NP3, contrasting with our data in HG001 indicating unmodified D-loop fragments (Figure S8). Further oligonucleotide UPLC-MS/MS analysis of the purified NP3 confirmed the presence of [D]20, albeit at sub-stoichiometric levels, as CID-MS/MS spectra indicated a mixture of modified and unmodified fragments (Figure 6E). Interestingly, nucleoside quantification also revealed partial modification with [mo^5^U]34 in tRNA^Gly^(UCC) P2, and with [s^4^U]8 in all P2, NP1+NP2, and NP3 samples.

Altogether, these results indicate that non-proteogenic tRNA^Gly^(UCC) exhibit minimal and heterogenous modification profiles, with [s^4^U]8 present in all species and [D]20 potentially restricted to specific tRNAs. This is supported by comparative analysis of non-proteogenic tRNA^Gly^(UCC) sequences from other *Staphylococci*, indicating that U20 is not strictly conserved (Figure S8). Finally, Nanopore sequencing revealed that non-proteogenic tRNA^Gly^(UUC) species were among the less abundant tRNAs in *S. aureus* HG001, with NP3 being the rarest of the three tRNAs (Figure S9).

### Codon decoding profiles and tRNA abundance

Inspection of the decoding capacities of the *S. aureus* tRNA set indicates that multiple codon boxes are decoded by single tRNA species (Figure 7B). Only the 4-fold degenerate Gly-GGN, Leu-CUN, Arg-CGN, and Ser-UCN boxes, as well as the split Leu-UUN box, are decoded each by two distinct isoacceptors. A special case is the Ile/Met box, which requires two tRNA species to decode the three Ile codons. Notably, superwobbling (97) appears to occurs for Ala, Pro, Thr, and Val, since a single tRNA carrying [mo^5^U]34 appears sufficient for decoding of the four synonymous codons for these amino acids.

**Figure 7.**
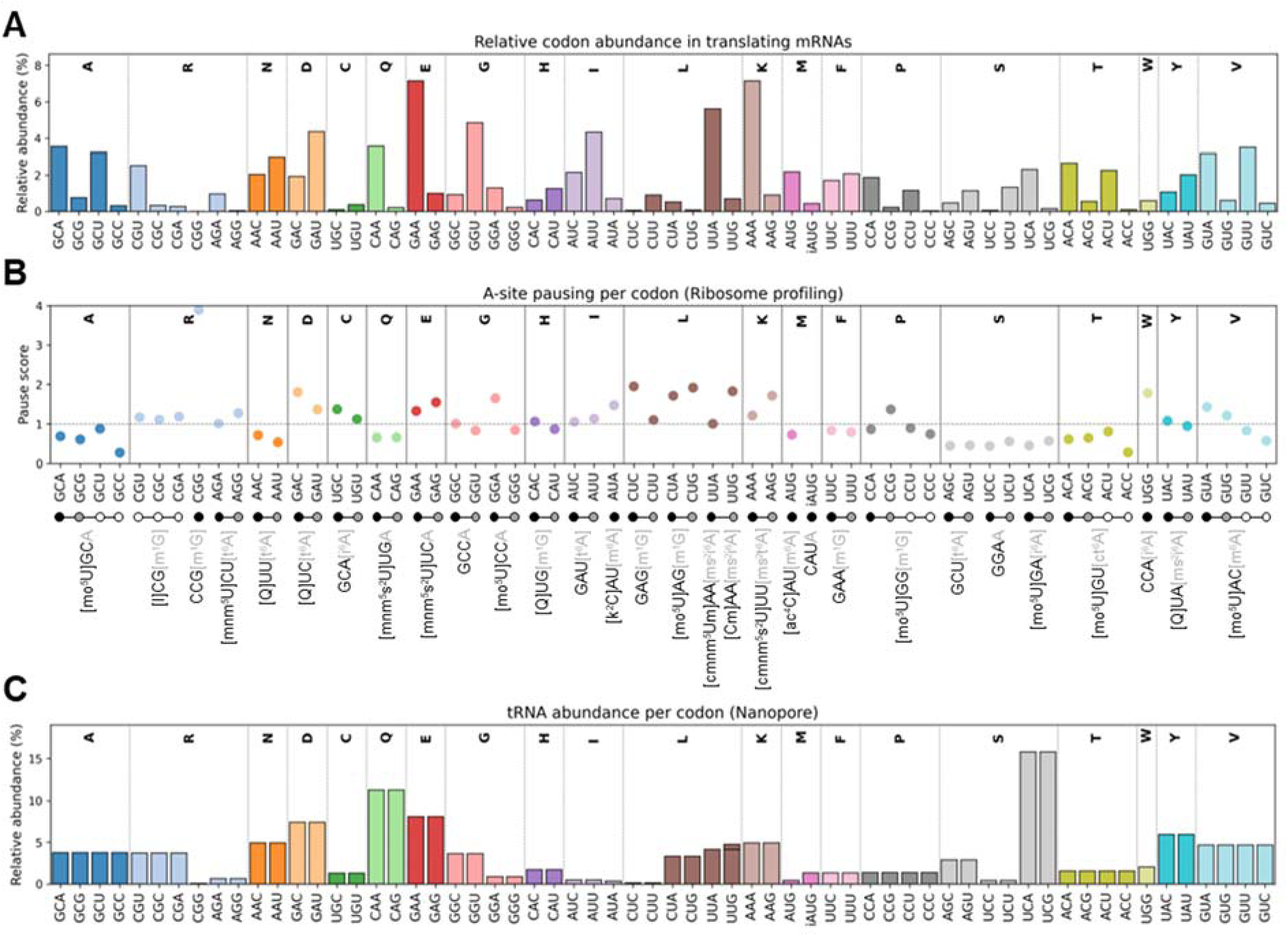
Dynamics of codon decoding in *S. aureus* HG001. (**A**) Relative abundance of codons in translating mRNAs. Codon counts from genes in ribosome-protected fragments was weighted by transcript TPM values to account for expression levels. The resulting profile closely mirrors codon usage derived from genomic coding sequences. Variations in the abundance of synonymous codons reflect the higher AT content of the *S. aureus* genome. (**B**) Decoding capacities of *S. aureus* tRNAs and codon decoding rates. The 61 + 1 (initiator AUG) sense codons are displayed along the horizontal plot axis. The tRNA anticodon sequences in bold are placed below their cognate codons, with symbols indicating canonical (●), wobble (●), or superwobble (○) base-pairing interaction at the third codon position. The nucleotide at position 37, which also contributes to codon decoding, is shown in gray. For tRNA^Lys^(UUU), [ms^2^t^6^A]37 is indicated but this position in HG001 carries only [t^6^A]37. The plot shows the global A-site pausing score for each codon, as determined by ribosome profiling. Higher scores indicate slower tRNA decoding. (**C**) Relative abundance of *S. aureus* tRNAs determined by direct Nanopore sequencing. For each codon, the abundance of its cognate tRNA(s) is shown. Because a single tRNA can decode multiple codons, identical abundance values appear for these codons. Note that UUG is decoded by two distinct tRNAs, and their abundances were stacked in the bar plot. The lower abundance corresponds to tRNA^Leu^(CAA). In panels (A) – (C), codons encoding the same amino acid are colored according to a common color code.

The *S. aureus* tRNA decoding rates were investigated using a recently adapted codon-resolution ribosome profiling method (98). In this approach, translating polysomes are digested with an RNase, and the resulting ribosome-protected mRNA fragments are sequenced and mapped to the coding sequences, revealing ribosome occupancy of codons at the A, P, and E-sites. Higher occupancies are generally associated with slower translation or ribosome pausing at these codons. This experiment was performed in the absence of procedures inducing pausing artifacts, such as antibiotic treatment and cell harvesting by filtration, thereby ensuring capture of biologically relevant translation dynamics (99). Consistently, we did not observe strongest A- and E-site pauses at Gly and Ser codons, which are typically associated with such technical biases (Figure S9). Here, we focused specifically on A-site pausing, which generally reflects tRNA decoding rates (Figure 7B).

In optimal rich medium, Ser codons, recognized by three distinct tRNA species, showed the fastest decoding rates. Conversely, the slowest decoding tRNAs were tRNA^Arg^(CCG) and tRNA^Trp^(CCA), both reading single codons. For several Leu codons, strong A-site pauses were also observed, but only for specific codons recognized by the same tRNA. For instance, tRNA^Leu^(GAG) decoded faster CUU than CUC, whereas tRNA^Leu^(UAA) read UUA more efficiently than UUG, despite the presence of tRNA^Leu^(CAA), also recognizing UUG. Although such differences were less evident for most of the other synonymous codons, which were decoded at more similar rates, the fastest codon-anticodon interactions tended to involve either canonical U34-A3 or wobble G34-U3 base pairings. Only tRNA^Gly^(UCC) showed the opposite behavior, decoding GGG faster than GGA through an [mo^5^U]34-G3 wobble base pair. These decoding dynamics showed correlation with mRNA usage of codons recognized by the same tRNA, as the most frequent tended to be decoded faster (Figure 7A). Interestingly, all codons within each of the 4-fold degenerate boxes read by a single tRNA (Ala, Pro, Thr, Val) were decoded at comparable rates, with C-ending codons consistently being the fastest, indicating that superwobble recognition *in vivo* is very efficient.

Since codon decoding rates are strongly influenced by tRNA availability, *S. aureus* tRNA species were quantified by Nanopore sequencing (Figure 7C) to relate their abundance to the decoding patterns observed by ribosome profiling. Although no general correlation emerged between tRNA abundance and A-site pausing, the most abundant tRNA^Ser^(UGA) corresponded to the fastest-decoded codons, whereas tRNA^Arg^(CCG) was the less abundant, consistent with its markedly slower decoding rate. In contrast, tRNA^Ser^(GGA) was detected at very low levels, yet the UCC and UCU codons displayed very high decoding rates in ribosome profiling, suggesting that tRNA^Ser^(UGA) may also participate in their decoding through superwobble interaction. Interestingly, this tRNA carries [mo^5^U]34 as found for the four tRNA – Ala(UGC), Pro(UGG), Thr(UGU), and Val(UAC) –, all forming superwobble interactions. Similar discrepancies were observed for Leu-CUN and Gly-GGN boxes, where codons read by the lower-abundant tRNA were decoded as fast as those recognized by the most abundant species. While potential tRNA^Leu^(UAG) superwobbling could account for the differences among Leu-CUN codons, this explanation seems unlikely for Gly-GGN codons, as tRNA^Gly^(GCC) is unable to compensate for tRNA^Gly^(UCC) *in vivo* (100).

## DISCUSSION

The role of tRNA modifications as key regulators of bacterial physiology and pathogenicity, particularly under stress, has only been recognized in recent years (11, 101). Their dynamic nature in response to environmental cues enabled integration into regulatory networks modulating translation through effects on tRNA stability, aminoacylation, recognition, and codon decoding (102–105). Yet, our understanding of epitranscriptomic regulation remains limited, as the full repertoire of tRNA modifications and their enzymes is known for only a few bacterial species (106, 107). In this context, several studies have characterized the global tRNA modification landscapes across diverse bacteria (27–29, 31–34, 108–111), revealing both conserved activities and numerous species-specific features, some with potential therapeutic relevance (60, 112–114). Motivated by these findings, we performed a detailed analysis in *Staphylococcus aureus*, using oligonucleotide mass spectrometry to confidently map modifications in each individual tRNA species of this clinically important pathogen.

Although oligonucleotide mass spectrometry provides high-confidence information about the nature and position of modified nucleotides, it has some important limitations (115). First, it requires isolation of individual tRNA species, which is labor-intensive and can limit detection of rare tRNAs. Second, it cannot distinguish between isobaric modifications such as different methylated nucleotides or U and Ψ. Even with improvements made to address these challenges, complete and reliable mapping of *S. aureus* tRNA modifications would not have been possible without integrating deep-sequencing approaches designed to detect specific types of modifications. The complementarity of the two methods is exemplified by the lower abundant tRNA^Ile^(CAU), which became accessible through our improved purification method (Figure 1E). Alone, mass spectrometry detected a methylated adenosine at position 37, but the assignment of [m^6^A]37 can only be inferred from data reported in other organisms (56). In contrast, GLORIseq, which specifically detects N^6^-modified adenosines, produced a signal at the same position that could be misinterpreted as t^6^A, which is more commonly occurring in tRNA^Ile^(CAU). Only by combining the two approaches, we managed to unambiguously assign [m^6^A]37, underscoring the importance on integrating orthogonal approaches for accurate modification mapping. This is even more critical for Ψ detection, as illustrated by the challenging assignment of [Ψ]32 sites in *S. aureus*, mapped confidently only in tRNA^Pro^(UGG) but remaining tentative for tRNA^Gly^(GCC) and tRNA^Phe^(GAA). In such cases, the use of mutants of modification enzymes becomes crucial to validate modified positions by deep-sequencing (116, 117).

Hence, our analysis provides a complete map of tRNA modifications in this major pathogen (Table 1). Compared to other bacteria, particularly Gram-negative species, *S. aureus* tRNAs are clearly less extensively modified. Viewed globally, the modified positions are well conserved, with only a few modifications, such as [m^1^A]22, [mo^5^U]34, and [ms^2^t^6^A]37, characteristic of Gram-positive bacteria (27). However, our detailed analysis at the level of individual tRNAs, which enabled assignment of tRNA-modifying enzymes, revealed several distinctive features that would otherwise remain unnoticed. One major difference concerns the modifications introduced by enzymes targeting both tRNA and rRNA in other bacteria. In *S. aureus*, the pseudouridine synthase YjbO, likely responsible for [Ψ]32 in tRNA^Pro^(UGG), does not appear to be an ortholog of the *E. coli* dual-specific RluA enzyme, which introduces the same modification in tRNAs and [Ψ]746 in 23S rRNA. Structurally, YjbO differs from RluA by the presence of an N-terminal S4-like domain characteristic of other bacterial pseudouridine synthases such as RsuA, RluA, and RluC, all of them targeting rRNA. In addition, the tRNA substrates modified by RluA and YjbO do not overlap and display distinct sequence motifs surrounding the target U32, indicating divergence in substrate recognition. These observations, together with the absence of the equivalent [Ψ]746 site in Gram-positive bacteria (79, 80), suggest that YjbO may represent a novel pseudouridine synthase, either functioning specifically on tRNA or as a dual-specific enzyme acting on other rRNA positions or even RNA classes, as has been observed for RluA (118).

A more intriguing aspect of tRNA modification in *S. aureus* involved the methyltransferase RlmN. Although the enzyme is conserved and modifies the same rRNA site as its dual-specific *E. coli* counterpart (82), it does not appear to install [m^2^A]37 in tRNAs. This was initially puzzling as sequence and structural comparisons with dual-specific orthologs suggested that *S. aureus* RlmN and its potential tRNA substrates possess all features required for tRNA methylation. However, closer examination of the enzyme revealed differences in the conservation of residues mediating non-sequence specific RNA contacts, which may impair tRNA recognition. Interestingly, similar patterns were observed in the RNA binding surface of RlmN in *B. subtilis* an *E. faecalis*, two Gram-positive bacteria in which m^2^A has been detected in tRNA (27, 85). This suggests that *S. aureus* RlmN may still be able to modify tRNAs, but only under specific conditions, potentially involving increased expression to compensate for its apparently lower tRNA affinity. The unexpected absence of [m^2^A]37 in *S. aureus* tRNAs provides a unique opportunity to investigate the specific role of 23S rRNA [m^2^A]2530 in translation, without confounding effects resulting from differential tRNA modification status. Interestingly, a mutation in *rlmN* gene identified in a clinical *S. aureus* isolate abolished the [m^2^A]2530 modification, had little effect on bacterial growth, but increased resistance to linezolid (82). These findings underscore the importance of understanding the evolutionary dynamics of bacterial modification enzymes, as alterations in modification profiles may promote antibiotic resistance with minimal fitness cost, facilitating their rapid dissemination among bacterial populations.

Detailed inspection of individual tRNAs also evidenced heterogenous modifications profiles, particularly in tRNA^Leu^(UAA) and tRNA^Lys^(UUU). Unlike *E. coli*, A37 methylthiolation in *S. aureus* appears to be largely restricted to these two tRNA species, which are modified by MiaB and MtaB, respectively. Previous studies in Gram-negative bacteria established that [ms^2^i^6^A]37 synthesis by MiaB requires prior [i^6^A]37 modification by MiaA (88, 119); however, here we detected [ms^2^A]37 in tRNA^Leu^(UAA), suggesting that MiaB in *S. aureus* can act on unmodified A37. This notion is supported by the accumulation of ms^2^A observed in *Streptomyces albidoflavus* Δ*miaA* upon MiaB overexpression (31), which correlates with MiaB being the most abundant tRNA-modifying enzyme in the *S. aureus* translatome (Figure 2B). Interestingly, GLORIseq indicated that tRNA^Leu^(UAA) was the only tRNA partially modified by MiaA, likely reflecting competition between MiaA and MiaB and the potential inability of MiaA to act on [ms^2^A]37. Intriguingly, [Um]34 in tRNA^Leu^(UAA), detected by RiboMethSeq at apparently stoichiometric levels (Figure 1D), co-occurred with [ms^2^A]37 (Figure 4B), contrasting with the reported requirement of [i^6^A]37 for TrmL modification (65, 75). This dependency seems to hold true for *S. aureus*, as the only TrmL substrates, tRNA^Leu^(UAA) and tRNA^Leu^(CAA), also contain [i^6^A]37, whereas tRNA^Phe^(GAA), which in *B. subtilis* carries [ms^2^i^6^A]37 and thus [Gm]34 (75), remains unmodified by TrmL in *S. aureus*, likely due to the presence of [m^1^G]37. Although further experiments will be required to clarify the substrate specificity of *S. aureus* MiaB, including its preference for tRNA^Leu^(UAA) over tRNA^Leu^(CAA) and tRNA^Tyr^(GUA), these observations suggest a more complex interdependency between A37 hypermodification and the nucleotide at position 34, which also extends to tRNA^Lys^(UUU). Notably, within the xm^5^U synthesis pathway (Figure S5), tRNA^Leu^(UAA) and tRNA^Lys^(UUU), both carrying methylthiolated A37, are the only species in which modification appears to halt at the cmnm^5^U intermediate. In *E. coli*, tRNA^Lys^(UUU), which is not methylthiolated at [t^6^A]37, is modified up to the final [mnm^5^s^2^U]34 product (120), but in *B. subtilis*, the same tRNA carries [ms^2^t^6^A]37 (121) and predominantly retains [cmnm^5^s^2^U]34 (122). In this context, the presence of [mnm^5^s^2^U]34 in HG001 tRNA Lys(UUU) may have resulted from MtaB inactivation in this strain.

The repertoire of tRNA-modifying enzymes for the existing modifications in *S. aureus* is overall conserved, with several distinctive features of Gram-positive bacteria (57, 66, 86, 123). Particularly notable among these are the dedicated cysteine desulfurases NifZ and YrvO, which support [s^4^U]8 (ThiI) and [s^2^U]34 (MnmA) synthesis, respectively (124, 125), along with two independent MnmL and MnmM enzymes enabling conversion of cmnm^5^U into mnm^5^U within the xm^5^U pathway (126, 127). A feature more specific to *S. aureus* is the presence of two enzymes, QueG and QueH, using distinct cofactors to catalyze redundantly the last step of [Q]34 synthesis (76). Our time-course proteomics experiment revealed distinct expression patterns among the *S. aureus* tRNA-modifying enzymes (Supplementary Data 3), with some expressed earlier (*e.g.* TruB, TrmK, ThiI) than others (*e.g.* TrmR, TmcAL, TsaC) and related enzymes showing coordinated expression (*e.g.* MiaA, MiaB, TrmL). However, the most distinctive *S. aureus* enzyme is DusB2, responsible alone for D synthesis at positions 17, 20, and 20a. Our comprehensive mass spectrometry mapping of D sites suggested that sequence features within the D-loop determine the sites targeted by DusB2, with position 17a potentially acting as a negative determinant. Interestingly, non-proteogenic tRNA^Gly^(UCC), which lack canonical D- and TΨC-interactions, were only partially modified by DusB2, which may reflect subtle structural differences in these unconventional tRNAs. Remarkably, recombinant *S. aureus* DusB2 modifies quite efficiently synthetic non-proteogenic tRNA^Gly^(UUC) NP2 (Figure 6C), in contrast to other Dus enzymes acting preferentially on modified tRNA (92, 128). Another distinctive *S. aureus* feature is the presence of a second inactive DusB2 consisting only in the C-terminal tRNA binding domain, whose location adjacent to genes involved in sulfide stress response (129, 130) (Figure S4) might suggest a potential role in modulation of D synthesis under these conditions.

*S. aureus* encodes fewer tRNA species than the 32 theoretically required to decode all 61 sense codons under the classical wobble rules (131), highlighting the importance of tRNA modification status for efficient codon decoding. Our ribosome profiling A-site pausing analysis suggested that most synonymous codons recognized by the same tRNA are read at comparable rates, with a slight preference for A- and U-rich codons – more frequently used due to the genome’s low GC content. Notably, the four tRNAs – Ala(UGC), Pro(UGG), Thr(UGU), and Val(UAC) –, all carrying [mo^5^U]34, efficiently decoded all four synonymous codons for their respective amino acids, including C-ending triplets, which are considered to form less favorable superwobble interactions (132, 133). Nanopore tRNA sequencing correlated the pronounced ribosome pausing at CGG codons with the low abundance of tRNA^Arg^(CCG), in line with the notion that rare codons are translated more slowly (134). It also revealed discrepancies between tRNA abundance and the decoding speed of Ser-UCN codons, suggesting that tRNA^Ser^(UGA), carrying [mo^5^U]34, substantially contribute to reading of UCC and UCU codons despite the presence of tRNA^Ser^(GGA), which is better adapted to these codons but is present at much lower levels. Perhaps the most outstanding feature in *S. aureus* is the modification at position 34 in tRNA^Gly^(UCC), which carries mo^5^U rather than the mnm^5^U observed in most bacteria, including *B. subtilis* (56). Unlike [mnm^5^U]34, which restricts pairing to GGA and GGG codons (135), the presence of [mo^5^U]34 likely extends the decoding capacities of tRNA^Gly^(UCC) to support recognition of GGC and GGU codons by tRNA^Gly^(GCC). Consistent with this, deletion of one of the two gene copies of tRNA^Gly^(GCC) in *S. aureus* RN4220 did not cause translation defects and led to increased levels of tRNA^Gly^(UCC), likely compensating for the loss (40). However, our data evidenced atypical decoding properties for tRNA^Gly^(UCC), as it recognized the least frequently used GGG codon more efficiently than GGA (Figure 7). While tRNA^Leu^(UAG) and tRNA^Ser^(UGA), both carrying [mo^5^U]34, were more abundant than their corresponding G34-containing isoacceptors, this was not the case for tRNA^Gly^(UCC), which was present at lower levels than tRNA^Gly^(GCC). Despite its lower abundance, tRNA^Gly^(UCC) decoded GGG at speed comparable to that of codons read by the more abundant tRNA^Gly^(GCC), whereas GGA remained translated more slowly. This discrepancy points to a potential role of [mo^5^U]34 in modulating the recognition of specific Gly codons, a hypothesis that will, of course, require more detailed experimental validation.

In summary, our integrative analysis provides a comprehensive view of the full set of tRNA modifications and their corresponding enzymes in *S. aureus*, delivering high-confidence modification maps for each tRNA species, including non-proteogenic tRNA^Gly^. While the overall modification repertoire is conserved in Gram-positive bacteria, examination of individual tRNA modification profiles uncovered significant *S. aureus*-specific features, including enzymes with distinct substrate specificities, coordinated and interdependent modification pathways, and a complex interplay between tRNA abundance, modification status, and codon decoding dynamics. Importantly, this study demonstrates the value of integrating complementary mass spectrometry and deep-sequencing analyses to achieve accurate and comprehensive tRNA modification mapping. Our work establishes a foundation for further research aimed at dissecting the roles of specific tRNA modifications in *S. aureus* physiology, stress adaptation, and pathogenesis.

## MATERIAL AND METHODS

### Bacterial strains and culture conditions

*Staphylococcus aureus* was manipulated in a BioSafety Level 2 Laboratory for class II pathogens. Strain HG001, a derivative of the RN1 (NCTC8325) strain with restored *rsbU* (SigB activator), was used for tRNA modification analysis by UPLC-MS/MS and deep-sequencing. Strain USA300 JE2 was employed to verify the presence of [ms^2^t^6^A]37, as the *mtaB* gene responsible for this modification is truncated in HG001. Mutant and complemented Δ*dusB2* strains in RN4220 and HG001 backgrounds were used to assess tRNA dihydrouridylation. Glycerol stocks were streaked onto blood agar plates (HG001 and USA300 JE2) or BHI agar plates containing 10 μg/mL chloramphenicol (Δ*dusB2* strains) and incubated overnight at 37°C. The following day, single colonies were inoculated into BHI, supplemented with antibiotic when needed, and incubated overnight at 37°C with agitation at 180 rpm. Overnight cultures were then diluted to OD_600 nm_ 0.05 in fresh BHI and grown under the same conditions for approximately 4 h to reach late exponential phase (OD_600 nm_ 2.0 - 3.0). Bacteria were harvested by centrifugation at 2100g for 10 min at 4°C and stored at -20°C until use. The culture volume was adjusted according to RNA requirements, with 1 L cultures in 5-L flasks used for large-scale RNA preparations and 25 mL cultures in 100-mL flasks for smaller preparations.

### Generation of ΔdusB2 mutants and complementation

The *dusB2* gene in *S. aureus* RN4220 (WKU35_RS00190) and HG001 (BSR30_RS00205) was deleted using the pCasSA CRISPR-Cas9 system as described in (136). Guide RNA (gRNA) sequences specific to the target gene were designed with the CCTop-CRISPR/Cas9 target online predictor (137). Primers used for cloning, screening and DNA sequencing are listed in Table S1. Construction of the pCasSA-UPDN-gRNA plasmid, carrying both the homologous recombination template (UPDN) and the gRNA, was performed in two steps. First, the 1-kb upstream (UP) and downstream (DN) flanking regions of *dusB2* in HG001 were fused by overlap PCR using primers Dus_5UP_F, Dus_5UP_R, Dus_3DN_F and Dus_3DN_R, and the resulting 2-kb UPDN fragment was ligated into the pCasSA vector *via* XhoI and XbaI restriction sites. Second, primers gRNA_DusB2_F and gRNA_DusB2_R specific to the *dusB2 locus* were phosphorylated, annealed, and inserted into pCasSA-UPDN by Golden Gate assembly using the two BsaI restriction sites. Cloning was carried out in *E. coli* TOP10 strain, with selection on LB agar plates containing 50 μg/mL kanamycin at 30°C for 24-36 h. The resulting plasmid pCasSA-UPDN-gRNA (1 - 5 μg) was first electroporated (2.5 kV, 5 ms) into *S. aureus* RN4220 and subsequently into HG001 to generate the Δ*dusB2* mutant strains. Transformants were selected on TSB agar plates supplemented with 10 μg/mL chloramphenicol at 30°C for 24-36 h. Several colonies were inoculated into 3 mL TSB containing 10 μg/mL chloramphenicol and incubated overnight at 30°C under agitation. The following day, 3 μL of each culture was diluted into 3 mL BHI without antibiotic, incubated at 42°C for 7 h, and streaked onto TSB agar plates with or without 5 μg/mL chloramphenicol to confirm plasmid curing. Genomic DNA was then extracted, and clones carrying the desired gene deletion were identified by PCR using primers Dus_up and Dus_down, and further confirmed by Sanger sequencing. Complementation of Δ*dusB2* mutants was achieved using plasmid pCN38-DusB2, which carries the *dusB2* gene along with 388 nucleotides upstream and 100 nucleotides downstream the coding sequence. Primers SphI_P_DusB2_F and KpnI_TT_DusB2_R were used to amplify the gene and add restriction sites for cloning. The empty plasmid pCN38, conferring chloramphenicol resistance, was also transformed in the Δ*dusB2* mutants.

### Total RNA extraction

RNA extraction was performed from bacterial pellets obtained from either 25-mL or 500-mL cultures. Parameters indicated in parentheses correspond to large-scale RNA preparations. Bacteria were resuspended in 0.5 mL (6 mL) PBS and transferred into a 2-mL (50-mL) tube containing 0.6 g (6 g) of Lysis Matrix B (MP Biomedicals) and 0.5 mL (6 mL) of acidic phenol:chloroform:isoamyl alcohol 25:24:1 (PCI). Cells were mechanically disrupted for 40 s at 6 m/s using the FastPrep-24 system (MP Biomedicals), and the tubes were centrifuged at 16000g (8370g) for 10 min at 4°C to remove beads and cell debris. The aqueous phase was transferred to a new 1.5-mL (50-mL) tube and re-extracted with an equal volume of acidic PCI by vortexing for 10 s. After centrifugation under the same conditions, the upper aqueous phase was collected and extracted sequentially with an equal volume of cold chloroform and then chloroform:isoamyl alcohol (19:1). RNA was precipitated by adding 1/10 volume of 3 M sodium acetate (pH 5.2) and 3 volumes cold absolute ethanol, followed by incubation overnight at -20°C. RNA was pelleted by centrifugation at 16000g (8370g) for 30 min (45 min) at 4°C, washed with 0.5 mL (5 mL) cold 80% ethanol, air-dried for 20 min (1 h) at 20°C, and resuspended in 50 μL (500 μL) milli-Q water.

### Purification of bulk tRNA

Anion exchange chromatography was used to enrich for tRNA, removing most of the ribosomal RNA and DNA. Total RNA (up to 6 mg) was applied to a 1-mL Fast Flow DEAE column (Cytiva) equilibrated in 100 mM HEPES-KOH (pH 7.4). The column was washed with 10 mL of equilibration buffer, followed by 10 mL of the same buffer supplemented with 100 mM NaCl, and bound RNA was eluted using a linear NaCl gradient from 100 to 1000 mM. Fractions enriched in tRNA, identified by agarose gel electrophoresis, were pooled, ethanol-precipitated, and resuspended in 400 μL milli-Q water. Alternatively, size-exclusion chromatography was used to prepare purer tRNA from small-scale RNA preparations. Total RNA was first treated with DNase I to remove residual DNA. The reaction was carried out in a final volume of 100 μL containing 50 U of RNase-free DNase I (Roche) in the appropriate buffer and incubated for 2 h at 20°C. DNA-free RNA was subsequently extracted with an equal volume of acidic PCI, followed by chloroform, ethanol-precipitated, and resuspended in 400 μL milli-Q water. The RNA sample (up to 1.5 mg) was applied onto a Superose 6 10/300 column (Cytiva) equilibrated and eluted at 0.4 mL/min with 300 mM ammonium acetate (pH 6.8). Fractions containing pure tRNA were pooled, ethanol-precipitated, and resuspended in 20 - 50 μL of milli-Q water.

### Chromatographic fractionation of tRNA species

RNA was extracted from five bacterial pellets obtained from 500-mL cultures harvested at late exponential phase. DNA and ribosomal RNA were first removed by DEAE anion exchange chromatography, and total tRNA was subsequently purified by size-exclusion chromatography under the conditions described above. All tRNA preparations were pooled (3 mg total) and applied to a DeltaPak C4 column (300 Å, 15 μm, 7.8 mm x 300 mm) equilibrated with buffer A (10 mM sodium phosphate pH 7.0, 1 M sodium formate, and 8 mM MgCl_2_). The column was washed with 7 mL of equilibration buffer, and bound tRNA was eluted using a linear gradient of buffer B (10 mM sodium phosphate pH 7.0, 20% methanol) from 0 to 100%. Fractions of 1 mL were collected, and 20 μL of each were dried under vacuum for 30 min using a SpeedVac centrifuge (Savant), resuspended in 8 μL of denaturing loading buffer (95% formamide, 0.025% xylene cyanol, 0.025% bromophenol blue), and analyzed by electrophoresis on a 12% denaturing urea-polyacrylamide gel in 1× TBE buffer. The remaining portions of each fraction were precipitated by adding 100 μL of 5 M ammonium acetate (pH 5.3) and 1 mL of isopropanol, followed by incubation overnight at -20°C. RNA was recovered by centrifugation at 16000g for 45 min at 4°C, washed twice with cold 80% ethanol, and air-dried for 20 min at 20°C. Fractions with similar electrophoretic profiles were pooled by sequentially resuspending the corresponding pellets in 15 μL of milli-Q water.

### Cyanoethylation of tRNA

Acrylonitrile treatment was performed based the optimal conditions described in (44). The reaction mixture contained 100 μL of 41% ethanol/1.1 M triethylammoniumacetate (pH 8.6), 5 - 13 μL of tRNA (∼100 μg), and 12 μL acrylonitrile (Sigma-Aldrich). The mixture was incubated at 70°C for 2 h, and the reaction was stopped by adding 12 μL 5M ammonium acetate (pH 5.3) and 300 μL of cold absolute ethanol. After incubation overnight at -20°C, the precipitated tRNA was collected by centrifugation at 16000g for 30 min at 4°C, washed with 80% ethanol, air-dried, and resuspended in 20 μL milli-Q water.

### Two-dimensional urea-polyacrylamide gel electrophoresis

The first dimension was performed on a denaturing 15% polyacrylamide gel in 1× TBE buffer containing 8.3 M urea, cast between two 30 x 40 cm glass plates with 1-mm spacers and a 15-well comb forming wells 1.5 mm deep and 0.75 mm wide. The gel was pre-run for at 30 min at 900 V using a PowerPac 3000 power supply (BioRad). Samples were prepared in a total volume of 8 μL containing 15-20 μg of tRNA in the presence of 47.5% formamide, 0.0125% xylene cyanol, and 0.0125 % bromophenol, and heated for 2 min at 95°C prior to loading. Electrophoresis was carried out in 1× TBE running buffer at 13 W for 24 h at 20°C using an S2 sequencing gel electrophoresis apparatus (BR Life Technologies). The gel was stained with ethidium bromide, visualized under UV light using a BenchTop UVP transilluminator (Fisher scientific), and the lane containing tRNA was excised and soaked in water. The second dimension was performed using a semidenaturing 20% polyacrylamide gel in 1X TBE buffer containing 4.15 M urea, cast between two 16 x 20 cm glass plates compatible with the Protean II xi Cell electrophoresis apparatus (BioRad). The excised 1D lane was placed horizontally at the top and between the two plates before pouring the gel solution to embed it after polymerization. Electrophoresis was then performed in 1× TBE running buffer at 6W/gel for 24 h at 20°C. The gel was visualized under UV light, photographed using a Gel Doc EZ imager (BioRad), and the individual spots were excised, transferred into 1.5 mL tubes, dried for 30 min at 20°C in SpeedVac centrifuge (Savant), and stored at -20°C.

### Purification of individual tRNAs

Individual *S. aureus* tRNAs were purified using specific biotinylated probes (Table S1). tRNA^Lys^(UUU) and tRNA^Thr^(UGU) were each purified from 750 μg of bulk tRNA isolated from the USA300 JE2 strain. Proteogenic tRNA^Gly^(UCC) P1, as well as non-proteogenic tRNA^Gly^(UCC) NP1+NP2 mixture and NP3, were purified from 2000 μg of bulk tRNA isolated from an RN4220 strain lacking one gene copy of proteogenic tRNA^Gly^(GCC) P1 (40). Bulk tRNA was prepared by DEAE anion-exchange chromatography and dissolved in buffer 10 mM Tris-HCl (pH 7.5), 0.9 M tetraethylammonium chloride, and 0.1 mM EDTA. For preparation of the affinity column, 200 μL of streptavidin sepharose resin (Cytiva) were washed with water (10 × 200 μL), equilibrated in 10 mM Tris-HCl (pH 7.5) and 150 mM NaCl (10 × 200 μL), and incubated with 100 μL of 100 μM biotinylated probe for 30 min at 37°C under agitation. Following washing with 10 mM Tris-HCl (pH 7.5) (10 × 200 μL), the tRNA sample was applied to the resin and incubated at 37°C for 1 h under agitation for tRNA^Lys^(UUU) and tRNA^Thr^(UGU), or overnight with continuous recirculation using a peristaltic pump for all tRNA^Gly^(UCC) P1, NP1+NP2, and NP3. The resin was washed with 10 mM Tris-HCl (pH 7.5) (4 × 500 μL) and tRNAs were eluted four times with 200 μL of the same buffer preheated to 65°C, incubating for 5 min each time. Fractions were pooled, ethanol-precipitated and resuspended in water.

### LC-MS/MS oligonucleotide mass spectrometry

Gel spots were digested following different protocol depending on the RNase used. For RNase T1 digestions, 50 μL of 100 mM ammonium acetate (pH 6.8) containing 1U/μL RNase T1 (Thermo Fischer Scientific) were added to each spot and incubated at 50°C for 3 h, then at 37°C under agitation (300 rpm) for 9 h. For RNase A digestions, the same conditions were used, except that 0.01 μg/μL RNase A (Thermo Fischer Scientific) was used. For RNase U2 digestions, 45 μL of 250 mM ammonium acetate (pH 4.7) were added to each spot and incubated at 50°C for 1 h, after which 5 μL of RNase U2 (0.01 mg/mL in the same buffer) was added, and incubation was continued for 30 min. Digestion in solution was performed in a final volume of 20 μL containing 100 ng - 500 ng of tRNA under the conditions described above.

Samples were incubated at 95°C for 2 min, and on ice for 3 min prior to desalting using ZipTip C18 pipette tips (0.2 μL resin bed, Merck Millipore). The resin was activated wit 50% acetonitrile (5 × 10 μL), washed with milli-Q water (5 × 10 μL), and equilibrated in 200 mM ammonium acetate (pH 6.8). The digestion reaction was passed 30 times through the ZipTip, the resin was sequentially washed with 200 mM ammonium acetate (10 × 10 μL) and milli-Q water (10 × 10 μL), and the bound RNase-derived oligonucleotides were finally eluted with 50% acrylonitrile (2 × 10 μL) into an HPLC vial insert (Fisher scientific). The vial was placed into a 1.5 mL tube, and the solution was dried for 30 min at 20°C in SpeedVac centrifuge (Savant). The pellet was resuspended in 3 μL milli-Q water and injected onto an Acquity UPLC peptide BEH C18 column (130 Å, 1.7 μm, 75 μm × 200 mm) equilibrated with buffer containing 7.5 mM triethylammonium acetate, 7 mM triethylamine, and 200 mM hexafluoroisopropanol using a nanoAcquity UPLC system (Waters). Oligonucleotides were eluted at 300 nL/min, first with a 15-35% methanol gradient over 2 min, followed by a 35 - 50% methanol gradient over 20 min. Tandem mass spectrometry (MS/MS) was performed using a SYNAPT G2-S instrument (Waters) equipped with a NanoLockSpray-ESI source operated in negative mode, with a capillary voltage set to 2.6 kV and a sample cone of 30 V. The source temperature was set to 130°C. MS analysis was performed over an *m/z* range from 500-15000, followed by fast data-dependent acquisition (FastDDA) of MS/MS scans using collision-induced dissociation (CID) for fragmentation. MS/MS data were processed with MassLynx 4.1 (Waters). Multi-charged CID spectra were deconvoluted into single-charged spectra using the MaxEnt3 tool and manually sequenced by following the typical fragmentation ion series (138). Correct spectra interpretation was verified using the MongoOligo online calculator (https://mstoolbox.github.io/). A mass difference threshold of 0.05 Da between observed and theoretical parent ion mass values was applied for reliable identification.

### LC-MS/MS nucleoside mass spectrometry

Digestion to nucleosides was performed for 2 h at 37°C in 20-μL reactions containing 0.6 U nuclease P1 from *Penicillium citrinum* (Sigma-Aldrich), 0.2 U snake venom phosphodiesterase from *Crotalus adamanteus* (Worthington), 10 U benzonase (Sigma-Aldrich), 0.2 U bovine intestin phosphatase (Sigma-Aldrich), and 200 ng pentostatin (Sigma-Aldrich) in 5 mM Tris-HCl pH 8 and 1 mM MgCl_2_. LC-MS/MS analysis was carried out on an Agilent 1260 LC series chromatography system coupled to an Agilent 6460 Triple Quadrupole mass spectrometer equipped with an electrospray ion source (ESI). Nucleosides were separated at 35°C on a Synergi Fusion RP18 column (80 Å, 4 μm, 2.0 mm × 250 mm; Phenomenex) equilibrated in buffer A (5 mM ammonium acetate pH 5.3) and eluted with buffer B (acetonitrile) at 0.35 mL/min. For analysis of total tRNA, 6 μg were digested and 1 μg of the hydrolysate was spiked with 50 ng of *E. coli* ^13^C-labeled RNA prior to chromatographic separation using a linear gradient of buffer B from 0 to 10% over 20 min, 10 to 25% over 10 min, 25 to 80% over 10 min, 80 to 0% over 3 min, and 100% buffer A for 11 min. For analysis of purified tRNAs Gly, 450 ng were digested and 400 ng of each hydrolysate was spiked with 50 ng of yeast ^13^C-labeled RNA prior to separation with a linear gradient of buffer B from 0 to 8% over 10 min, from 8 to 40% over 10 min, 40 to 0% over 3 min, and 100% buffer A for 7 min. The UV signal at 254 nm was recorded using a diode array detector. The ESI parameters were the following: gas temperature 350°C, gas flow 8 L/min, nebulizer pressure 50 psi, sheath gas temperature 350°C, sheath gas flow 12 L/min, capillary voltage 3000 V, and nozzle voltage 0 V. The mass spectrometer was run in positive mode, and dynamic multiple reaction monitoring (dMRM) was performed using Agilent MassHunter software. Absolute quantification was carried out using external calibrations with synthetic standards and internal calibration with ^13^C-labeled RNA spiked into the samples as described in (139). Raw data were converted to open mzML format using ProteoWizard MSconverter and the resulting files were processed using custom Python scripts employing Pyteomics. The area under the curve (AUC) for each each ^12^C and ^13^C nucleoside was computed and the ratio AUC(^12^C)/AUC(^13^C) was used to generate calibration curves. The tRNA amount was estimated from the UV signal of adenosine, assuming an average of 16 adenosine residues per tRNA for analysis of total tRNA. Relevant parameters for dMRM acquisition, sample lists, AUC values, and quantification results are provided in Supplementary Data 3 for analysis of total tRNA, and in Supplementary Data 4 for analysis of purified tRNA^Gly^.

### Analysis of tRNA modification by next generation sequencing

#### RiboMethSeq

Detection of 2’-O-ribomethylations (Nm) by RiboMethSeq is based on the resistance of the phosphodiester bond following the Nm residue to alkaline hydrolysis. The analysis was performed essentially as previously described (47). Briefly, total tRNA (150 ng) was subjected to alkaline hydrolysis (pH 9.4) followed by ethanol precipitation. Fragments were end-repaired and ligated to adapters using the NEBNext Small RNA kir for Illumina. Sequencing was performed on a NextSeq 2000 instrument. After removal of adapter sequences by trimmomatic v0.39, reads were mapped to *S. aureus* HG001 tRNA sequences using bowtie2 in end-to-end mode. Reads’ 5’ and 3’-ends were counted and the different RiboMethSeq score were calculated (53). Only MethScore values associated to ScoreMEAN > 0.92 and ScoreA > 0.5 were considered relevant.

#### AlkAnilineSeq

Detection of m^7^G and D by AlkAnilineSeq relies on aniline-mediated scission at the abasic site that arise from nucleobase loss following cleavage of the fragile N-glycosidic under alkaline, high-temperature conditions. The analysis was performed as previously described (49). Briefly, tRNA (100 ng) was subjected to fragmentation under mild alkaline conditions (5 min, 96°C) followed by ethanol precipitation. Fragments were end-repaired and then subjected to aniline treatment, resulting in deprotection of a 5’-phosphate at the N+1 nucleotide, which serves for selective ligation of sequencing adapters. RNA fragments were converted to libraries and sequenced as described above. Trimmed reads were aligned to tRNA sequences and the reads’ 5’-ends were counted after applying a -1 shift. Two scores were used for data analysis: normalized cleavage score, which represents the 1000x proportion of reads starting at a given position to the total number of reads mapped to the corresponding tRNA sequence, and stop ratio, which is calculated as the ratio of reads starting at a given position to the total number of reads cover it.

#### HydraPsiSeq

Detection of Ψ and 5-substituted U by HydraPsiSeq is based on their resistance to hydrazine cleavage, while unmodified U are efficiently cleaved. The analysis was performed as previously described (52). In brief, tRNA (200 ng) was subjected to hydrazine cleavage, followed by aniline cleavage of the phosphodiester bond at all unmodified U. Fragments were converted to libraries and sequenced as described above. After trimming, reads were mapped on the tRNA sequences and HydraPsiSeq scores were calculated from the normalized U-profile. Only Ψ-Score values associated to ScoreMEAN > 0.92 and ScoreA > 0.5 were considered relevant.

#### BIDseq

Detection of Ψ by BIDseq relies on the deletion signature induced by a bisulfite-Ψ adduct. This analysis was performed following an adapted version of the original BIDseq protocol (54). Briefly, tRNA (100 ng) were subjected to fragmentation followed by bisulfite treatment and desulphonation as described (140). Treated and untreated RNAs were converted to libraries and sequenced as described above. After trimming, reads were mapped on the tRNA sequences allowing to retain 1-3 nt gapped reads. Further analysis was performed using samtools’s mpileup utility to count the deletion score at every position in the reference.

#### GLORIseq

Detection of N^6^-substituted A is based on selective deamination of unmodified A, which is performed on an RNA sample treated with glyoxal to protect G from nitrite-induced deamination. This is followed by deprotection and reverse transcription to convert RNA into cDNA. The conditions used were as described in (50). In brief, tRNA (100 ng) was subjected to glyoxal treatment (30 min, 50°C) followed by boric acid treatment (30 min, 50°C). The protected RNA was then subjected to nitrite-mediated deamination, deprotected in 500 mM TEAA/formamide buffer, and ethanol precipitated. Fragments were end-repaired, converted to libraries and sequenced as described above. Trimmed reads were mapped to the A → G converted tRNA sequences allowing 1 nt mismatch. Further analysis was done with samtools’s mpileup utility to count the mismatch profile at every position in the reference. GtoA mismatches correspond to deamination-resistant modified residues (*e.g.* m^6^A, t^6^A, i^6^A). The score value correspond to the molar ratio of the modified A.

### Dihydrouridine quantification

Dihydrouridine was quantified using a colorimetric assay based on the detection of the ureido group generated by ring-opening upon alkali treatment (141). The assay was performed following the protocol described in (93). Bulk tRNA was prepared by DEAE anion-exchange chromatography as described above. A total of 100 μg of tRNA in 50 μL milli-Q water were combined with 5 μL 1 N KOH and incubated at 37°C for 30 min. The solution was then acidified with 25 μL of concentrated H_2_SO_4_, followed by the addition of 50 μL of a freshly prepared 1:1 mixture of 2 mg/mL N-phenyl-*p*-phenylendiamine and 3% (m/v) 2, 3-butanedione monoxime as described in (142). Samples were incubated at 95°C for 10 min, then at 50°C for 10 min, after which 50 μL of 1 mM FeCl_3_ in concentrated H_2_SO_4_ was immediately added. The reactions were transferred into a 96-well plate, and the absorbances at 550 nm and 465 nm were measured using a Multiskan SkyHigh microplate reader (Thermo Fischer). The difference between these two readings was correlated with dihydrouridine concentration using a standard curve prepared from multiple concentrations (up to 250 μM) of dihydrouracil (Thermo Fischer).

### tRNA dihydrouridylation assay

Synthetic tRNA^Gly^(UCC) NP1 and NP2 were produced as described in (96). Purification of recombinant *S. aureus* DusB2 is described in supplementary methods. *In vitro* dihydrouridylation was performed under aerobic conditions in a 50-μL reaction containing 8 μM tRNA and 2 μM DusB2 in 50 mM HEPES-KOH pH 7.5, 150 mM NaCl, 10 mM MgCl_2_, 5 mM DTT, and 15 % glycerol. The reaction was initiated by adding 2 μL of NADPH 50 mM and incubated for 1 h at 37°C. Quenching was performed by sequential extraction with acidic PCI and chloroform, followed by ethanol precipitation. Recovered tRNA was digested in solution with RNase U2 and analyzed by UPLC-MS/MS as described in the main text.

### Protein mass spectrometry

Time course-proteomics analysis was performed in four biological replicates of *S. aureus* grown in rich medium. Overnight cultures were diluted to an OD_600_ of 0.05 in 50 mL of BHI (250-mL flasks) and incubated at 37°C with shaking at 180 rpm. Culture volumes corresponding to 10 OD_600_ units were collected at 2, 3, 4 and 4.5 h by centrifugation at 2100g for 10 min at 4°C. Bacterial pellets were resuspended in 400 μL buffer UCTF (7 M urea, 4% CHAPS, 2 M thiourea, 20 mM Tris-HCl pH 8.0), transferred into 2 mL tubes containing 0.6 g Lysis Matrix B (MP Biomedicals) and lysed mechanically for 40 s at 6 m/s using a FastPrep-24 system (MP Biomedicals). Tubes were centrifuged at 16000g for 10 min at 4°C to remove beads and the supernatant was recovered and stored -20°C. For mass spectrometry analysis, 10 μg of *S. aureus* proteins were precipitated twice with 0.1 M ammonium acetate in 100% methanol (5 volumes, - 20°C) to get rid of detergents and other contaminants. Proteins (300 ng) were then reduced with 5 M dithiothreitol for 10 min at 95°C, alkylated with 10 mM Iodoacetamide for 20 min at 20°C, and digested overnight with 1:50 (v:v) of sequencing-grade trypsin (Promega). Peptides (200 ng) were analyzed by nanoLC-MS/MS on a reversed phase nanoElute2 coupled to a TIMS-TOF Pro 2 mass spectrometer (Bruker Daltonik Gmbh) using a data-dependent acquisition strategy. Peptides were further separated on the integrated emitter column IonOpticks Aurora Elite (15 cm × 75 μm, 1.7 μm particle size, and 120 Å pore size; AUR3-15075C18-CSI). Data were searched against a homemade *S. aureus* HG001 protein database with Mascot algorithm (version 2.8, Matrix Science) and proteins were aligned and quantified with label-free Proline 2.1 software (exhaustive alignment with 40 s cross-assignment within groups) (143). Prostar 1.32.3 was used for statistical analysis of the intensities. After a 0.15 quantile centering normalization, partially observed values and values missing in the entire condition were imputed with det quantile 1%. A LIMMA statistical test with a logFC threshold of 1 and a Benjamini-Hochberg correction were used to generate log_2_FC and adjusted *P-values* for comparison among conditions B vs A, C vs B, D vs C, and D vs A. For comparative analysis of expression trends, the abundance of each protein was scaled based on the highest (1.0) and lowest (0.0) values to obtain relative abundances. The averaged relative abundance of each condition was then normalized to the first condition (1.0) to assess positive (> 1.0) or negative (< 1.0) trends across conditions.

### Nanopore tRNA sequencing

The relative abundance of the *S. aureus* tRNA species was estimated by Nanopore tRNA sequencing of bulk tRNA samples purified as described above from cultures at late exponential phase (OD_600_ 2.5 – 3.0) in three biological replicates. First, 3 μg of tRNA were polyadenylated in a 20 μL reaction containing 0.25 U of E. coli Poly(A) polymerase (NEB), 1 mM ATP, 20 U RNase inhibitor (Promega), 50 mM Tris-HCl pH 8.1, 250 mM NaCl and 10 mM MgCl_2_, for 30 min at 37°C. The reaction was purified with 1.6 volumes of SPRI beads (Mag-Bind TotalPure NGS, Omega), washed twice with 200 μL fresh 80 % EtOH, and eluted in 10 μL RNase-free H2O. Polyadenylated tRNA were then barcoded following the WarpDemux strategy (144). Custom RTA adapters (10 μM forward, 10 μM reverse) were annealed in 30 mM HEPES-KOH (pH 7.5), 100 mM K-Acetate by incubation for 1 min at 95 °C followed by cooling to 25°C at a rate of 0.5 °C/min. The annealed RTA were then diluted to 0.7 μM and stored at -20°C until use. For each sample, 100 ng of polyA-selected RNA were ligated to 1 μL of 0.7 μM annealed RTA in the presence of 2 μL 5x NEBNext Quick Ligation Buffer (NEB), 0.5 μL T4 DNA ligase high concentration (NEB) and 0.2 μL RNAsin (Promega) in a final volume of 10 μL by incubation at 22°C for 15 min. To stop the ligation, 2 μL of 0.5 M EDTA was added and samples were pooled and purified with 0.5 volumes of SPRI beads (Mag-Bind TotalPure NGS, Omega Bio-tek), washed twice with 200 μL fresh 80 % EtOH, and eluted in 10 μL RNase-free H_2_O. Reverse transcription was performed by adding 8 μL of 5X Induro buffer, 2 μL of 10 mM dNTP, 2 μL of 10 μM random hexamer oligonucleotides, 2 μL Induro RT (NEB), 0.5 μL RNasin in a final volume of 40 μL and incubation 2 min at 22°C (annealing), 15 min at 60°C (reverse transcription) and 10 min at 70°C (heat-inactivation). The reaction was purified with 1.4 volumes of SPRI beads, washed twice with 200 μL fresh 80 % EtOH, and eluted in 10 μL RNase-free H2O. The sequencing adapter ligation was realized using the SQK-RNA004 library kit (ONT) according to the manufacturer’s instruction. Briefly, 10 μL purified library was mixed with 6 μL 5x NEBNext Quick Ligation Buffer, 5 μL RLA sequencing adapter (ONT SQK-RNA004) and 3 μL T4 DNA Ligase high concentration in a final volume of 30 μL and incubated 15 min at 22°C. The reaction was purified with 0.5 volumes of SPRI beads, washed twice with 100 μL RNA wash buffer (WSB, ONT SQK-RNA004). The library was then eluted in 32 μL RNA elution buffer (REB, ONT SQK-RNA004). The library was loaded onto a Promethion (FLO-PRO004RA) flow cell for sequencing. Data acquisition was performed using MinKNOW version 24.11. Basecalling was performed with dorado v0.9.1 using the model parameters rna004_130bps_sup@v5.1.0. Demultiplexing was performed using the demux command of warpdemux v0.4.4. Following basecalling, reads were processed to remove polyA using cutadapt v5.0 with the parameters cutadapt -a A{10} –times 1 -e 0.1 -m 20 –overlap 10, then aligned to tRNA references using bwa (v0.7.18) with the parameters bwa mem -W13 -k6 xont2d -T20. Reads corresponding to isodecoders were merged. For each tRNA in each sample, read counts were normalized by the total number of reads to obtain relative abundance.

### Ribosome profiling

Ribosome profiling was performed in two biological replicates of *S. aureus* HG001 grown in rich medium to late exponential phase. Overnight cultures were diluted to an OD_600_ of 0.05 in 50 mL of BHI (500-mL flasks) and incubated at 37°C with shaking at 180 rpm for 3.6 h, reaching a final OD_600_ of ≈ 2.6. Cultures were rapidly cooled by swirling in an ice bath for 3 min, divided into 40-mL and 10-mL fractions, and centrifuged at 2100g for 5 min at 4°C. The 10-mL pellets were frozen in liquid nitrogen and used for total RNA sequencing, as previously described (98). The 40-mL pellets were resuspended in 550 μL of cold lysis buffer (20 Mm Tris-HCl pH 8.0, 50 mM MgCl_2_, 100 mM NH_4_Cl, 0.4% Triton X-100, 0.1% Nonidet P-40, 0.1 U/μL DNase I, 1 mM GMPPNP), transferred to 2-mL tubes containing Lysis Matrix B (MP Biomedicals), and mechanically lysed in FastPrep-24 system (MP Biomedicals) for 40 s at 6 m/s. Following centrifugation at 16000g for 10 min at 4°C, the supernatants containing polysomes were recovered, and their absorbance at 260 nm (10-mm path) was measured using a NanoDrop ND-1000 spectrophotometer (Thermo Fisher Scientific). Ribosome-protected fragments (RPFs) were generated by treating 40, 000 AU of polysomes with 250 U of RNase If (NEB) for 20 min at 37°C under agitation at 800 rpm. The reaction was stopped by adding 100 U of SUPERase-In (Thermo Fisher Scientific) prior to separation on an 11-mL 5-50% sucrose gradient in buffer G (20 Mm Tris-HCl pH 8.0, 150 mM MgCl_2_, 100 mM NH_4_Cl, 2 mM DTT) at 39, 000g for 2h 46 min at 4°C. Monosomes were isolated using a piston gradient fractionator (Biocomp) and the RPFs were extracted using hot phenol-chloroform and size-selected on a denaturing 15%-polyacrylamide/8M-urea gel as previously described (98).

Total RNA and RPF samples were RNA-depleted using a Ribo-Seq ribo-POOL kit adapted to *S. aureus* (siTOOLs Biotech). RPF samples were dephosphorylated with Antarctic phosphatase (NEB) and rephosphorylated with T4 polynucleotide kinase (NEB) prior to library preparation using the NEBNext Small RNA Library Prep Set for Illumina (NEB). For total RNA samples, libraries were prepared using the NEBNext Ultra II Directional RNA Library Prep Kit for Illumina after RNA fragmentation in 50 mM bicarbonate buffer at 96°C for 8 min. Libraries were sequenced on an Illumina NextSeq 1000 instrument and the demultiplexed FASTQ sequencing files were processed as previously described (98), using published and publicly available workflows (145). Raw reads in each sample were converted to normalized RPKM and TPM values, which were then used to compare the relative abundance of the tRNA-modifying enzymes. For pausing analysis, only 28-nucleotides length reads were considered, using -13, -16, and -19 shifts for codons in the A, P, and E-sites respectively.

## DATA AVAILABILITY

Mass spectrometry data are available in the PRIDE database, under the accession number XXXX for RNA samples and XXXX for protein samples. Deep sequencing data are available in ENA (The European Bioinformatics Institute EMBL-EBI) under the accession number XXXX for RNA modification analysis and XXXX for ribosome profiling.

## SUPPLEMENTARY DATA

Supplementary methods, figures, and data are available.

## AKCNOWLEDGMENTS

We thank Constantinos Stathopoulos for providing templates for *in vitro* transcription of non-proteogenic tRNA Gly and the edited *S. aureus* RN4220 strain, Patick Limbach for providing the plasmid carrying the RNase U2 coding sequence, Valéry de Crécy-Lagard for helpful discussions and sharing unpublished data, Antony Lechner for RNA mass spectrometry input, Roberto Bahena-Ceron for helpful discussions, and Laura Antoine for setting the foundation of this study.

## REFERENCES

1. Shepherd, J. and Ibba, M. (2015) Bacterial transfer RNAs. FEMS Microbiol Rev, 39, 280–300.

2. Dana, A. and Tuller, T. (2014) The effect of tRNA levels on decoding times of mRNA codons. Nucleic Acids Res, 42, 9171–9181.

3. Novoa, E.M. and Ribas de Pouplana, L. (2012) Speeding with control: codon usage, tRNAs, and ribosomes. Trends in Genetics, 28, 574–581.

4. Samatova, E., Daberger, J., Liutkute, M. and Rodnina, M.V. (2021) Translational Control by Ribosome Pausing in Bacteria: How a Non-uniform Pace of Translation Affects Protein Production and Folding. Front. Microbiol., 11.

5. Zhang, M. and Lu, Z. (2025) tRNA modifications: greasing the wheels of translation and beyond. RNA Biology, 22, 1–25.

6. Zhang, W., Foo, M., Eren, A.M. and Pan, T. (2022) tRNA modification dynamics from individual organisms to metaepitranscriptomics of microbiomes. Mol Cell, 82, 891– 906.

7. Quax, T.E.F., Claassens, N.J., Söll, D. and van der Oost, J. (2015) Codon Bias as a Means to Fine-Tune Gene Expression. Mol Cell, 59, 149–161.

8. Avcilar-Kucukgoze, I., Bartholomäus, A., Cordero Varela, J.A., Kaml, R.F.-X., Neubauer, P., Budisa, N. and Ignatova, Z. (2016) Discharging tRNAs: a tug of war between translation and detoxification in Escherichia coli. Nucleic Acids Res, 44, 8324–8334.

9. Lyons, S.M., Fay, M.M. and Ivanov, P. (2018) The role of RNA modifications in the regulation of tRNA cleavage. FEBS Letters, 592, 2828–2844.

10. Peschek, J. and Tuorto, F. (2025) Interplay Between tRNA Modifications and Processing. Journal of Molecular Biology, 437, 169198.

11. Fruchard, L., Salinas, C., Carvalho, A. and Baharoglu, Z. tRNA-modifying enzymes in bacterial stress adaptation. Open Biol, 15, 250194.

12. McCown, P.J., Ruszkowska, A., Kunkler, C.N., Breger, K., Hulewicz, J.P., Wang, M.C., Springer, N.A. and Brown, J.A. (2020) Naturally occurring modified ribonucleosides. Wiley Interdiscip Rev RNA, 11, e1595.

13. Machnicka, M.A., Olchowik, A., Grosjean, H. and Bujnicki, J.M. (2015) Distribution and frequencies of post-transcriptional modifications in tRNAs. RNA Biol, 11, 1619– 1629.

14. Schultz, S.K. and Kothe, U. (2024) RNA modifying enzymes shape tRNA biogenesis and function. J Biol Chem, 300, 107488.

15. Ontiveros, R.J., Stoute, J. and Liu, K.F. (2019) The chemical diversity of RNA modifications. Biochem J, 476, 1227–1245.

16. Agris, P.F., Eruysal, E.R., Narendran, A., Väre, V.Y.P., Vangaveti, S. and Ranganathan, S.V. (2018) Celebrating wobble decoding: Half a century and still much is new. RNA Biology, 15, 537–553.

17. Jenner, L.B., Demeshkina, N., Yusupova, G. and Yusupov, M. (2010) Structural aspects of messenger RNA reading frame maintenance by the ribosome. Nat Struct Mol Biol, 17, 555–560.

18. Yared, M.-J., Marcelot, A., Barraud, P., Yared, M.-J., Marcelot, A. and Barraud, P. (2024) Beyond the Anticodon: tRNA Core Modifications and Their Impact on Structure, Translation and Stress Adaptation. Genes, 15.

19. Schultz, S.K., Katanski, C.D., Halucha, M., Peña, N., Fahlman, R.P., Pan, T. and Kothe, U. (2024) Modifications in the T arm of tRNA globally determine tRNA maturation, function, and cellular fitness. Proceedings of the National Academy of Sciences, 121, e2401154121.

20. McKenney, K.M., Rubio, M.A.T. and Alfonzo, J.D. (2017) The Evolution of Substrate Specificity by tRNA Modification Enzymes. Enzymes, 41, 51–88.

21. de Crécy-Lagard, V. and Jairoch, M. (2021) Functions of bacterial tRNA modifications: from ubiquity to diversity. Trends Microbiol, 29, 41–53.

22. Barraud, P. and Tisné, C. (2019) To be or not to be modified: Miscellaneous aspects influencing nucleotide modifications in tRNAs. IUBMB Life, 71, 1126–1140.

23. Kasprzak, J.M., Czerwoniec, A. and Bujnicki, J.M. (2012) Molecular evolution of dihydrouridine synthases. BMC Bioinformatics, 13, 153.

24. Bou-Nader, C., Montémont, H., Guérineau, V., Jean-Jean, O., Brégeon, D. and Hamdane, D. (2018) Unveiling structural and functional divergences of bacterial tRNA dihydrouridine synthases: perspectives on the evolution scenario. Nucleic Acids Res, 46, 1386–1394.

25. Sudol, C., Kilz, L.-M., Marchand, V., Thuillier, Q., Guérineau, V., Goyenvalle, C., Faivre, B., Toubdji, S., Lombard, M., Jean-Jean, O., et al. (2024) Functional redundancy in tRNA dihydrouridylation. Nucleic Acids Res, 52, 5880–5894.

26. Benítez-Páez, A., Villarroya, M. and Armengod, M.-E. (2012) The Escherichia coli RlmN methyltransferase is a dual-specificity enzyme that modifies both rRNA and tRNA and controls translational accuracy. RNA, 18, 1783–1795.

27. Crécy-Lagard, V. de, Ross, R.L., Jaroch, M., Marchand, V., Eisenhart, C., Brégeon, D., Motorin, Y., Limbach, P.A., Crécy-Lagard, V. de, Ross, R.L., et al. (2020) Survey and Validation of tRNA Modifications and Their Corresponding Genes in Bacillus subtilis sp Subtilis Strain 168. Biomolecules, 10.

28. Hardy, L., Marchand, V., Bourguignon, V., Thuillier, Q., Dias, C., Krin, E., Fruchard, L., Yaacov, D.B., Mazel, D., Motorin, Y., et al. (2025) The tRNA epitranscriptomic landscape and RNA modification enzymes in Vibrio cholerae. 10.1101/2025.10.15.682536.

29. Kimura, S., Dedon, P.C. and Waldor, M.K. (2020) Comparative tRNA sequencing and RNA mass spectrometry for surveying tRNA modifications. Nat Chem Biol, 16, 964–972.

30. Tomasi, F.G., Kimura, S., Rubin, E.J. and Waldor, M.K. (2023) A tRNA modification in Mycobacterium tuberculosis facilitates optimal intracellular growth. eLife, 12, RP87146.

31. Koshla, O., Vogt, L.-M., Rydkin, O., Sehin, Y., Ostash, I., Helm, M. and Ostash, B. (2022) Landscape of Post-Transcriptional tRNA Modifications in Streptomyces albidoflavus J1074 as Portrayed by Mass Spectrometry and Genomic Data Mining. Journal of Bacteriology, 205, e00294–22.

32. Quaiyum, S., Sun, J., Marchand, V., Sun, G., Reed, C.J., Motorin, Y., Dedon, P.C., Minnick, M.F. and de Crécy-Lagard, V. (2024) Mapping the tRNA modification landscape of Bartonella henselae Houston I and Bartonella quintana Toulouse. Front. Microbiol., 15.

33. Sun, J., Wu, J., Yuan, Y., Fan, L., Chua, W.L.P., Ling, Y.H.S., Balamkundu, S., Dwijapriya, Chay, H.S.S., Begley, T.J., et al. (2025) tRNA modification profiling reveals epitranscriptome regulatory networks in Pseudomonas aeruginosa. Nucleic Acids Res, 53, gkaf696.

34. Tsui, H.-C.T., Chan, C.-K., Yuan, Y., Elias, R., Sun, J., Marchand, V., Jaroch, M., Sun, G., Manzoor, I., Kutchuashvili, A., et al. (2025) tRNA Modification Landscapes in Streptococci: Shared Losses and Clade-Specific Adaptations. 10.1101/2025.10.13.681863.

35. Andachi, Y., Yamao, F., Muto, A. and Osawa, S. (1989) Codon recognition patterns as deduced from sequences of the complete set of transfer RNA species in *Mycoplasma capricolum*. Journal of Molecular Biology, 209, 37–54.

36. Antoine, L., Wolf, P., Westhof, E., Romby, P. and Marzi, S. (2019) Mapping post-transcriptional modifications in *Staphylococcus aureus* tRNAs by nanoLC/MSMS. Biochimie, 164, 60–69.

37. Chan, P.P. and Lowe, T.M. (2016) GtRNAdb 2.0: an expanded database of transfer RNA genes identified in complete and draft genomes. Nucleic Acids Res, 44, D184– D189.

38. Giannouli, S., Kyritsis, A., Malissovas, N., Becker, H.D. and Stathopoulos, C. (2009) On the role of an unusual tRNAGly isoacceptor in *Staphylococcus aureus*. Biochimie, 91, 344–351.

39. Rietmeyer, L., Fix-Boulier, N., Le Fournis, C., Iannazzo, L., Kitoun, C., Patin, D., Mengin-Lecreulx, D., Ethève-Quelquejeu, M., Arthur, M. and Fonvielle, M. (2021) Partition of tRNAGly isoacceptors between protein and cell-wall peptidoglycan synthesis in Staphylococcus aureus. Nucleic Acids Res, 49, 684–699.

40. Kouvela, A., Jaramillo Ponce, J.R., Giarimoglou, N., Chicher, J., Marzi, S., Stathopoulos, C. and Stamatopoulou, V. (2025) Coupling tRNAGly gene redundancy with staphylococcal cell wall integrity, antibiotic susceptibility, and virulence potential. Nucleic Acids Res, 53, gkaf599.

41. Rohrer, S. and Berger-Bächi, B. (2003) FemABX Peptidyl Transferases: a Link between Branched-Chain Cell Wall Peptide Formation and β-Lactam Resistance in Gram-Positive Cocci. Antimicrobial Agents and Chemotherapy, 47, 837–846.

42. Antoine, L. and Wolf, P. (2020) Mapping of Posttranscriptional tRNA Modifications by Two-Dimensional Gel Electrophoresis Mass Spectrometry. In Arluison, V., Wien, F. (eds), RNA Spectroscopy: Methods and Protocols. Springer US, New York, NY, pp. 101–110.

43. Lechner, A. and Wolf, P. (2024) In-Gel Cyanoethylation for Pseudouridines Mass Spectrometry Detection of Bacterial Regulatory RNA. In Arluison, V., Valverde, C. (eds), Bacterial Regulatory RNA: Methods and Protocols. Springer US, New York, NY, pp. 273–287.

44. Mengel-Jørgensen, J. and Kirpekar, F. (2002) Detection of pseudouridine and other modifications in tRNA by cyanoethylation and MALDI mass spectrometry. Nucleic Acids Res, 30, e135.

45. Favre, A., Michelson, A.M. and Yaniv, M. (1971) Photochemistry of 4-thiouridine in *Escherichia coli* transfer RNA1Val. Journal of Molecular Biology, 58, 367–379.

46. Houser, W.M., Butterer, A., Addepalli, B. and Limbach, P.A. (2015) Combining recombinant ribonuclease U2 and protein phosphatase for RNA modification mapping by liquid chromatography–mass spectrometry. Analytical Biochemistry, 478, 52–58.

47. Marchand, V., Blanloeil-Oillo, F., Helm, M. and Motorin, Y. (2016) Illumina-based RiboMethSeq approach for mapping of 2′-O-Me residues in RNA. Nucleic Acids Res, 44, e135.

48. Marchand, V., Ayadi, L., Bourguignon-Igel, V., Helm, M. and Motorin, Y. (2021) AlkAniline-Seq: A Highly Sensitive and Specific Method for Simultaneous Mapping of 7-Methyl-guanosine (m7G) and 3-Methyl-cytosine (m3C) in RNAs by High-Throughput Sequencing. In McMahon, M. (ed), RNA Modifications: Methods and Protocols. Springer US, New York, NY, pp. 77–95.

49. Marchand, V., Ayadi, L., Ernst, F.G.M., Hertler, J., Bourguignon-Igel, V., Galvanin, A., Kotter, A., Helm, M., Lafontaine, D.L.J. and Motorin, Y. (2018) AlkAniline-Seq: Profiling of m7G and m3C RNA Modifications at Single Nucleotide Resolution. Angewandte Chemie International Edition, 57, 16785–16790.

50. Shen, W., Sun, H., Liu, C., Yi, Y., Hou, Y., Xiao, Y., Hu, Y., Lu, B., Peng, J., Wang, J., et al. (2024) GLORI for absolute quantification of transcriptome-wide m6A at single-base resolution. Nat Protoc, 19, 1252–1287.

51. Liu, H., Zeng, T., He, C., Rawal, V.H., Zhou, H. and Dickinson, B.C. (2022) Development of mild chemical catalysis conditions for m1A-to-m6A rearrangement on RNA. ACS Chem Biol, 17, 1334–1342.

52. Marchand, V., Pichot, F., Neybecker, P., Ayadi, L., Bourguignon-Igel, V., Wacheul, L., Lafontaine, D.L.J., Pinzano, A., Helm, M. and Motorin, Y. (2020) HydraPsiSeq: a method for systematic and quantitative mapping of pseudouridines in RNA. Nucleic Acids Res, 48, e110.

53. Pichot, F., Marchand, V., Ayadi, L., Bourguignon-Igel, V., Helm, M. and Motorin, Y. (2020) Holistic Optimization of Bioinformatic Analysis Pipeline for Detection and Quantification of 2′-O-Methylations in RNA by RiboMethSeq. Front. Genet., 11.

54. Dai, Q., Zhang, L.-S., Sun, H.-L., Pajdzik, K., Yang, L., Ye, C., Ju, C.-W., Liu, S., Wang, Y., Zheng, Z., et al. (2023) Quantitative sequencing using BID-seq uncovers abundant pseudouridines in mammalian mRNA at base resolution. Nat Biotechnol, 41, 344– 354.

55. van Kempen, M., Kim, S.S., Tumescheit, C., Mirdita, M., Lee, J., Gilchrist, C.L.M., Söding, J. and Steinegger, M. (2024) Fast and accurate protein structure search with Foldseek. Nat Biotechnol, 42, 243–246.

56. Boccaletto, P., Stefaniak, F., Ray, A., Cappannini, A., Mukherjee, S., Purta, E., Kurkowska, M., Shirvanizadeh, N., Destefanis, E., Groza, P., et al. (2021) MODOMICS: a database of RNA modification pathways. 2021 update. Nucleic Acids Res, 50, D231–D235.

57. Urbonavičius, J., Skouloubris, S., Myllykallio, H. and Grosjean, H. (2005) Identification of a novel gene encoding a flavin-dependent tRNA:m5U methyltransferase in bacteria—evolutionary implications. Nucleic Acids Res, 33, 3955–3964.

58. Gutgsell, N., Englund, N., Niu, L., Kaya, Y., Lane, B.G. and Ofengand, J. (2000) Deletion of the Escherichia coli pseudouridine synthase gene truB blocks formation of pseudouridine 55 in tRNA in vivo, does not affect exponential growth, but confers a strong selective disadvantage in competition with wild-type cells. RNA, 6, 1870– 1881.

59. Dégut, C., Roovers, M., Barraud, P., Brachet, F., Feller, A., Larue, V., Al Refaii, A., Caillet, J., Droogmans, L. and Tisné, C. (2019) Structural characterization of B. subtilis m1A22 tRNA methyltransferase TrmK: insights into tRNA recognition. Nucleic Acids Res, 47, 4736–4750.

60. Sweeney, P., Galliford, A., Kumar, A., Raju, D., Krishna, N.B., Sutherland, E., Leo, C.J., Fisher, G., Lalitha, R., Muthuraj, L., et al. (2022) Structure, dynamics, and molecular inhibition of the *Staphylococcus aureus* m1A22-tRNA methyltransferase TrmK. Journal of Biological Chemistry, 298, 102040.

61. Tomikawa, C. (2018) 7-Methylguanosine Modifications in Transfer RNA (tRNA). Int J Mol Sci, 19, 4080.

62. Steinberg, S., Misch, A. and Sprinzl, M. (1993) Compilation of tRNA sequences and sequences of tRNA genes. Nucleic Acids Res, 21, 3011–3015.

63. Losey, H.C., Ruthenburg, A.J. and Verdine, G.L. (2006) Crystal structure of Staphylococcus aureus tRNA adenosine deaminase TadA in complex with RNA. Nat Struct Mol Biol, 13, 153–159.

64. Suzuki, T. and Miyauchi, K. (2010) Discovery and characterization of tRNAIle lysidine synthetase (TilS). FEBS Letters, 584, 272–277.

65. Zhou, M., Long, T., Fang, Z.-P., Zhou, X.-L., Liu, R.-J. and Wang, E.-D. (2015) Identification of determinants for tRNA substrate recognition by Escherichia coli C/U34 2′-O-methyltransferase. RNA Biol, 12, 900–911.

66. Taniguchi, T., Miyauchi, K., Sakaguchi, Y., Yamashita, S., Soma, A., Tomita, K. and Suzuki, T. (2018) Acetate-dependent tRNA acetylation required for decoding fidelity in protein synthesis. Nat Chem Biol, 14, 1010–1020.

67. Sakai, Y., Kimura, S. and Suzuki, T. (2019) Dual pathways of tRNA hydroxylation ensure efficient translation by expanding decoding capability. Nat Commun, 10, 2858.

68. Sakaguchi, R., Giessing, A., Dai, Q., Lahoud, G., Liutkeviciute, Z., Klimasauskas, S., Piccirilli, J., Kirpekar, F. and Hou, Y.-M. (2012) Recognition of guanosine by dissimilar tRNA methyltransferases. RNA, 18, 1687–1701.

69. Jeong, H., Lee, Y. and Kim, J. (2022) Structural and functional characterization of TrmM in m6A modification of bacterial tRNA. Protein Sci, 31, e4319.

70. Matuszewski, M., Wojciechowski, J., Miyauchi, K., Gdaniec, Z., Wolf, W.M., Suzuki, T. and Sochacka, E. (2017) A hydantoin isoform of cyclic N6-threonylcarbamoyladenosine (ct6A) is present in tRNAs. Nucleic Acids Res, 45, 2137–2149.

71. Miyauchi, K., Kimura, S. and Suzuki, T. (2013) A cyclic form of N6-threonylcarbamoyladenosine as a widely distributed tRNA hypermodification. Nat Chem Biol, 9, 105–111.

72. Hur, S. and Stroud, R.M. (2007) How U38, 39, and 40 of Many tRNAs Become the Targets for Pseudouridylation by TruA. Molecular Cell, 26, 189–203.

73. Rajakovich, L.J., Tomlinson, J. and Dos Santos, P.C. (2012) Functional Analysis of Bacillus subtilis Genes Involved in the Biosynthesis of 4-Thiouridine in tRNA. Journal of Bacteriology, 194, 4933–4940.

74. Emilsson, V., Näslund, A.K. and Kurlad, C.G. (1992) Thiolation of transfer RNA in Escherichia coli varies with growth rate. Nucleic Acids Res, 20, 4499–4505.

75. Yoo, J., Lee, Y., Cho, G., Lim, J. and Kim, J. (2025) Structural and functional characterization of CspR, a 2′-O-methyltransferase acting on wobble position within tRNA. Nucleic Acids Res, 53, gkaf751.

76. de Crécy-Lagard, V., Hutinet, G., Cediel-Becerra, J.D.D., Yuan, Y., Zallot, R., Chevrette, M.G., Ratnayake, R.M.M.N., Jaroch, M., Quaiyum, S. and Bruner, S. (2024) Biosynthesis and function of 7-deazaguanine derivatives in bacteria and phages. Microbiology and Molecular Biology Reviews, 88, e00199–23.

77. Addepalli, B. and Limbach, P.A. (2016) Pseudouridine in the Anticodon of *Escherichia coli* tRNATyr(QΨA) Is Catalyzed by the Dual Specificity Enzyme RluF*. Journal of Biological Chemistry, 291, 22327–22337.

78. Wrzesinski, J., Nurse, K., Bakin, A., Lane, B.G. and Ofengand, J. (1995) A dual-specificity pseudouridine synthase: an Escherichia coli synthase purified and cloned on the basis of its specificity for psi 746 in 23S RNA is also specific for psi 32 in tRNA(phe). RNA, 1, 437–448.

79. Ofengand, J. and Bakin, A. (1997) Mapping to nucleotide resolution of pseudouridine residues in large subunit ribosomal RNAs from representative eukaryotes, prokaryotes, archaebacteria, mitochondria and chloroplasts1. Journal of Molecular Biology, 266, 246–268.

80. Popova, A.M., Jain, N., Dong, X., Abdollah-Nia, F., Britton, R.A. and Williamson, J.R. (2024) Complete list of canonical post-transcriptional modifications in the Bacillus subtilis ribosome and their link to RbgA driven large subunit assembly. Nucleic Acids Res, 52, 11203–11217.

81. Hoang, C., Chen, J., Vizthum, C.A., Kandel, J.M., Hamilton, C.S., Mueller, E.G. and Ferré-D’Amaré, A.R. (2006) Crystal Structure of Pseudouridine Synthase RluA: Indirect Sequence Readout through Protein-Induced RNA Structure. Molecular Cell, 24, 535–545.

82. LaMarre, J.M., Howden, B.P. and Mankin, A.S. (2011) Inactivation of the Indigenous Methyltransferase RlmN in Staphylococcus aureus Increases Linezolid Resistance. Antimicrobial Agents and Chemotherapy, 55, 2989–2991.

83. Fitzsimmons, C.M. and Fujimori, D.G. (2016) Determinants of tRNA Recognition by the Radical SAM Enzyme RlmN. PLOS ONE, 11, e0167298.

84. Schwalm, E.L., Grove, T.L., Booker, S.J. and Boal, A.K. (2016) Crystallographic capture of a radical S-adenosylmethionine enzyme in the act of modifying tRNA. Science, 352, 309–312.

85. Lee, W.L., Sinha, A., Lam, L.N., Loo, H.L., Liang, J., Ho, P., Cui, L., Chan, C.S.C., Begley, T., Kline, K.A., et al. (2023) An RNA modification enzyme directly senses reactive oxygen species for translational regulation in Enterococcus faecalis. Nat Commun, 14, 4093.

86. Anton, B.P., Russell, S.P., Vertrees, J., Kasif, S., Raleigh, E.A., Limbach, P.A. and Roberts, R.J. (2010) Functional characterization of the YmcB and YqeV tRNA methylthiotransferases of Bacillus subtilis. Nucleic Acids Res, 38, 6195–6205.

87. Herbert, S., Ziebandt, A.-K., Ohlsen, K., Schäfer, T., Hecker, M., Albrecht, D., Novick, R. and Götz, F. (2010) Repair of Global Regulators in Staphylococcus aureus 8325 and Comparative Analysis with Other Clinical Isolates. Infection and Immunity, 78, 2877–2889.

88. Esakova, O.A., Grove, T.L., Yennawar, N.H., Arcinas, A.J., Wang, B., Krebs, C., Almo, S.C. and Booker, S.J. (2021) Structural basis for tRNA methylthiolation by the radical SAM enzyme MiaB. Nature, 597, 566–570.

89. Forouhar, F., Arragain, S., Atta, M., Gambarelli, S., Mouesca, J.-M., Hussain, M., Xiao, R., Kieffer-Jaquinod, S., Seetharaman, J., Acton, T.B., et al. (2013) Two Fe-S clusters catalyze sulfur insertion by radical-SAM methylthiotransferases. Nat Chem Biol, 9, 333–338.

90. Faivre, B., Lombard, M., Fakroun, S., Vo, C.-D.-T., Goyenvalle, C., Guérineau, V., Pecqueur, L., Fontecave, M., De Crécy-Lagard, V., Brégeon, D., et al. Dihydrouridine synthesis in tRNAs is under reductive evolution in Mollicutes. RNA Biol, 18, 2278–2289.

91. Byrne, R.T., Jenkins, H.T., Peters, D.T., Whelan, F., Stowell, J., Aziz, N., Kasatsky, P., Rodnina, M.V., Koonin, E.V., Konevega, A.L., et al. (2015) Major reorientation of tRNA substrates defines specificity of dihydrouridine synthases. Proceedings of the National Academy of Sciences, 112, 6033–6037.

92. Yu, F., Tanaka, Y., Yamashita, K., Suzuki, T., Nakamura, A., Hirano, N., Suzuki, T., Yao, M. and Tanaka, I. (2011) Molecular basis of dihydrouridine formation on tRNA. Proceedings of the National Academy of Sciences, 108, 19593–19598.

93. Savage, D.F., de Crécy-Lagard, V. and Bishop, A.C. (2006) Molecular determinants of dihydrouridine synthase activity. FEBS Letters, 580, 5198–5202.

94. Uhlenbeck, O.C. and Schrader, J.M. (2018) Evolutionary tuning impacts the design of bacterial tRNAs for the incorporation of unnatural amino acids by ribosomes. Current Opinion in Chemical Biology, 46, 138–145.

95. Green, C.J. and Vold, B.S. (1993) Staphylococcus aureus has clustered tRNA genes. Journal of Bacteriology, 175, 5091–5096.

96. Giannouli, S., Kyritsis, A., Malissovas, N., Becker, H.D. and Stathopoulos, C. (2009) On the role of an unusual tRNAGly isoacceptor in *Staphylococcus aureus*. Biochimie, 91, 344–351.

97. Alkatib, S., Scharff, L.B., Rogalski, M., Fleischmann, T.T., Matthes, A., Seeger, S., Schöttler, M.A., Ruf, S. and Bock, R. (2012) The Contributions of Wobbling and Superwobbling to the Reading of the Genetic Code. PLOS Genetics, 8, e1003076.

98. Kohl, M.P., Chane-Woon-Ming, B., Bahena-Ceron, R., Jaramillo-Ponce, J., Antoine, L., Herrgott, L., Romby, P. and Marzi, S. (2024) Ribosome Profiling Methods Adapted to the Study of RNA-Dependent Translation Regulation in Staphylococcus aureus. In Arluison, V., Valverde, C. (eds), Bacterial Regulatory RNA: Methods and Protocols. Springer US, New York, NY, pp. 73–100.

99. Mohammad, F., Green, R. and Buskirk, A.R. (2019) A systematically-revised ribosome profiling method for bacteria reveals pauses at single-codon resolution. eLife, 8, e42591.

100. Rogalski, M., Karcher, D. and Bock, R. (2008) Superwobbling facilitates translation with reduced tRNA sets. Nat Struct Mol Biol, 15, 192–198.

101. Teixeira, C., Vandenesch, F. and Moreau, K. (2025) tRNA modifications as regulators of bacterial virulence and stress responses. PLOS Pathogens, 21, e1013600.

102. Chionh, Y.H., McBee, M., Babu, I.R., Hia, F., Lin, W., Zhao, W., Cao, J., Dziergowska, A., Malkiewicz, A., Begley, T.J., et al. (2016) tRNA-mediated codon-biased translation in mycobacterial hypoxic persistence. Nat Commun, 7, 13302.

103. Kimura, S. and Waldor, M.K. (2019) The RNA degradosome promotes tRNA quality control through clearance of hypomodified tRNA. Proceedings of the National Academy of Sciences, 116, 1394–1403.

104. Alexandrov, A., Chernyakov, I., Gu, W., Hiley, S.L., Hughes, T.R., Grayhack, E.J. and Phizicky, E.M. (2006) Rapid tRNA Decay Can Result from Lack of Nonessential Modifications. Molecular Cell, 21, 87–96.

105. Shi, H., Wei, J. and He, C. (2019) Where, When, and How: Context-Dependent Functions of RNA Methylation Writers, Readers, and Erasers. Molecular Cell, 74, 640–650.

106. Björk, G.R. and Hagervall, T.G. (2014) Transfer RNA Modification: Presence, Synthesis, and Function. EcoSal Plus, 6, 10.1128/ecosalplus.ESP-0007–2013.

107. de Crécy-Lagard, V., Marck, C., Brochier-Armanet, C. and Grosjean, H. (2007) Comparative RNomics and Modomics in Mollicutes: Prediction of Gene Function and Evolutionary Implications. IUBMB Life, 59, 634–658.

108. Mandler, M.D., Maligireddy, S.S., Guiblet, W.M., Fitzsimmons, C.M., McDonald, K.S., Warrell, D.L. and Batista, P.J. (2024) The modification landscape of Pseudomonas aeruginosa tRNAs. RNA, 30, 1025–1040.

109. Wolf, P., Villette, C., Zumsteg, J., Heintz, D., Antoine, L., Chane-Woon-Ming, B., Droogmans, L., Grosjean, H. and Westhof, E. (2020) Comparative patterns of modified nucleotides in individual tRNA species from a mesophilic and two thermophilic archaea. RNA, 26, 1957–1975.

110. Yu, N., Jora, M., Solivio, B., Thakur, P., Acevedo-Rocha, C.G., Randau, L., de Crécy-Lagard, V., Addepalli, B. and Limbach, P.A. (2019) tRNA Modification Profiles and Codon-Decoding Strategies in Methanocaldococcus jannaschii. Journal of Bacteriology, 201, 10.1128/jb.00690-18.

111. Panosyan, H., Traube, F.R., Brandmayr, C., Wagner, M. and Carell, T. (2022) tRNA modification profiles in obligate and moderate thermophilic bacilli. Extremophiles, 26, 11.

112. Zhong, W., Pasunooti, K.K., Balamkundu, S., Wong, Y.H., Nah, Q., Gadi, V., Gnanakalai, S., Chionh, Y.H., McBee, M.E., Gopal, P., et al. (2019) Thienopyrimidinone Derivatives That Inhibit Bacterial tRNA (Guanine37-N1)-Methyltransferase (TrmD) by Restructuring the Active Site with a Tyrosine-Flipping Mechanism. J. Med. Chem., 62, 7788–7805.

113. Shapiro, A.B., Plant, H., Walsh, J., Sylvester, M., Hu, J., Gao, N., Livchak, S., Tentarelli, S. and Thresher, J. (2014) Discovery of ATP-Competitive Inhibitors of tRNAIle Lysidine Synthetase (TilS) by High-Throughput Screening. SLAS Discovery, 19, 1137–1146.

114. Kopina, B.J., Missoury, S., Collinet, B., Fulton, M.G., Cirio, C., van Tilbeurgh, H. and Lauhon, C.T. (2021) Structure of a reaction intermediate mimic in t6A biosynthesis bound in the active site of the TsaBD heterodimer from Escherichia coli. Nucleic Acids Res, 49, 2141–2160.

115. Bahena-Ceron, R., Jaramillo-Ponce, J., Kanazawa, H., Antoine, L., Wolf, P., Marchand, V., Klaholz, B.P., Motorin, Y., Romby, P. and Marzi, S. (2023) Methods to Analyze Post-transcriptional Modifications Applied to Stable RNAs in Staphylococcus aureus. In Barciszewski, J. (ed), RNA Structure and Function. Springer International Publishing, Cham, pp. 233–258.

116. Motorin, Y., Marchand, V., Motorin, Y. and Marchand, V. (2021) Analysis of RNA Modifications by Second- and Third-Generation Deep Sequencing: 2020 Update. Genes, 12.

117. Motorin, Y. and Helm, M. (2019) Methods for RNA Modification Mapping Using Deep Sequencing: Established and New Emerging Technologies. Genes (Basel), 10, 35.

118. Schaening-Burgos, C., LeBlanc, H., Fagre, C., Li, G.-W. and Gilbert, W.V. (2024) RluA is the major mRNA pseudouridine synthase in Escherichia coli. PLOS Genetics, 20, e1011100.

119. Pierrel, F., Douki, T., Fontecave, M. and Atta, M. (2004) MiaB Protein Is a Bifunctional Radical-*S*-Adenosylmethionine Enzyme Involved in Thiolation and Methylation of tRNA*. Journal of Biological Chemistry, 279, 47555–47563.

120. Moukadiri, I., Garzón, M.-J., Björk, G.R. and Armengod, M.-E. (2014) The output of the tRNA modification pathways controlled by the Escherichia coli MnmEG and MnmC enzymes depends on the growth conditions and the tRNA species. Nucleic Acids Res, 42, 2602–2623.

121. Yamada, Y. and Ishikura, H. (1981) The Presence of N-[(9-β-D-Ribofuranosyl-2-methylthiopurin-6-yl)carbamoyl]threonine in Lysine tRNA1 from Bacillus subtilis1. J Biochem, 89, 1589–1591.

122. Moukadiri, I., Villarroya, M., Benítez-Páez, A. and Armengod, M.-E. (2018) Bacillus subtilis exhibits MnmC-like tRNA modification activities. RNA Biology, 15, 1167– 1173.

123. Roovers, M., Kaminska, K.H., Tkaczuk, K.L., Gigot, D., Droogmans, L. and Bujnicki, J.M. (2008) The YqfN protein of Bacillus subtilis is the tRNA: m1A22 methyltransferase (TrmK). Nucleic Acids Res, 36, 3252–3262.

124. Rajakovich, L.J., Tomlinson, J. and Dos Santos, P.C. (2012) Functional Analysis of Bacillus subtilis Genes Involved in the Biosynthesis of 4-Thiouridine in tRNA. Journal of Bacteriology, 194, 4933–4940.

125. Edwards, A.M., Black, K.A. and Dos Santos, P.C. (2022) Sulfur Availability Impacts Accumulation of the 2-Thiouridine tRNA Modification in Bacillus subtilis. Journal of Bacteriology, 204, e00009–22.

126. Jaroch, M., Sun, G., Tsui, H.-C.T., Reed, C., Sun, J., Jörg, M., Winkler, M.E., Rice, K.C., Dziergowska, A., Stich, T.A., et al. (2024) Alternate routes to mnm5s2U synthesis in Gram-positive bacteria. Journal of Bacteriology, 206, e00452–23.

127. Cho, G., Lee, J. and Kim, J. (2023) Identification of a novel 5-aminomethyl-2-thiouridine methyltransferase in tRNA modification. Nucleic Acids Res, 51, 1971–1983.

128. Rider, L.W., Ottosen, M.B., Gattis, S.G. and Palfey, B.A. (2009) Mechanism of Dihydrouridine Synthase 2 from Yeast and the Importance of Modifications for Efficient tRNA Reduction. J Biol Chem, 284, 10324–10333.

129. Grossoehme, N., Kehl-Fie, T.E., Ma, Z., Adams, K.W., Cowart, D.M., Scott, R.A., Skaar, E.P. and Giedroc, D.P. (2011) Control of Copper Resistance and Inorganic Sulfur Metabolism by Paralogous Regulators in Staphylococcus aureus. J Biol Chem, 286, 13522–13531.

130. Luebke, J.L., Shen, J., Bruce, K.E., Kehl-Fie, T.E., Peng, H., Skaar, E.P. and Giedroc, D.P. (2014) The CsoR-like sulfurtransferase repressor (CstR) is a persulfide sensor in Staphylococcus aureus. Mol Microbiol, 94, 1343–1360.

131. Crick, F.H.C. (1966) Codon—anticodon pairing: The wobble hypothesis. Journal of Molecular Biology, 19, 548–555.

132. Weixlbaumer, A., Murphy, F.V., Dziergowska, A., Malkiewicz, A., Vendeix, F.A.P., Agris, P.F. and Ramakrishnan, V. (2007) Mechanism of expanding the decoding capacity of tRNAs by modification of uridines. Nat Struct Mol Biol, 14, 498–502.

133. Kothe, U. and Rodnina, M.V. (2007) Codon Reading by tRNAAla with Modified Uridine in the Wobble Position. Molecular Cell, 25, 167–174.

134. Quax, T.E.F., Claassens, N.J., Söll, D. and van der Oost, J. (2015) Codon Bias as a Means to Fine-Tune Gene Expression. Mol Cell, 59, 149–161.

135. Ran, W. and Higgs, P.G. (2010) The Influence of Anticodon–Codon Interactions and Modified Bases on Codon Usage Bias in Bacteria. Mol Biol Evol, 27, 2129–2140.

136. Rapid and Efficient Genome Editing in Staphylococcus aureus by Using an Engineered CRISPR/Cas9 System | Journal of the American Chemical Society.

137. CCTop: An Intuitive, Flexible and Reliable CRISPR/Cas9 Target Prediction Tool | PLOS One.

138. Wu, J. and McLuckey, S.A. (2004) Gas-phase fragmentation of oligonucleotide ions. International Journal of Mass Spectrometry, 237, 197–241.

139. Kellner, S., Ochel, A., Thüring, K., Spenkuch, F., Neumann, J., Sharma, S., Entian, K.-D., Schneider, D. and Helm, M. (2014) Absolute and relative quantification of RNA modifications via biosynthetic isotopomers. Nucleic Acids Res, 42, e142.

140. Zhang, L.-S., Ye, C., Ju, C.-W., Gao, B., Feng, X., Sun, H.-L., Wei, J., Yang, F., Dai, Q. and He, C. (2024) BID-seq for transcriptome-wide quantitative sequencing of mRNA pseudouridine at base resolution. Nat Protoc, 19, 517–538.

141. Jacobson, M. and Hedgcoth, C. (1970) Determination of 5, 6-dihydrouridine in ribonucleic acid. Analytical Biochemistry, 34, 459–469.

142. Hunninghake, D. and Grisolia, S. (1966) A sensitive and convenient micromethod for estimation of urea, citrulline, and carbamyl derivatives. Analytical Biochemistry, 16, 200–205.

143. Bouyssié, D., Hesse, A.-M., Mouton-Barbosa, E., Rompais, M., Macron, C., Carapito, C., Gonzalez de Peredo, A., Couté, Y., Dupierris, V., Burel, A., et al. (2020) Proline: an efficient and user-friendly software suite for large-scale proteomics. Bioinformatics, 36, 3148–3155.

144. van der Toorn, W., Naarmann-de Vries, I.S., Liu-Wei, W., Dieterich, C. and von Kleist, M. (2025) WarpDemuX-tRNA: barcode multiplexing for nanopore tRNA sequencing. Nucleic Acids Res, 53, gkaf873.

145. Mohammad, F. and Buskirk, A.R. (2019) Protocol for Ribosome Profiling in Bacteria. Bio Protoc, 9, e3468.

