## Supplementary Material for "Complete post-transcriptional modification profiles in individual *Staphylococcus aureus* tRNA species"

* To whom correspondence should be addressed.

### SUPPLEMENTARY METHODS

#### Cloning and production of RNase U2

The plasmid pET22b-MPB-U2 was constructed to express *Ustilago sphaerogena* RNase U2 in the *E. coli* periplasm, fused at its N-terminus to the maltose binding protein (MBP) and carrying a C-terminal 6xHis tag. The RNase U2 coding sequence was amplified from plasmid pET22b-U2 (1) using primers BamHITEVU2_F and U2TEVXhoI_R (Table S2), which introduced TEV protease cleavage sites at both termini and restriction sites for cloning into pET22b. The resulting plasmid, pET22b-U2(TEV), was linearized using NcoI and BamHI restriction sites, located between the *pelB* signal sequence and the RNase U2 coding region. The MBP coding sequence was amplified from plasmid pMBP-MS2 (2) using primers MBP_F and MBP_R, generating a fragment with ends complementary to those of the linearized pET22b-U2(TEV). The two fragments were assembled using the NEBuilder HiFi DNA assembly kit (New England Biolabs) to obtain the final construct. Recombinant MBP-U2 was produced in *E. coli* BL21(DE3) cells grown in 500 mL of LB medium supplemented with 100 µg/mL ampicillin. Cultures were incubated at 30°C until an $OD_{600}$ of 0.8 was reached, and protein expression was induced with 0.5 mM IPTG overnight at 16°C. Cells were harvested by centrifugation, washed with PBS, and stored at -20°C. Sequential purification on amylose and Ni-NTA resins was performed to ensure purification of the full-length MBP-U2 fusion protein. Bacteria were thawed on ice, resuspended in 25 mL of binding buffer (20 mM Tris-HCl pH 7.5, 250 mM NaCl), and lysed by sonication at 120 V for 7 min on ice (Annemasse Ultrasons apparatus). The lysate was clarified by centrifugation at 160000 × *g* for 1 h at 4°C, and the supernatant was loaded by gravity onto a 1-mL amylose resin column (New England Biolabs) equilibrated with binding buffer. The column was washed sequentially with 15 mL of binding buffer, 15 mL of buffer containing 500 mM NaCl, and again with 15 mL of binding buffer, followed by elution with 12 mL of buffer 20 mM Tris-HCl (pH 7.5), 250 mM NaCl, and 10 mM maltose. The eluate was then applied by gravity to a 1 mL Ni-NTA column (Qiagen) equilibrated with binding buffer, washed with 15 mL of the same buffer, and eluted with 12 mL of buffer 10 mM Tris-HCl (pH 7.5), 250 mM NaCl, 20% glycerol, and 500 mM imidazole. The sample was concentrated to 1 mL using an Amicon ultrafiltration devicce (MWCO 50 kDa, Merck Millipore) and dialyzed in 100 mL of storage buffer 100 mM ammonium acetate (pH 4.7) and 50% glycerol.

#### Cloning and purification of *S. aureus* DusB2

The *dusB2* (HG001_00036) coding sequence was amplified from *S. aureus* HG001 genomic DNA using primers DusB2exp_F and DusB2exp_R (Table S2) and cloned into a modified pET15b vector *via* NcoI and XhoI restriction sites. This vector allows production of a recombinant protein containing a C-terminal TEV protease cleavage site followed by a 6xHis tag. The target protein contained a glycine insertion after the first methionine, introduced by the NcoI site. Recombinant DusB2 was produced in *E. coli* BL21(DE3) pLyS at 30°C in LB medium supplemented with 100 µg/mL ampicillin and 34 µg/mL chloramphenicol. Cells were grown to $OD_{600}$ 0.6 - 0.8, induced with 0.5 mM IPTG and incubated for an additional 3-5 h before harvesting by centrifugation and washing with PBS. Protein purification was performed by fast protein liquid chromatography using an ÄKTA system (Cytiva). The frozen bacterial pellet was thawed on ice and resuspedend in buffer 50 mM sodium phosphate pH 8.0, 300 mM NaCl, 10 mM imidazole, 10% glycerol, and 5 mM β-mercaptoethanol. Cells were lysed by sonication at 120V for 7 min on ice (Annemasse Ultrasons apparatus) and the crude extract was centrifuged at 160000 × *g* for 1 h at 4°C. The supernatant was loaded onto a 1-mL Ni-NTA agarose column (Qiagen) equlibrated with the same buffer and washed with 40 mL prior to elution of bound proteins using a linear imidazole gradient from 10 to 500 mM over 20 mL. Fractions containing recombinant DusB2 (~40 kDa) according to SDS-PAGE were pooled, supplemented with 6xHis-tagged TEV protease (1 µg of TEV protease per 25 µg protein, Protean), and dialyzed overnight at 4°C with gentle agitation in buffer lacking imidazole. A second Ni-NTA chromatography was then performed under the same conditions, except that the column was washed with 20 mL buffer, to recover untagged DusB2 in the flow-through while retaining TEV protease, uncleaved DusB2, and non-specific contaminants on the column. Untagged DusB2 was concentrated using an Amicon ultrafiltration device (MWCO 10 kDa, Merck Millipore) and dialyzed as described above in buffer 50 mM sodium phosphate pH 8.0, 50 mM NaCl, 10% glycerol and 5 mM β-mercaptoethanol. The sample was loaded onto a monoQ 5/50 anion-exchange column (Cytiva) equilibrated in the same buffer. The column was washed with 10 mL of buffer and eluted with linear NaCl gradient from 50 to 1000 mM over 20 mL. Fractions corresponding to pure DusB2 were pooled, concentrated, and dialyzed in storage buffer 25 mM HEPES-NaOH pH 7.5, 150 mM NaCl, 50% glycerol, and 5 mM β-mercaptoethanol.

**Table S1.** List of primers and DNA oligonucleotides used in the present study. Restriction sites are underlined.

| Name | Sequence | Purpose |
|---|---|---|
| Dus_5UP_F | gtaaagtctagaattaatatcatttctttagaacctgg | Deletion of *dusB2* gene using pCasSA CRISPR-Cas9 |
| Dus_5UP_R | ccataaactttattactcataccctctttataattagtatctcg | |
| Dus_3DN_F | cgagatactaattataaagagggtatgagtaataaagtttatgg | |
| Dus_3DN_R | ctttacctcgagataaccttgttgaaatcgacgg | |
| gRNA_DusB2_F | gaaatttagtcgcttgaataattt | |
| gRNA_DusB2_R | aaacaaattattcaagcgactaaa | |
| Dus_up | cgatgggcttagttctctctgcacc | |
| Dus_mid | gacttccctttcaaatggttgcatc | |
| Dus_down | cgtcaaataaagcatcttttggcgttgc | |
| SphI_P_DusB2_F | taagcagcatgcgcgaatagcagaaaaataggtatcgc | Complementation plasmid pCN38-DusB2 |
| KpnI_TT_DusB2_R | tgcttaggtacccgggagttatcctatcaggaaaaccaag | |
| DusB2exp_F | cttaagccatgggtaaagaaaatttttggagtgaattacc | Production of *S. aureus* DusB2 in *E. coli* |
| DusB2exp_R | cgccgcctcgagtaattcaattttaacgtcttcgtcc | |
| BamHITEVU2_F | agcaggatccggagaatctttattttcagtcttgtgatatcccacagtctacgaactgcg | Production of RNase U2 |
| U2TEVXhoI_R | ccgcctcgagagactgaaaataaagattctcactacactgggtgaatccatcgtacgacg | |
| MBP_F | gctgcccagccggcgatggccatggataaaactgaagaaggtaaactgg | |
| MBP_R | ctgaaaataaagattctccggatccgaagtctgcgcgtctttc | |
| Lys_UUU_Biotin | 5Biotin/ctctgattaaaagtcagatgctctaccaactgagcta | Purification of individual tRNAs |
| Thr_UGU_Biotin | acaagtcagttgctctaccaattgag/3Biotin | |
| Gly_P2_Biotin | atcagcttggaaggctgaggttttgccattaaacta/3Biotin | |
| Gly_NP1NP2_Biotin | 5Biotin/tcgttccgggaaggaacgtgttctaaaagttgaacta | |
| Gly_NP3_Biotin | tcattccaggaaggaatgtattctaagagttgaaata/3Biotin | |

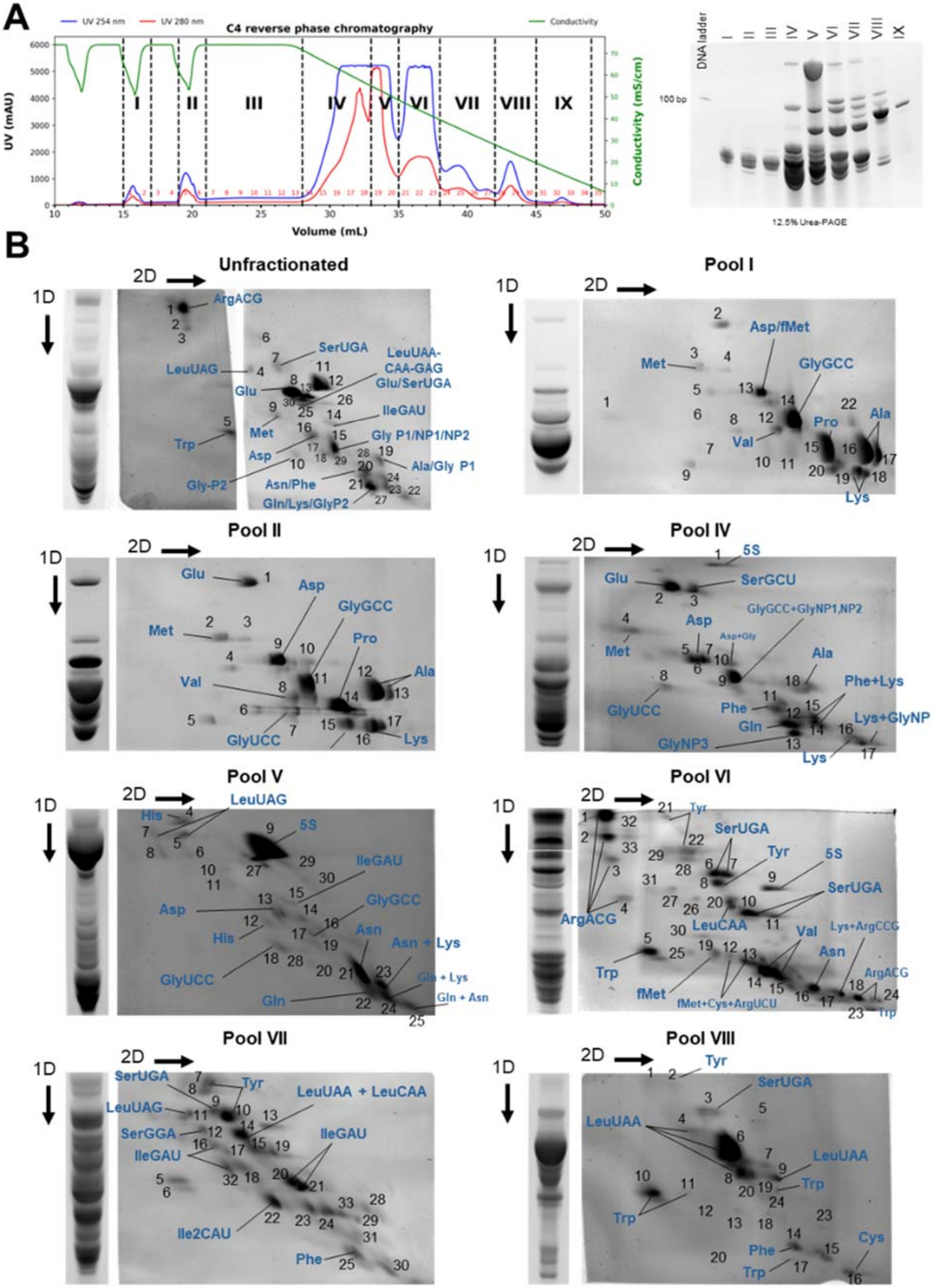
A
UV 254 nm
UV 280 nm
C4 reverse phase chromatography
Conductivity
UV (mAU)
Conductivity (mS/cm)
Volume (mL)
I
II
III
IV
V
VI
VII
VIII
IX
DNA ladder
100 bp
12.5% Urea-PAGE
B
Unfractionated
Pool I
Pool II
Pool IV
Pool V
Pool VI
Pool VII
Pool VIII
1D
2D
ArgACG
SerUGA
LeuUAG
LeuUAA-CAA-GAG
Glu/SerUGA
Glu
IleGAU
Met
Trp
Asp
Gly P1/NP1/NP2
Ala/Gly P1
Gly-P2
Asn/Phe
Gln/Lys/GlyP2
Asp/fMet
GlyGCC
Val
Pro
Ala
Lys
Glu
Asp
GlyUCC
5S
SerGCU
GlyGCC+GlyNP1,NP2
Asp+Gly
Phe+Lys
Phe
Gln
Lys+GlyNP
GlyNP3
His
Asn
Asn + Lys
Gln + Lys
Gln + Asn
Tyr
LeuCAA
fMet
fMet+Cys+ArgUCU
Lys+ArgCCG
LeuUAA + LeuCAA
SerGGA
Ile2CAU
LeuUAA
Cys

**Figure S1. Isolation of individual tRNA species for UPLC-MS/MS analysis.** (**A**) Chromatographic separation of bulk tRNA on a reversed-phase C4 column. The chromatogram (left) shows UV absorbance signals at 254 (blue) and 280 nm (red), conductivity (green) and the collected fractions (numbers). Sample loading was performed in three sequential injections, visible as drops in conductivity before eluting with a linear methanol gradient. Fractions were combined in pools as indicated by Roman numerals between dashed lines: I (1-2), II (5-6), III (7-13), IV (14-18), V (19-20), VI (21-23), VII (24-27), VIII (28-30), IX (31-34). The resulting tRNA pools were analyzed by denaturing polyacrylamide gel electrophoresis (right). Smaller tRNAs eluted earlier than longer species. Pools I and II were not retained and eluted after 1 column volume. Pool IX consisted mainly of tRNA Leu(UAA). (**B**) 2D-PAGE of tRNA pools. Representative gels are shown for bulk tRNA (unfractionated) and for pools I, II, IV, V, VI, VII and VIII resulting from reversed-phase chromatography. Pool III is not shown, as its profile closely resembled pool II. In the first dimension, tRNA were separated under denaturing conditions, and the excised 1D lane was embedded in a semi-denaturing gel for the second dimension, allowing resolution of individual tRNAs.

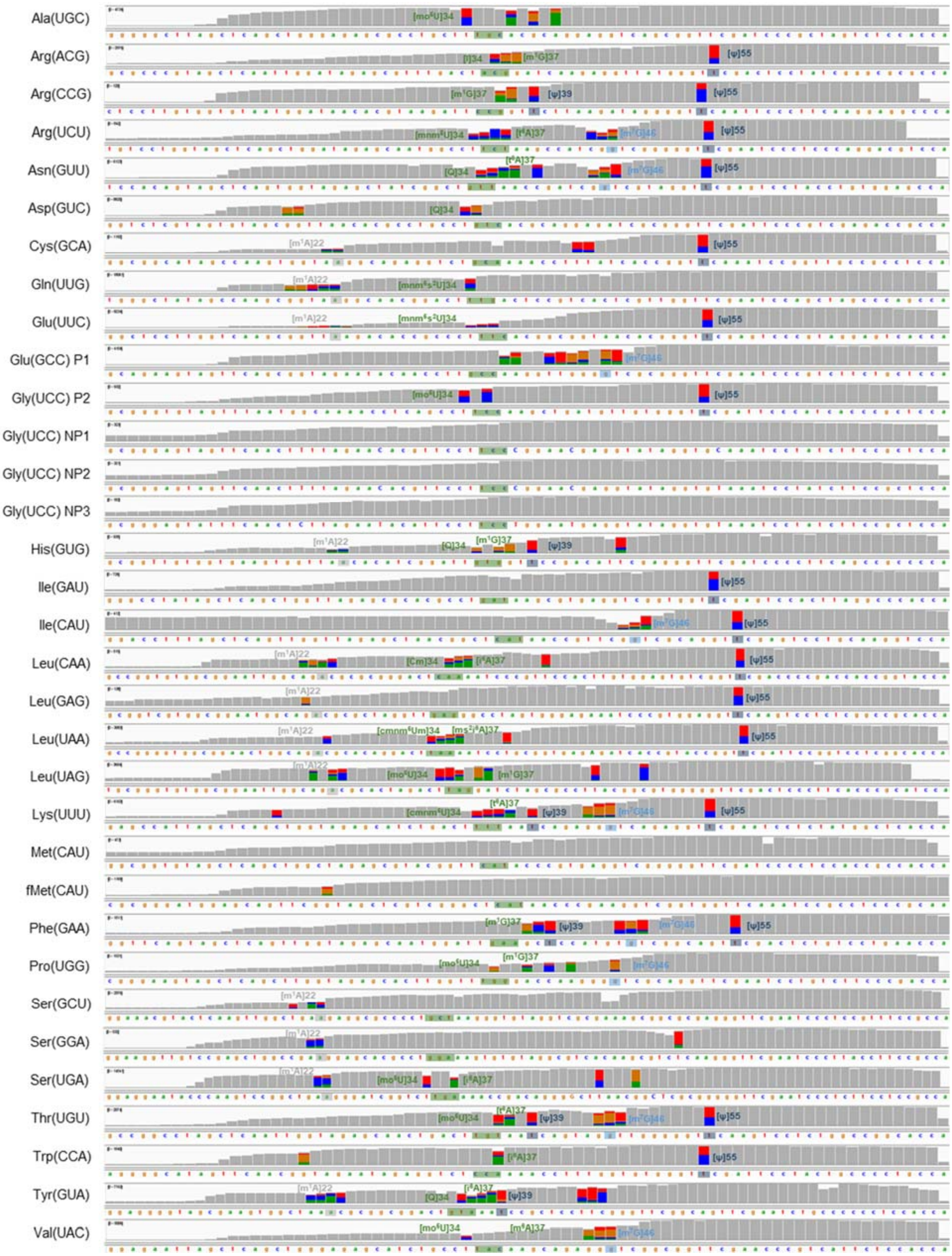
Ala(UGC)
Arg(ACG)
Arg(CCG)
Arg(UCU)
Asn(GUU)
Asp(GUC)
Cys(GCA)
Gln(UUG)
Glu(UUC)
Glu(GCC) P1
Gly(UCC) P2
Gly(UCC) NP1
Gly(UCC) NP2
Gly(UCC) NP3
His(GUG)
Ile(GAU)
Ile(CAU)
Leu(CAA)
Leu(GAG)
Leu(UAA)
Leu(UAG)
Lys(UUU)
Met(CAU)
fMet(CAU)
Phe(GAA)
Pro(UGG)
Ser(GCU)
Ser(GGA)
Ser(UGA)
Thr(UGU)
Trp(CCA)
Tyr(GUA)
Val(UAC)

**Figure S2. Representative Nanopore sequencing profiles of *S. aureus* tRNAs.** IGV snapshots show the sequence coverage for all 30 proteogenic species, as well as the 3 non-proteogenic tRNA Gly. Positions with mismatch frequencies above the default IGV threshold are colored according to the mismatched base: A (green), C (blue), G (orange) and T (red). The same color code is applied to nucleotides in the reference sequence. The anticodon of each tRNA is highlighted with a green rectangle, and modifications at position 34 and/or 37 are indicated when a mismatch signal is detected. A22 is highlighted with a gray rectangle in tRNAs showing a signal due to $m^1A$. G46 is highlighted in light blue when displaying a signal due to $m^7G$. Positions T39 and T55 are highlighted with dark blue rectangles when showing T-to-C mismatches, characteristic of ψ.

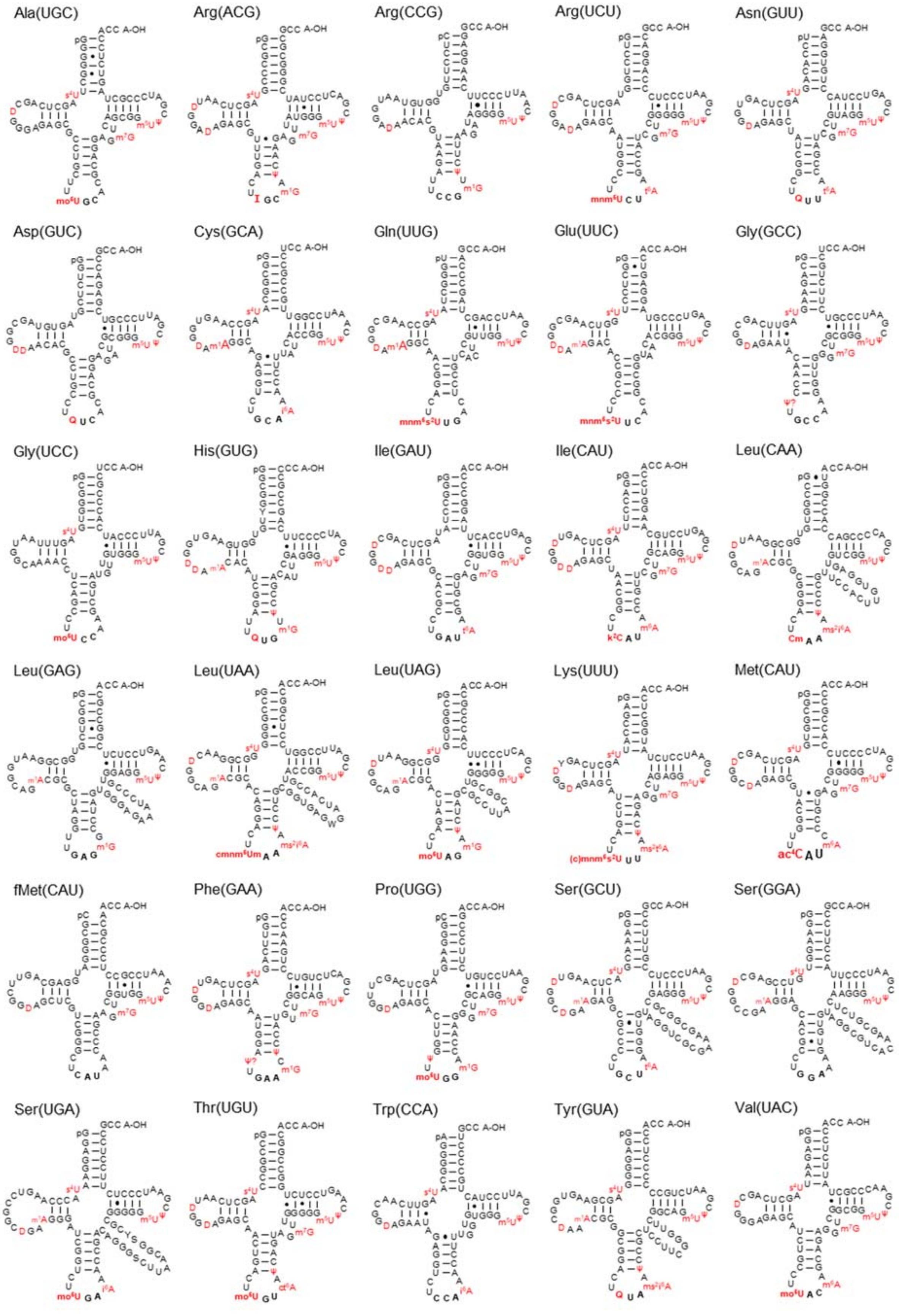
Ala(UGC)
Arg(ACG)
Arg(CCG)
Arg(UCU)
Asn(GUU)
Asp(GUC)
Cys(GCA)
Gln(UUG)
Glu(UUC)
Gly(GCC)
Gly(UCC)
His(GUG)
Ile(GAU)
Ile(CAU)
Leu(CAA)
Leu(GAG)
Leu(UAA)
Leu(UAG)
Lys(UUU)
Met(CAU)
fMet(CAU)
Phe(GAA)
Pro(UGG)
Ser(GCU)
Ser(GGA)
Ser(UGA)
Thr(UGU)
Trp(CCA)
Tyr(GUA)
Val(UAC)

**Figure S3. Atlas of tRNA modifications in *S. aureus*.** Cloverleaf representations of the 30 proteogenic tRNAs, with identified modified nucleosides highlighted in red. Anticodon sequences are shown in bold. Canonical Watson-Crick base pairs are indicated by lines, and G-U pairs by black dots. Single-nucleotide polymorphisms in tRNAs His(GUG), Leu(UAA), Lys(UUU), and Ser(UGA) are indicated using conventional nucleotide codes, Y (C or U), S: (G or C), and W (A or U). Chemical groups shown in parentheses denote alternative forms detected, meaning that fragments were observed with one modification, the other, or both.

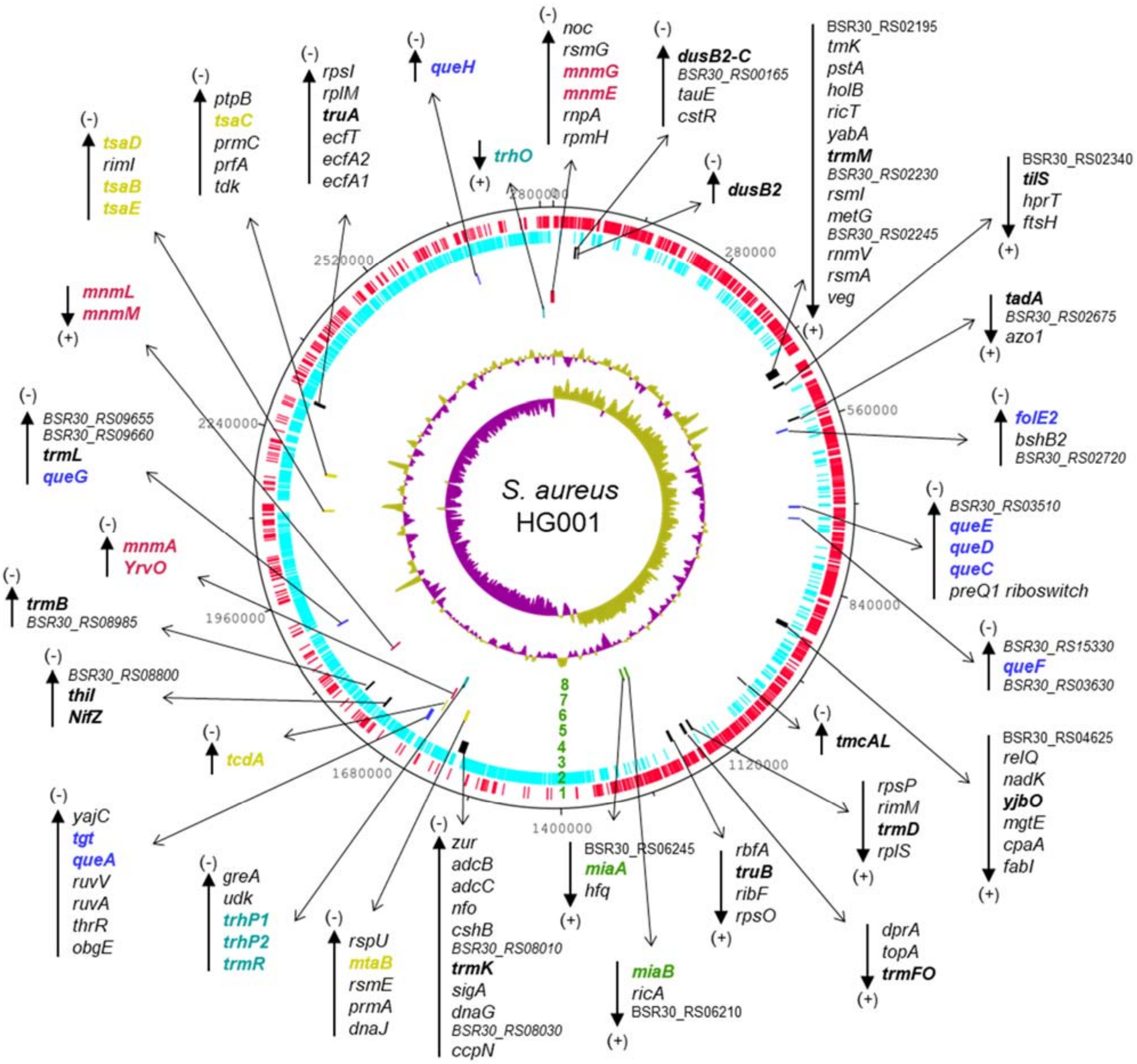
S. aureus
HG001
2800000
280000
560000
840000
1120000
1400000
1680000
1960000
2240000
2520000
8
7
6
5
4
3
2
1
(-)
noc
rsmG
mnmG
mnmE
rnpA
rpmH
(-)
dusB2-C
BSR30_RS00165
tauE
cstR
(-)
dusB2
BSR30_RS02195
tmK
pstA
holB
ricT
yabA
trmM
BSR30_RS02230
rsmI
metG
BSR30_RS02245
rnmV
rsmA
veg
(+)
BSR30_RS02340
tilS
hprT
ftsH
(+)
tadA
BSR30_RS02675
azo1
(+)
(-)
folE2
bshB2
BSR30_RS02720
(-)
BSR30_RS03510
queE
queD
queC
preQ1 riboswitch
(-)
BSR30_RS15330
queF
BSR30_RS03630
(-)
tmcAL
BSR30_RS04625
relQ
nadK
yjbO
mgtE
cpaA
fabI
(+)
rpsP
rimM
trmD
rplS
(+)
dprA
topA
trmFO
(+)
rbfA
truB
ribF
rpsO
(+)
miaB
ricA
BSR30_RS06210
(+)
BSR30_RS06245
miaA
hfq
(+)
(-)
zur
adcB
adcC
nfo
cshB
BSR30_RS08010
trmK
sigA
dnaG
BSR30_RS08030
ccpN
(-)
rspU
mtaB
rsmE
prmA
dnaJ
(-)
greA
udk
trhP1
trhP2
trmR
(-)
yajC
tgt
queA
ruvV
ruvA
thrR
obgE
(-)
tcdA
(-)
BSR30_RS08800
thiI
NifZ
(-)
trmB
BSR30_RS08985
(-)
mnmA
YrvO
(-)
BSR30_RS09655
BSR30_RS09660
trmL
queG
mnmL
mnmM
(+)
(-)
tsaD
rimI
tsaB
tsaE
(-)
ptpB
tsaC
prmC
prfA
tdk
(-)
rpsI
rplM
truA
ecfT
ecfA2
ecfA1
(-)
queH
trhO
(+)

**Figure S4. Genomic context of tRNA-modifying enzymes in *Staphylococcus aureus* HG001.** The circular genome plot displays the position of 43 tRNA modification enzymes (in bold) distributed in eight tracks. Tracks 1 (red) and 2 (light blue) represent all genes on the (+) and (-) strands, respectively. Track 3 (black) highlights stand-alone modifying enzymes, together with NifZ and ThiI, which participate in [$s^4$U]8 synthesis. Track 4 (blue) shows the genes of the [Q]34 synthesis pathway. Track 5 (light green) corresponds to the [$ms^2t^6$A]37 synthesis pathway. Track 6 (dark red) marks the genes required for the synthesis of [$xm^5$U]34 derivatives. Track 7 (dark teal) represents the [$mo^5$U]34 synthesis pathway. Track 8 (dark green) indicates MiaA and MiaB, which catalyze [$ms^2i^6$A]37 formation. Genes potentially co-transcribed with tRNA modification enzymes are displayed around the circle, with arrows denoting their orientation on the (+) or (-) strand. The outer inner track shows the mean-centered GC%, with values above the mean in green and below the mean in violet. The innermost track displays the GC skew, where transition from positive (green) to negative (violet) values, or vice versa, indicate the replication origin and terminus.

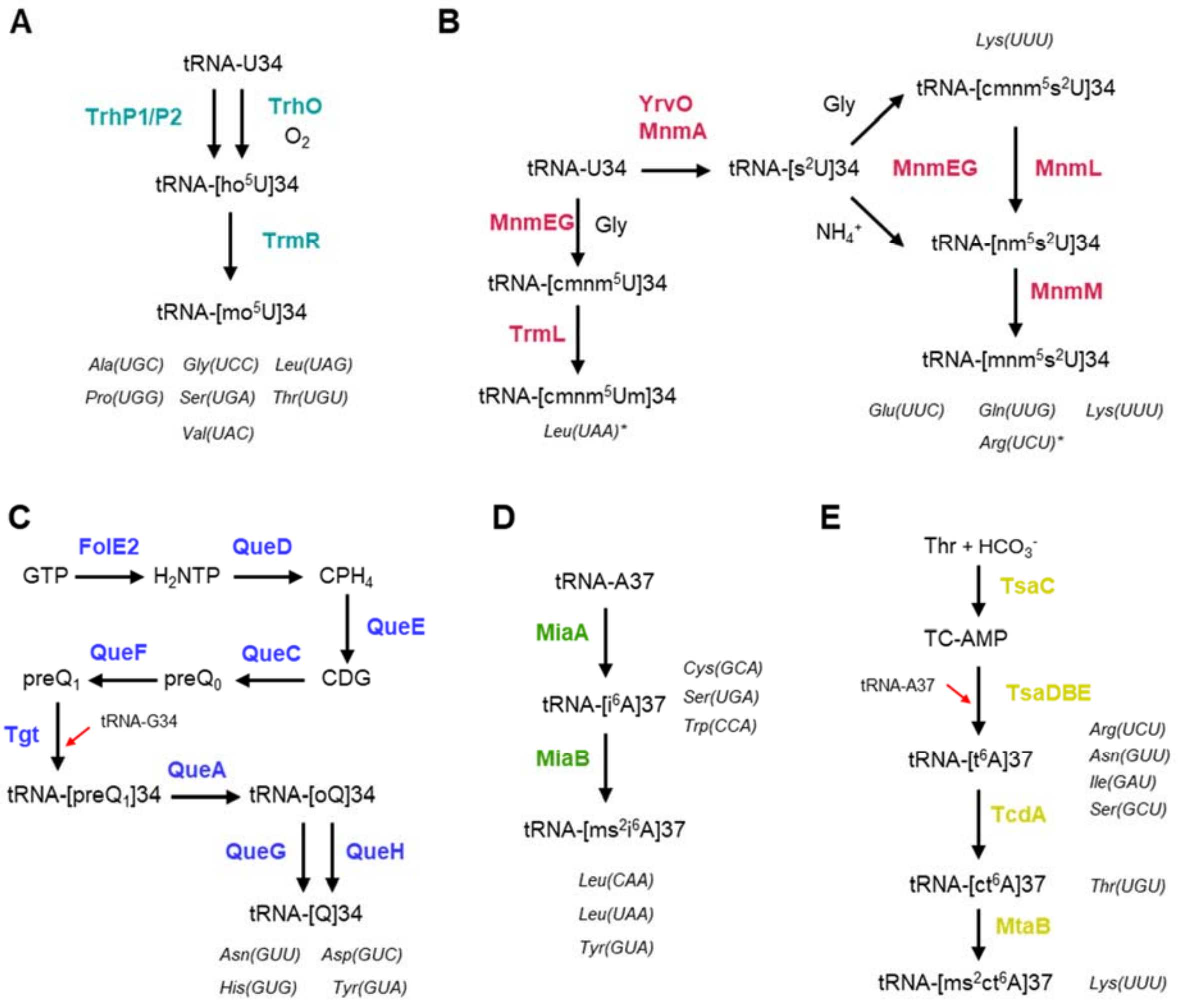
A
tRNA-U34
TrhP1/P2
TrhO
O2
tRNA-[ho5U]34
TrmR
tRNA-[mo5U]34
Ala(UGC) Gly(UCC) Leu(UAG)
Pro(UGG) Ser(UGA) Thr(UGU)
Val(UAC)
B
tRNA-U34
YrvO
MnmA
tRNA-[s2U]34
Gly
NH4+
MnmEG
Gly
tRNA-[cmnm5U]34
TrmL
tRNA-[cmnm5Um]34
Leu(UAA)*
Lys(UUU)
tRNA-[cmnm5s2U]34
MnmEG
MnmL
tRNA-[nm5s2U]34
MnmM
tRNA-[mnm5s2U]34
Glu(UUC) Gln(UUG) Lys(UUU)
Arg(UCU)*
C
GTP
FolE2
H2NTP
QueD
CPH4
QueE
CDG
QueC
preQ0
QueF
preQ1
Tgt
tRNA-G34
tRNA-[preQ1]34
QueA
tRNA-[oQ]34
QueG
QueH
tRNA-[Q]34
Asn(GUU) Asp(GUC)
His(GUG) Tyr(GUA)
D
tRNA-A37
MiaA
tRNA-[i6A]37
Cys(GCA)
Ser(UGA)
Trp(CCA)
MiaB
tRNA-[ms2i6A]37
Leu(CAA)
Leu(UAA)
Tyr(GUA)
E
Thr + HCO3-
TsaC
TC-AMP
tRNA-A37
TsaDBE
tRNA-[t6A]37
Arg(UCU)
Asn(GUU)
Ile(GAU)
Ser(GCU)
TcdA
tRNA-[ct6A]37
Thr(UGU)
MtaB
tRNA-[ms2ct6A]37
Lys(UUU)

**Figure S5. Pathways for the synthesis of complex modifications in *S. aureus* tRNAs.** The different tRNA species carrying each modification are shown in italics. (**A**) Synthesis of [$mo^5$U]34 proceeds through a two-step pathway: U34 is first hydroxylated by either the TrhP1/P2 complex or the oxygen-dependent TrhO, and the resulting [$ho^5$U]34 is methylated by TrmR. (**B**) Alternative pathways leading to [$xm^5$U]34 modifications. Thiolation at the C2 position of U34 is catalyzed by MnmA together with YrvO and occurs only in specific tRNAs. Those species marked with an asterisk were not thiolated in *S. aureus* HG001. The MnmEG complex uses either glycine to generate [$cmnm^5(s^2)$U]34 or ammonium to produce [$nm^5(s^2)$U]34. Among these, only [$nm^5(s^2)$U]34 can be further modified by MnmM to yield [$mnm^5(s^2)$U]34, which is the final modification in tRNAs Glu(UUC), Gln(UUG), and Arg(UCU). In contrast, [$cmnm^5(s^2)$U]34 remains the predominant species in tRNAs Lys(UUU) and Leu(UAA), but in the other tRNAs it is converted by MnmL into [$nm^5(s^2)$U]34, enabling completion of the pathway. In tRNA Leu(UAA), [$cmnm^5$U]34 undergoes further modification by TrmL, which methylates the 2'-O position of the ribose, yielding [$cmnm^5$Um]34. (**C**) Synthesis of [Q]34 in tRNAs Asn(GUU), Asp(GUC), His(GUG), and Tyr(GUA). Eight different enzymes are involved in *de novo* Q synthesis in *S. aureus*. FolE2, QueD, QueE, QueC, and QueF act sequentially to convert GTP into precursor $preQ_1$, which is then inserted at position 34 of the tRNA by the Tgt enzyme. QueA subsequently converts [$preQ_1$]34 into the epoxy form [oQ]34, which is then reduced by either QueG or QueH to yield the final [Q]34 modification. $H_2NTP$: 7,8-dihydroneopterin triphosphate, $CPH_4$: 6-carboxy-5,6,7,8-tetrahydropterin, CDG: 7-carboxy-7-deazaguanine. (**D**) Synthesis of [$ms^2i^6$A]37 proceeds through two consecutive enzymatic steps: MiaA installs an isopentenyl group at $N^6$ of A37, followed by MiaB-mediated methylthiolation at C2. Not all [$i^6$A]37-containing species are targeted by MiaB in *S. aureus.* (**E**) Biosynthesis of [$t^6$A]37 derivatives. TsaC uses *L*-threonine, bicarbonate and ATP to generate threonylcarbamoyl-AMP (TC-AMP). The TsaDBE complex transfers the TC group onto the $N^6$ atom of tRNA A37 to produce [$t^6$A]37. TcdA converts [$t^6$A]37 into the cyclic hydantoin [$ct^6$A]37, which was detected in Thr(UGU) but is potentially present in all six *S. aureus* tRNAs carrying [$t^6$A]37. MtaB in tRNA Lys(UUU) methylthiolate C2 position of the [$ct^6$A]37 base, yielding [$ms^2ct^6$A]37.

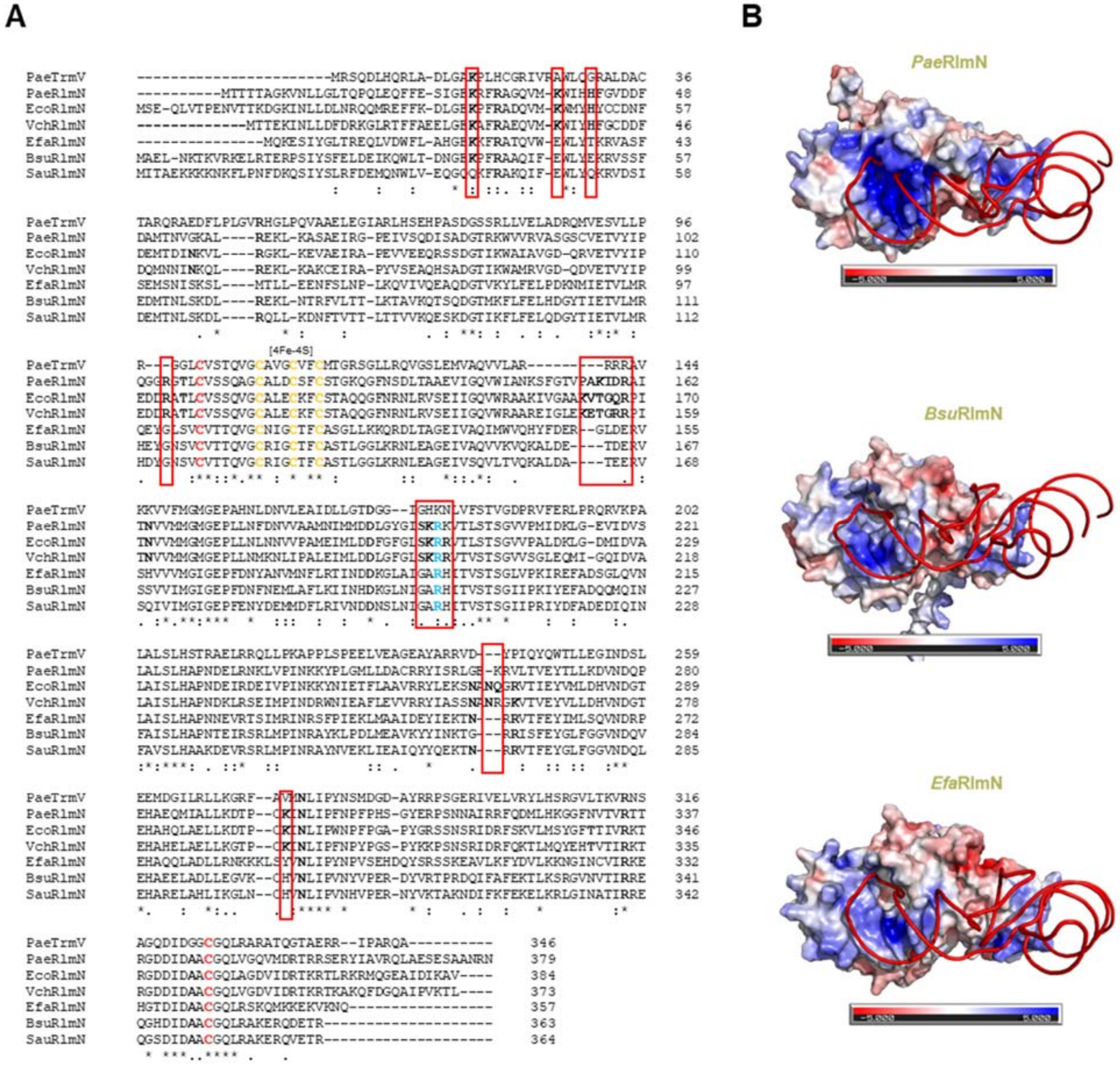
A
PaeTrmV ----------------------MRSQDLHQRLA-DLGAKPLHCGRIVRAWLQGRALDAC 36
PaeRlmN ----------MTTTTAGKVNLLGLTQPQLEQFFE-SIGEKRFRAGQVM-KWIHHFGVDDF 48
EcoRlmN MSE-QLVTPENVTTKDGKINLLDLNRQQMREFFK-DLGEKPFRADQVM-KWMYHYCCDNF 57
VchRlmN -------------MTTEKINLLDFDRKGLRTFFAEELGEKAFRAEQVM-KWIYHFGCDDF 46
EfaRlmN ---------------MQKESIYGLTREQLVDWFL-AHGEKKFRATQVW-EWLYTKRVASF 43
BsuRlmN MAEL-NKTKVRKELRTERPSIYSFELDEIKQWLT-DNGEKPFRAAQIF-EWLYEKRVSSF 57
SauRlmN MITAEKKKKNKFLPNFDKQSIYSLRFDEMQNWLV-EQGQQKFRAKQIF-EWLYQKRVDSI 58
PaeTrmV TARQRAEDFLPLGVRHGLPQVAAELEGIARLHSEHPASDGSSRLLVELADRQMVESVLLP 96
PaeRlmN DAMTNVGKAL----REKL-KASAEIRG-PEIVSQDISADGTRKWVVRVASGSCVETVYIP 102
EcoRlmN DEMTDINKVL----RGKL-KEVAEIRA-PEVVEEQRSSDGTIKWAIAVGD-QRVETVYIP 110
VchRlmN DQMNNINKQL----REKL-KAKCEIRA-PYVSEAQHSADGTIKWAMRVGD-QDVETVYIP 99
EfaRlmN SEMSNISKSL----MTLL-EENFSLNP-LKQVIVQEAQDGTVKYLFELPDKNMIETVLMR 97
BsuRlmN EDMTNLSKDL----REKL-NTRFVLTT-LKTAVKQTSQDGTMKFLFELHDGYTIETVLMR 111
SauRlmN DEMTNLSKDL----RQLL-KDNFTVTT-LTTVVKQESKDGTIKFLFELQDGYTIETVLMR 112
[4Fe-4S]
PaeTrmV R---GGLCVSTQVGCAVGCVFCMTGRSGLLRQVGSLEMVAQVVLAR----------RRRAV 144
PaeRlmN QGGRGTLCVSSQAGCALDCSFCSTGKQGFNSDLTAAEVIGQVWIANKSFGTVPAKIDRAI 162
EcoRlmN EDDRATLCVSSQVGCALECKFCSTAQQGFNRNLRVSEIIGQVWRAAKIVGAAKVTGQRPI 170
VchRlmN EDDRATLCVSSQVGCALECKFCSTAQQGFNRNLRVSEIIGQVWRAAREIGLEKETGRRPI 159
EfaRlmN QEYGLSVCVTTQVGCNIGCTFCASGLLKKQRDLTAGEIVAQIMWVQHYFDER--GLDERV 155
BsuRlmN HEYGNSVCVTTQVGCRIGCTFCASTLGGLKRNLEAGEIVAQVVKVQKALDE----TDERV 167
SauRlmN HDYGNSVCVTTQVGCRIGCTFCASTLGGLKRNLEAGEIVSQVLTVQKALDA----TEERV 168
PaeTrmV KKVVFMGMGEPAHNLDNVLEAIDLLGTDGG--IGHKNLVFSTVGDPRVFERLPRQRVKPA 202
PaeRlmN TNVVMMGMGEPLLNFDNVVAAMNIMMDDLGYGISKRKVTLSTSGVVPMIDKLG-EVIDVS 221
EcoRlmN TNVVMMGMGEPLLNLNNVVPAMEIMLDDFGFGLSKRRVTLSTSGVVPALDKLG-DMIDVA 229
VchRlmN TNVVMMGMGEPLLNMKNLIPALEIMLDDLGFGLSKRRVTVSTSGVVSGLEQMI-GQIDVA 218
EfaRlmN SHVVVMGIGEPFDNYANVMNFLRTINDDKGLAIGARHITVSTSGLVPKIREFADSGLQVN 215
BsuRlmN SSVVIMGIGEPFDNFNEMLAFLKIINHDKGLNIGARHITVSTSGIIPKIYEFADQQMQIN 227
SauRlmN SQIVIMGIGEPFENYDEMMDFLRIVNDDNSLNIGARHITVSTSGIIPRIYDFADEDIQIN 228
PaeTrmV LALSLHSTRAELRRQLLPKAPPLSPEELVEAGEAYARRVD---YPIQYQWTLLEGINDSL 259
PaeRlmN LALSLHAPNDELRNKLVPINKKYPLGMLLDACRRYISRLGE-KRVLTVEYTLLKDVNDQP 280
EcoRlmN LAISLHAPNDEIRDEIVPINKKYNIETFLAAVRRYLEKSNANQGRVTIEYVMLDHVNDGT 289
VchRlmN LAISLHAPNDKLRSEIMPINDRWNIEAFLEVVRRYIASSNANRGKVTVEYVLLDHVNDGT 278
EfaRlmN LAISLHAPNNEVRTSIMRINRSFPIEKLMAAIDEYIEKTN---RRVTFEYIMLSQVNDRP 272
BsuRlmN FAISLHAPNTEIRSRLMPINRAYKLPDLMEAVKYYINKTG---RRISFEYGLFGGVNDQV 284
SauRlmN FAVSLHAAKDEVRSRLMPINRAYNVEKLIEAIQYYQEKTN---RRVTFEYGLFGGVNDQL 285
PaeTrmV EEMDGILRLLKGRF--AVMNLIPYNSMDGD-AYRRPSGERIVELVRYLHSRGVLTKVRNS 316
PaeRlmN EHAEQMIALLKDTP--CKINLIPFNPFPHS-GYERPSNNAIRRFQDMLHKGGFNVTVRTT 337
EcoRlmN EHAHQLAELLKDTP--CKINLIPWNPFPGA-PYGRSSNSRIDRFSKVLMSYGFTTIVRKT 346
VchRlmN EHAHELAELLKGTP--CKINLIPFNPYPGS-PYKKPSNSRIDRFQKTLMQYEHTVTIRKT 335
EfaRlmN EHAQQLADLLRNKKKLSYVNLIPYNPVSEHDQYSRSSKEAVLKFYDVLKKNGINCVIRKE 332
BsuRlmN EHAEELADLLEGVK--CHVNLIPVNYVPER-DYVRTPRDQIFAFEKTLKSRGVNVTIRRE 341
SauRlmN EHARELAHLIKGLN--CHVNLIPVNHVPER-NYVKTAKNDIFKFEKELKRLGINATIRRE 342
PaeTrmV AGQDIDGGCGQLRARATQGTAERR--IPARQA---------- 346
PaeRlmN RGDDIDAACGQLVGQVMDRTRRSERYIAVRQLAESESAANRN 379
EcoRlmN RGDDIDAACGQLAGDVIDRTKRTLRKRMQGEAIDIKAV---- 384
VchRlmN RGDDIDAACGQLVGDVIDRTKRTKAKQFDGQAIPVKTL---- 373
EfaRlmN HGTDIDAACGQLRSKQMKKEKVKNQ----------------- 357
BsuRlmN QGHDIDAACGQLRAKERQDETR-------------------- 363
SauRlmN QGSDIDAACGQLRAKERQVETR-------------------- 364
B
PaeRlmN
-5.000
5.000
BsuRlmN
-5.000
5.000
EfaRlmN
-5.000
5.000

**Figure S6. Comparison of RNA-modifying enzymes responsible for $m^2A$ synthesis in bacteria.** (**A**) Sequence alignment (Clustal omega) of various bacterial RlmN orthologs and *P. aeruginosa* TrmV. Cysteine residues involved in coordination of [4Fe-4S] cluster (orange) and in catalysis (red) are strictly conserved. Residues mediating non-sequence-specific tRNA contacts in the crystal structure of the RlmN-tRNA Glu(UUC) complex (PDB 5HR6) are shown in bold in *E. coli* and in other orthologs when conserved. RNA-binding regions showing differences among the compared proteins are indicated with red boxes. RlmN from *S. aureus*, *B. subtilis*, and *E. faecalis*, all Gram-positive bacteria, lacks several residues involved in tRNA interactions. Arginine 206, which makes a sequence-specific contact with guanosine 29 critical for tRNA modification in *E. coli,* is conserved in all RlmN orthologs but not in *P. aeruginosa* TrmV. The absence of this residue, together with most of the non-sequence specific contacts in RlmN, suggests that TrmV uses a different mechanism for tRNA recognition. (**B**) AlphaFold3 models of RlmN from *P. aeruginosa*, *B. subtilis*, and *E. faecalis* showing their surface potential. Consistent with the absence of several residues involved in tRNA binding, both *B. subtilis* and *E. faecalis* RlmN display RNA-binding surfaces with weaker positive potential compared to *P. aeruginosa* RlmN. *Pae*: *Pseudomonas aeruginosa*, *Eco*: *Escherichia coli*, *Vch*: *Vibrio cholerae, Efa*: *Enterococcus faecalis*, *Bsu*: Bacillus *subtilis*, *Sau*: *Staphylococcus aureus.*

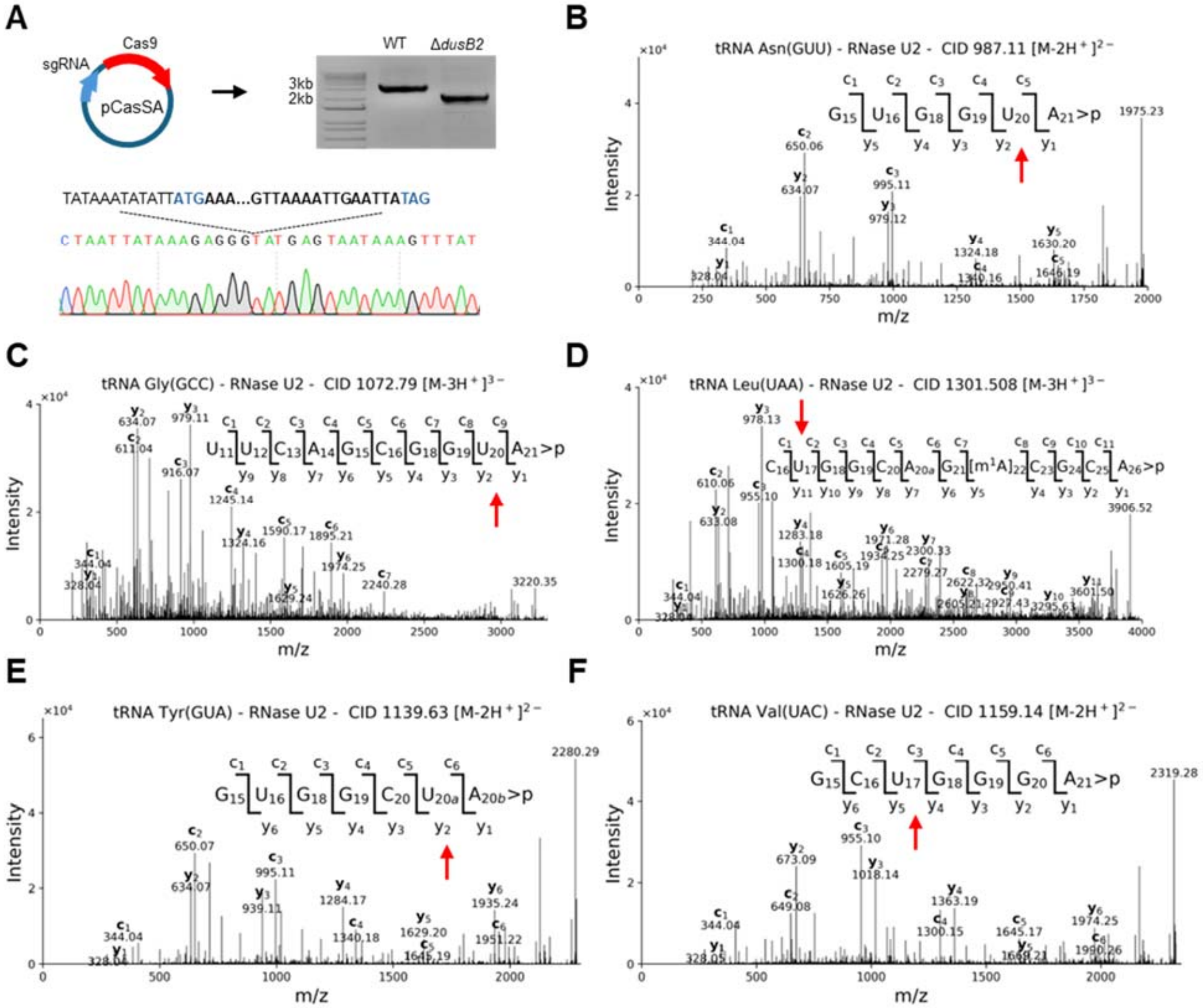

**Figure S7. DusB2 is responsible for [D]17, [D]20, and [D]20a synthesis in *S. aureus* tRNA.** (**A**) Deletion of *dusB2* using pCasSA CRISPR-Cas9. Agarose gel electrophoresis and Sanger sequencing chromatogram confirmed deletion of *dusB2* gene (WKU35_RS00150) in the *S. aureus* RN4220 strain. Start and stop codons are colored in blue. Similar results were obtained for deletion of HG001_00036 gene in HG001 strain. (**B**) Deconvoluted CID MS/MS spectra for tRNA Asn(GUU), (**C**) Gly(GCC), (**D**) Leu(UAA), (**E**) Tyr(GUA), (**F**) Val(UAC), showing the absence of D in the D-loop. Positions modified by DusB2 in wild-type strains are indicated with red arrows.

**A** Spot IV-09: NP1+NP2+P1

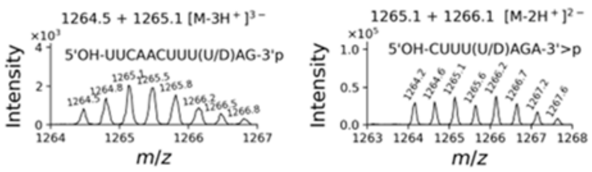


**B** Spot IV-13: NP3

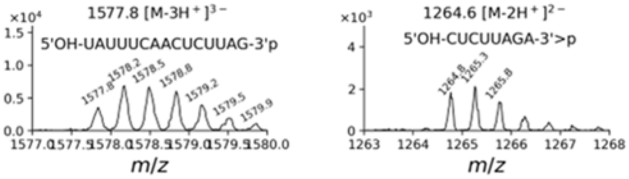


**C**

| Type | Bacteria | Acc-stem | | D-stem | D-loop | D-stem | | Ac-stem | Ac-loop | Ac-stem | V-region | T-stem | T-loop | T-stem | Acc-stem | | CCA |
|---|---|---|---|---|---|---|---|---|---|---|---|---|---|---|---|---|---|
| | | 1 | 8 | 10 | 14 | 22 | 26 | 27 | 32 | 39 | 44 | 49 | 54 | 61 | 66 | 73 | 74 |
| NP1 | *Sau* HG001 | GCGGGAG | [s$^4$U]A | GUUC | AACUUUUA | GAAC | A | CGUUC | CUUCCCG | GAACG | AGGU | AUAGG | UGCAAAU | CCUAU | CUUCCGC | U | CCA |
| | *Sau* N315 | GCGGGAG | UA | GUUC | AACUUUUA | GAAC | A | CGUUC | CUUCCCG | GAACG | AGGU | AUAGG | UGCAAAU | CCUAU | CUUCCGC | U | CCA |
| | *Sau* Newman | GCGGGAG | UA | GUUC | AACUUUUA | GAAC | A | CGUUC | CUUCCCG | GAACG | AGGU | AUAGG | UGCAAAU | CCUAU | CUUCCGC | U | CCA |
| | *Sha* JCSC1435 | GCGGGAG | UA | GUUC | AAUUUUA | GAAC | A | CAUUC | CUUCCCG | GAAUG | AGGU | AUAGG | UGCAAGU | CCUAU | CUUCCGC | U | CCA |
| | *Sca* TM300 | | | | | | | | | | | | | | | | |
| | *Ssa* ATCC 15305 | | | | | | | | | | | | | | | | |
| | *Shy* ATCC 11249 | | | | | | | | | | | | | | | | |
| | *Sep* Texas 26 | GCGGGAG | [s$^4$U]A | GUUC | AAUUU[D]A | GAAC | A | CAUUC | CUUCCCG | GAAUG | AGGU | AUAGG | UGCAAGU | CCUAU | CUUCCGC | U | CCA |
| | *Sep* RP62A | GCGGGAG | UA | GUUC | AACUCUCA | GAAC | A | CAUUC | CUUCCCG | GAAUG | AGAU | AUAGG | UGCAAAU | CCUAU | CUUCCGC | U | CCA |
| | *Sep* PM221 | GCGGGAG | UA | GUUC | AACUCUCA | GAAC | A | CAUUC | CUUCCCG | GAAUG | AGAU | AUAGG | UGCAAAU | CCUAU | CUUCCGC | U | CCA |
| | *Sep* SEI | GCGGGAG | UA | GUUC | AACUCUCA | GAAC | A | CAUUC | CUUCCCG | GAAUG | AGAU | AUAGG | UGCAAAU | CCUAU | CUUCCGC | U | CCA |
| | *Sep* ATCC 12228 | GCGGGAG | UA | GUUC | AACUCUCA | GAAC | A | CAUUC | CUUCCCG | GAAUG | AGAU | AUAGG | UGCAAAU | CCUAU | CUUCCGC | U | CCA |
| NP2 | *Sau* HG001 | GCGGGAG | [s$^4$U]A | GUUC | AACUUU[D]A | GAAC | A | CGUUC | CUUCCCG | GAACG | AGGU | AUAGG | UGUAAAU | CCUAU | CUUCCGC | U | CCA |
| | *Sau* N315 | GCGGGAG | UA | GUUC | AACUUUUA | GAAC | A | CGUUC | CUUCCCG | GAACG | AGGU | AUAGG | UGUAAAU | CCUAU | CUUCCGC | U | CCA |
| | *Sau* Newman | GCGGGAG | UA | GUUC | AACUUUUA | GAAC | A | CGUUC | CUUCCCG | GAACG | AGGU | AUAGG | UGUAAAU | CCUAU | CUUCCGC | U | CCA |
| | *Sha* JCSC1435 | | | | | | | | | | | | | | | | |
| | *Sca* TM300 | GCGGGAG | UA | GUUC | AACUCUUA | GAAC | A | CGUUC | CUUCCCG | GAACG | AGGU | AUAGG | UGUAAAU | CCUAU | CUUCCGC | U | CCA |
| | | GCGGGAG | UA | GUUC | AACUCUUA | GAAC | A | CGUUC | CUUCCCG | GAACG | AGGU | AUAGG | UGUAAAU | CCUAU | UUUCCGC | U | CCA |
| | *Ssa* ATCC 15305 | GCGGGGG | UA | GUUC | AACUCUUA | GAAC | A | CGUUC | CUUCCCG | GAACG | AGGU | GUAGG | UGUAAGU | CCUAU | CCUCCGC | U | CCA |
| | | GCGGGAG | UA | GUUC | AACUUUUA | GAAC | A | CAUUC | CUUCCCG | GAAUG | AGGU | AUAGG | UGUAAGU | CCUAU | CUUCCGC | U | CCA |
| | | GCGGGAG | UA | GUUC | AACUUUUA | GAAC | A | CGUUC | CUUCCCG | GAAUG | AGGU | AUAGG | UGUAAGU | CCUAU | CUUCCGC | U | CCA |
| | *Shy* ATCC 11249 | GCGGGAG | UA | GUUC | AACUCUCA | GAAC | A | CUUUC | CUUCCCG | GAAAG | AGGU | ACAGG | UGUAAAU | CCUGU | CUUCCGC | U | CCA |
| | | GCGGGAG | UA | GUUC | AACUCUCA | GAAC | A | CUUUC | CUUCCCG | GAAAG | AGGU | AUAGG | UGUAAAU | CCUAU | CUUCCGC | U | CCA |
| | *Sep* Texas 26 | | | | | | | | | | | | | | | | |
| | *Sep* RP62A | GCGGGAG | UA | GUUC | AACUCUUA | GAAC | A | CAUUC | CUUCCCG | GAAUG | AGAU | AUAGG | UGUAAAU | CCUAU | CUUCCGC | U | CCA |
| | | GCGGGAG | UA | GUUC | AACUCUCA | GAAC | A | CAUUC | CUUCCCG | GAAUG | AGAU | AUAGG | UGUAAAU | CCUAU | CUUCCGC | U | CCA |
| | *Sep* PM221 | GCGGGAG | UA | GUUC | AACUCUUA | GAAC | A | CAUUC | CUUCCCG | GAAUG | AGAU | AUAGG | UGUAAAU | CCUAU | CUUCCGC | U | CCA |
| | | GCGGGAG | UA | GUUC | AACUCUCA | GAAC | A | CAUUC | CUUCCCG | GAAUG | AGAU | AUAGG | UGUAAAU | CCUAU | CUUCCGC | U | CCA |
| | *Sep* SEI | GCGGGAG | UA | GUUC | AACUCUUA | GAAC | A | CAUUC | CUUCCCG | GAAUG | AGAU | AUAGG | UGUAAAU | CCUAU | CUUCCGC | U | CCA |
| | | GCGGGAG | UA | GUUC | AACUCUCA | GAAC | A | CAUUC | CUUCCCG | GAAUG | AGAU | AUAGG | UGUAAAU | CCUAU | CUUCCGC | U | CCA |
| | *Sep* ATCC 12228 | GCGGGAG | UA | GUUC | AACUCUUA | GAAC | A | CAUUC | CUUCCCG | GAAUG | AGAU | AUAGG | UGUAAAU | CCUAU | CUUCCGC | U | CCA |
| | | GCGGGAG | UA | GUUC | AACUCUCA | GAAC | A | CAUUC | CUUCCCG | GAAUG | AGAU | AUAGG | UGUAAAU | CCUAU | CUUCCGC | U | CCA |
| NEW | *Sau* HG001 | GCGGGAG | [s$^4$U]A | UUUC | AACUCU[D]A | GAAU | A | CAUUC | CUUCCCG | GAAUG | AGGU | AUAGG | UGUAAAU | CCUAU | CUUCCGC | U | CCA |
| | *Sau* N315 | GCGGGAG | UA | UUUC | AACUCUUA | GAAU | A | CAUUC | CUUCCCG | GAAUG | AGGU | AUAGG | UGUAAAU | CCUAU | CUUCCGC | U | CCA |
| | *Sau* Newman | GCGGGAG | UA | UUUC | AACUCUUA | GAAU | A | CAUUC | CUUCCCG | GAAUG | AGGU | AUAGG | UGUAAAU | CCUAU | CUUCCGC | U | CCA |
| | *Sha* JCSC1435 | GCGGGAG | UA | UUUC | AACUGUUA | GAAU | A | UGUUC | CUUCCCG | GAACA | AGGU | AUAGG | UGUAAAU | CCUAU | CUUCCGC | U | CCA |
| | | GCGGGAG | UA | UUUC | AACUUUUA | GAAU | A | CGUUC | CUUCCCG | GAACG | AGAU | AUAGG | UGCAAAU | CCUAU | CUUCCGC | U | CCA |
| | *Sca* TM300 | GCGGGAG | UA | UUUC | AACUUUCA | GAAU | U | CAUUC | CUUCCCG | GAAUG | AGGU | AUAGG | UGCAAAU | CCUAU | CUUCCGC | U | CCA |
| | *Ssa* ATCC 15305 | | | | | | | | | | | | | | | | |
| | *Shy* ATCC 11249 | | | | | | | | | | | | | | | | |
| | *Sep* Texas 26 | GCGGGAG | [s$^4$U]A | UUUC | AACUUU[D]A | GAAU | A | CGUUC | CUUCCCG | GAACG | AGAU | AUAGG | UGCAAAU | CCUAU | CUUCCGC | U | CCA |
| | *Sep* RP62A | | | | | | | | | | | | | | | | |
| | *Sep* PM221 | | | | | | | | | | | | | | | | |
| | *Sep* SEI | | | | | | | | | | | | | | | | |
| | *Sep* ATCC 12228 | | | | | | | | | | | | | | | | |

**Figure S8. Conservation and potential dihydrouridylation of non-proteogenic tRNAs Gly(UCC) species in *Staphylococci.*** (**A**) ESI-MS spectra of D-loop fragments derived from spot IV-09, containing both *S. aureus* HG001 NP1 and NP2, and (**B**) from spot IV-13, containing only NP3. The fragments were generated by digestion with RNase T1 (left) or U2 (right). In the NP1+NP2 sample, both spectra indicate a mixture of ions corresponding to unmodified and dihydrouridylated fragments. In NP3, the spectra are consistent with unmodified fragments. (**C**) Sequence alignment of non-proteogenic tRNA Gly(UCC) species from various *Staphylococci* species and strains. Sequences are grouped according to their similarity to one of the three *S. aureus* variants: NP1-type (G10-C25 base pair and U54-G55-C56 sequence), NP2-type (G10-C25 base pair and U54-G55-U56 sequence), and NP3 (unpaired U10 and U25). These characteristic features are highlighted in red. Anticodon nucleotides are underlined, and C32 and G38, shown in bold, are strictly conserved and likely form an extra base pair. EF-Tu recognition determinants (base pairs 49-65, 50-64, and 51-63) are also indicated in bold. Positions 8 and 20, potentially modified with $s^4U$ and D, respectively, are highlighted in blue. Experimentally verified [$s^4U$]8 and [D]20 modifications are indicated for *S. aureus* HG001 and *S. epidermidis* Texas 26 strains (3–5). Empty rows indicate the absence of the corresponding type, which in most cases reflects the presence of multiple sequences belonging to the same group. *Sau: Staphylococcus aureus*, *Sha*: *Staphylococcus haemolyticus*, *Sca*: *Staphylococcus carnosus*, *Ssa*: *Staphylococcus saprophyticus*, *Shy*: *Staphylococcus hyicus*, *Sep*: *Staphylococcus epidermidis*.

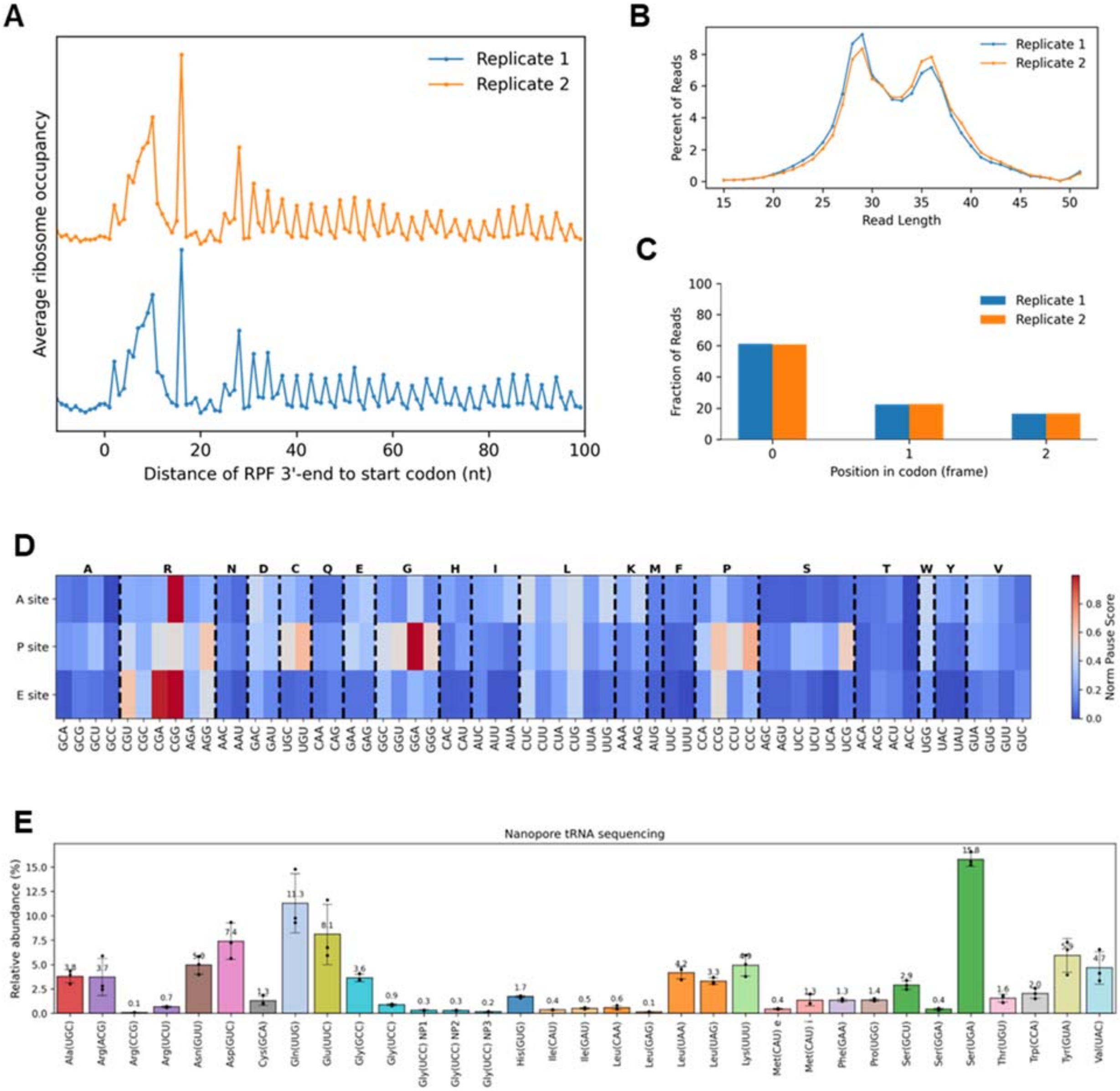
A
Replicate 1
Replicate 2
Average ribosome occupancy
Distance of RPF 3'-end to start codon (nt)
B
Percent of Reads
Read Length
C
Fraction of Reads
Position in codon (frame)
D
A site
P site
E site
Norm Pause Score
A R N D C Q E G H I L K M F P S T W Y V
GCA GCG GCU GCC CGU CGC CGA CGG AGA AGG AAC AAU GAC GAU UGC UGU CAA CAG GAA GAG GGC GGU GGA GGG CAC CAU AUC AUU AUA CUC CUU CUA CUG UUA UUG AAA AAG AUG UUC UUU CCA CCG CCU CCC AGC AGU UCC UCU UCA UCG ACA ACG ACU ACC UGG UAC UAU GUA GUG GUU GUC
E
Nanopore tRNA sequencing
Relative abundance (%)
Ala(UGC) Arg(ACG) Arg(CCG) Arg(UCU) Asn(GUU) Asp(GUC) Cys(GCA) Gln(UUG) Glu(UUC) Gly(GCC) Gly(UCC) Gly(UCC) NP1 Gly(UCC) NP2 Gly(UCC) NP3 His(GUG) Ile(CAU) Ile(GAU) Leu(CAA) Leu(GAG) Leu(UAA) Leu(UAG) Lys(UUU) Met(CAU) e Met(CAU) i Phe(GAA) Pro(UGG) Ser(GCU) Ser(GGA) Ser(UGA) Thr(UGU) Trp(CCA) Tyr(GUA) Val(UAC)

**Figure S9. Analysis of ribosome pausing and tRNA abundance in *S. aureus*.** (**A**) Metagene analysis of ribosome-protected fragments (RPFs) showing the characteristic 3-nucleotide periodicity of translating ribosomes in the two biological replicates. The x-axis represents positions relative to the start codon, and the y-axis indicates the fractions of RPF 3' ends mapping to each position. Peaks recurring every three nucleotides reflect codon-wise ribosome progression along mRNAs. (**B**) Size-distribution of RPFs mapping to the coding sequences. (**C**) Frame analysis of RPFs showing the distribution of reads among the three possible reading frames (0, 1, 2) based on their 3' end positions. A significant fraction of reads map to a preferred frame. (**D**) Ribosome pausing at A, P, and E-sites. Heatmap showing pausing score of codons at the ribosome A, P, and E sites. Amino acids are indicated above each group of synonymous codons. Pause values were independently normalized for each site to range from 0 to 1 for color mapping. Higher scores indicate increased ribosome occupancy, reflecting slower translation rates. Experiments were performed without chloramphenicol treatment and harvesting by filtration, and therefore no strong artifactual pauses were observed at Ser and Gly codons in A and E sites (6). (**E**) Relative abundance of tRNA species in *S. aureus*. Nanopore tRNA sequencing reads were used to estimate the relative abundance of the different tRNA isoacceptors in three biological replicates. Bar represent the average relative abundance of each tRNA and their corresponding values are labeled. Individual points for each replicate are shown. Error bars are based on standard deviation.

#### REFERENCES


1. Houser,W.M., Butterer,A., Addepalli,B. and Limbach,P.A. (2015) Combining recombinant ribonuclease U2 and protein phosphatase for RNA modification mapping by liquid chromatography–mass spectrometry. *Anal. Biochem.*, **478**, 52–58.

2. Mercier,N., Prévost,K., Massé,E., Romby,P., Caldelari,I. and Lalaouna,D. (2021) MS2-Affinity Purification Coupled with RNA Sequencing in Gram-Positive Bacteria. *J. Vis. Exp. JoVE*, 10.3791/61731.

3. Roberts,R.J. (1972) Structures of Two Glycyl-tRNAs from Staphylococcus epidermidis. *Nature. New Biol.*, **237**, 44–45.

4. Roberts,R.J., Lovinger,G.G., Tamura,T. and Strominger,J.L. (1974) Staphylococcal Transfer Ribonucleic Acids: I. ISOLATION AND PURIFICATION OF THE ISOACCEPTING GLYCINE TRANSFER RIBONUCLEIC ACIDS FROM STAPHYLOCOCCUS EPIDERMIDIS TEXAS 26. *J. Biol. Chem.*, **249**, 4781–4786.

5. Stewart,T.S., Roberts,R.J. and Strominger,J.L. (1971) Novel Species of tRNA. *Nature*, **230**, 36–38.

6. Mohammad,F., Green,R. and Buskirk,A.R. (2019) A systematically-revised ribosome profiling method for bacteria reveals pauses at single-codon resolution. *eLife*, **8**, e42591.
